# Decapping factor Dcp2 controls mRNA abundance and translation to adjust metabolism and filamentation to nutrient availability

**DOI:** 10.1101/2023.01.05.522830

**Authors:** Anil Kumar Vijjamarri, Xiao Niu, Matthew D. Vandermeulen, Chisom Onu, Fan Zhang, Hongfang Qiu, Neha Gupta, Swati Gaikwad, Miriam L. Greenberg, Paul J. Cullen, Zhenguo Lin, Alan G. Hinnebusch

## Abstract

Degradation of most yeast mRNAs involves decapping by Dcp1/Dcp2. DEAD-box protein Dhh1 has been implicated as an activator of decapping, in coupling codon non-optimality to enhanced degradation, and as a translational repressor, but its functions in cells are incompletely understood. RNA-Seq analyses coupled with CAGE sequencing of all capped mRNAs revealed increased abundance of hundreds of mRNAs in *dcp2*Δ cells that appears to result directly from impaired decapping rather than elevated transcription, which was confirmed by ChIP-Seq analysis of RNA Polymerase II occupancies genome-wide. Interestingly, only a subset of mRNAs requires Dhh1 for targeting by Dcp2, and also generally requires the other decapping activators Pat1, Lsm2, Edc3 or Scd6; whereas most of the remaining transcripts utilize NMD factors for Dcp2-mediated turnover. Neither inefficient translation initiation nor stalled elongation appears to be a major driver of Dhh1-enhanced mRNA degradation. Surprisingly, ribosome profiling revealed that *dcp2*Δ confers widespread changes in relative TEs that generally favor well-translated mRNAs. Because ribosome biogenesis is reduced while capped mRNA abundance is increased by *dcp2*Δ, we propose that an increased ratio of mRNA to ribosomes increases competition among mRNAs for limiting ribosomes to favor efficiently translated mRNAs in *dcp2*Δ cells. Interestingly, genes involved in respiration or utilization of alternative carbon or nitrogen sources are derepressed, and both mitochondrial function and cell filamentation (a strategy for nutrient foraging) are elevated by *dcp2*Δ, suggesting that mRNA decapping sculpts gene expression post-transcriptionally to fine-tune metabolic pathways and morphological transitions according to nutrient availability.

## INTRODUCTION

The translation and degradation of mRNA are closely related, as the post-transcriptionally added appendages, the 5’ m^7^G cap and 3’ poly(A) tail are involved in both mechanisms. In general, translation is initiated by association of initiation factor eIF4F (comprised of cap-binding protein eIF4E, scaffolding protein eIF4G, and helicase eIF4A) with the capped 5’ end of mRNA followed by recruitment of the small (40S) subunit of the ribosome pre-loaded with various other initiation factors and initiator methionyl tRNA. Interaction between eIF4G and poly(A)-binding protein (PABP) stabilizes a “closed-loop” mRNP conformation that serves both to protect the mRNA from degradation at both ends and enhance translation (Ghosh and Jacobson 2010) (Hinnebusch 2014). In yeast, degradation of mRNAs is generally initiated by shortening of the poly(A) tail (deadenylation), catalyzed by the Pan2/Pan3 and Ccr4-Not deadenylase complexes. In a minor pathway, deadenylation is followed by 3’ to 5’ exonucleolytic degradation by the cytoplasmic exosome complex. More commonly, deadenylated mRNAs are decapped by the conserved Dcp1/Dcp2 holoenzyme, which hydrolyzes the cap to release m^7^GDP and 5’ monophosphate RNA, which is then degraded 5’ to 3’ by the exoribonuclease Xrn1 (Parker 2012). Dcp2-mediated decapping is critical for multiple mRNA decay pathways including bulk 5’ to 3’ decay (Decker and Parker 1993), nonsense-mediated RNA decay (NMD) triggered by premature termination codons (He and Jacobson 2001), AU-rich mRNA decay (Barreau et al. 2005), microRNA-mediated turnover (Jonas and Izaurralde 2015) and transcript-specific degradation (Badis et al. 2004).

Dcp2 is a bi-lobed enzyme, consisting of an N-terminal regulatory domain (NRD) followed by a Nudix superfamily hydrolase domain, attached to an intrinsically disordered C-terminal region (IDR) that contains short leucine-rich helical motifs (HLMs). The N-terminal and C-terminal domains are attached via short, flexible linkers that allow Dcp2 to adopt multiple conformations in solution (Floor et al. 2012). Dcp1 enhances Dcp2 catalytic activity by interacting with the Dcp2 NRD and is essential for mRNA decapping in vivo (Beelman et al. 1996; Steiger et al. 2003). Although Dcp1 promotes the closed conformation of Dcp2, the active site is not fully formed and the RNA binding site is blocked by the NRD, thus forming a catalytically incompetent enzyme complex (Wurm et al. 2017). Additional factors stimulate the catalytic activity of the Dcp1/Dcp2 complex, including enhancer of decapping (Edc) proteins. Interaction of Dcp1 with a short proline-rich motif present in Edc1-type activators, including yeast Edc2 (Borja et al. 2011) and mammalian PNRC (Lai et al. 2012), stabilizes the active conformation of the Dcp1/Dcp2 complex in the presence of substrate (Wurm et al. 2017). Autoinhibitory motifs located in the IDR interact with the core domain of Dcp2 and stabilize the closed, catalytically inactive conformation of Dcp2 (He and Jacobson 2015; Paquette et al. 2018). Edc1 alone cannot overcome the inhibitory effect of the IDR, requiring additional stimulation by Edc3 (Paquette et al. 2018). The LSm domain of Edc3 binds to HLMs located in the IDR and activates decapping by alleviating autoinhibition and promoting RNA binding by Dcp2 in yeast. Moreover, deletion of the autoinhibitory region in Dcp2 bypasses activation by Edc3 (Fromm et al. 2012; He and Jacobson 2015).

In addition to Edc proteins, Dhh1, Pat1, Scd6 and Lsm1-7 function in general mRNA decay by activating Dcp1/Dcp2 (Parker 2012). It appears that Edc3 and Scd6 may act interchangeably in recruitment of Dhh1 to the same segment of the Dcp2 IDR (He et al. 2022). The HLMs in Dcp2 additionally mediate interaction with Pat1 (Charenton et al. 2017); although the Pat1/Lsm1-7 complex can also bind to the 3’ ends of deadenylated mRNAs to stimulate decapping by Dcp1/Dcp2 (Tharun et al. 2000; Tharun and Parker 2001). Numerous direct interactions were identified among the decapping activators, as Dhh1 interacts with Pat1, Scd6 and Edc3, and Pat1 interacts with Scd6 in addition to Lsm1-7, as well as with Xrn1 (Nissan et al. 2010; Sharif et al. 2013; He and Jacobson 2015). Interestingly Dhh1 and the Pat1/Lsm1-7 complex appear to target distinct subsets of mRNAs with overlapping substrate specificities (He et al. 2018). From these results and others (He et al. 2022), it can be proposed that the decapping activators assemble into distinct decapping complexes that target different subsets of mRNAs for degradation. Dhh1, Pat1, Scd6 and Lsm1-7, along with Dcp1/Dcp2, are found concentrated in processing bodies (PBs) with translationally repressed mRNAs (Parker and Sheth 2007), consistent with a role in translational repression in addition to mRNA turnover. Recently, Edc3 and Scd6 were shown to retain Dcp2 in the cytoplasm, preventing import of Dcp1/Dcp2 into the nucleus where it cannot function in cytoplasmic mRNA decay (Tishinov and Spang 2021). Thus, decapping activators can stimulate decapping by multiple, distinct mechanisms.

Dhh1 is a highly conserved ATP-dependent DEAD-box helicase involved in translation repression and mRNA decay in yeast. Dhh1 and its orthologues in *Schizosaccharomyces pombe* (Ste13), *Caenorhabditis elegans* (CGH-1), *Xenopus laevis* (Xp54), *Drosophila melanogaster* (Me31b), and mammals (RCK/p54) interact with factors involved in deadenylation, decapping and translational repression (Coller et al. 2001; Fischer and Weis 2002; Maillet and Collart 2002; Weston and Sommerville 2006). Dhh1 promotes decapping-mediated mRNA turnover, as deadenylated capped mRNAs are stabilized in *dhh1*Δ cells. Moreover, recombinant Dhh1 stimulates the decapping activity of the purified decapping enzyme (Coller et al. 2001; Fischer and Weis 2002). The mRNAs preferentially targeted by both Dhh1 and Pat1/Lsm1-7 appear to be inefficiently translated at the elongation stage (He et al. 2018).

Although most yeast mRNAs are targeted by a common degradation pathway, the half-lives of individual mRNAs vary greatly (Coller and Parker 2004). Specific sequence and structural elements present in 5’ or 3’ UTRs, and poly(A) tail lengths, can modulate the degradation rate; however, these features alone do not appear to explain the variation in half-lives among all transcripts (Muhlrad and Parker 1992; Lee et al. 2013; Geisberg et al. 2014). Recently, codon-optimality—the balance between the supply of charged tRNA molecules in the cytoplasmic pool and the demand of tRNA usage by translating ribosomes—was identified as a determinant of mRNA decay. Evidence indicates that non-optimal codons, which are decoded by ribosomes more slowly than optimal codons, accelerate mRNA decay co-translationally (Hu et al. 2009; Presnyak et al. 2015). Consistent with this, Caf1, a subunit of the Ccr4-Not deadenylase complex appears to selectively target transcripts with slower elongation rates and reduced association with Pab1 (Webster et al. 2018). Multiple lines of evidence support the model that Dhh1 binds to ribosomes slowly elongating through non-optimum codons and triggers their decapping and degradation (Radhakrishnan et al. 2016). Recently, cryo-EM analysis revealed that the N-terminal region of Not5 interacts with the E-site of ribosomes stalled at sub-optimal codons with an empty A-site, leading to the model that ribosomes stalled at sub-optimal codons are detected by Not5/Caf1 and targeted for degradation via Dhh1 (Buschauer et al. 2020). Much of the evidence indicating that codon non-optimality is a major determinant of mRNA instability derives from reporter transcripts with relatively long strings of non-optimal or optimal codons, which likely does not pertain to many native mRNAs.

A different model proposes that translation initiation and decay are inversely related, with functional initiation complexes impeding both decapping and deadenylation. Thus, mutations impairing the cap-binding eIF4F complex or initiation factor eIF3 that reduce translation initiation were found to accelerate degradation of particular yeast mRNAs via elevated deadenylation and decapping (Schwartz and Parker 1999). In addition, strong secondary structures in the 5’ UTR (Muhlrad et al. 1995) and poor start codon context (LaGrandeur and Parker 1999) both increased decapping. Moreover, binding of eIF4E to the cap structure was sufficient to inhibit decapping by Dcp1/Dcp2 (Schwartz and Parker 2000). From this model it can be predicted that translation initiation, directly or indirectly, competes with the decay machinery in a manner that influences mRNA turnover rates. Indeed, non-invasive measurements of mRNA half-lives suggested that competition between translation initiation and mRNA decay factors is a major determinant of yeast mRNA turnover, whereas global inhibition of elongation generally led to stabilization of transcripts (Chan et al. 2018).

Starving yeast for glucose engenders a rapid loss of protein synthesis, accompanied by a shift of mRNAs from polysomes to free mRNPs and an increase in both size and number of PBs (Ashe et al. 2000; Teixeira et al. 2005; Arribere et al. 2011). Deletion of *DHH1* or *PAT1* partially impaired the loss of polysomes evoked by glucose depletion, whereas deletion of both genes simultaneously abrogated translational repression, suggesting that either Dhh1 or Pat1 is sufficient for repression of translation initiation in glucose starvation (Holmes et al. 2004; Coller and Parker 2005). Furthermore, overexpression of Dhh1 or Pat1 in WT cells conferred a general repression of translation with attendant PB formation (Coller and Parker 2005). These studies suggest that Dhh1 and Pat1 act as general repressors of translation in glucose-starved yeast. Consistent with this, Dhh1 can inhibit translation initiation in vitro by blocking assembly of 48S preinitiation complexes, and can impede elongation by associating with slowly-moving ribosomes (Coller and Parker 2005; Sweet et al. 2012). Moreover, the Not4 subunit of Ccr4-Not complex, as well as Dhh1 and Dcp1, were implicated in translational repression of mRNAs that exhibit transient ribosome stalling in a manner that limits protein misfolding during nutrient limitation (Preissler et al. 2015). Together, these findings suggest that multiple factors that stimulate mRNA decay can also repress translation at the initiation or elongation steps during nutrient starvation. Interestingly, Dhh1 helicase activity was reported to enhance the translation of *ATG1* and *ATG13* mRNAs, encoding proteins required for autophagy, under nitrogen-starvation conditions (Liu et al. 2019). Ribosome profiling of *dhh1*Δ cells grown in rich medium uncovered hundreds of mRNAs displaying either increased or decreased relative translation efficiencies (TEs), suggesting that Dhh1 can function as a translational repressor or activator for distinct sets of mRNAs, even in nonstarvation conditions (Radhakrishnan et al. 2016; Jungfleisch et al. 2017; Zeidan et al. 2018).

Based on its central role in mRNA decapping, previous studies on Dcp2 have focused primarily on mRNA decay, and the participation of Dcp2 in regulating translational efficiencies has not been explored. Here, we demonstrate that in nutrient-replete cells Dcp2 modulates translational efficiencies in addition to mRNA turnover by pathways both dependent and independent of decapping activator Dhh1. Surprisingly, a large fraction of mRNAs appear to be targeted for degradation by Dcp2 independently of Dhh1, Pat1, Edc3, or Scd6, which are enriched for NMD substrates, whereas the remaining fraction is controlled concurrently by all four decapping activators. Codon non-optimality does not appear to be a major driver of mRNA degradation for either set of Dcp2-targeted mRNAs. Unexpectedly, ribosome profiling revealed that *dcp2*Δ confers widespread changes in relative TEs that generally favor mRNAs that are well-translated in WT cells at the expense of more poorly translated mRNAs. This competition can be attributed to increased mRNA abundance, resulting from impaired decapping and attendant 5’-3’ decay, coupled with reduced ribosome production elicited by the Environmental Stress Response (Gasch et al. 2000). Superimposed on this general translational reprogramming is a Dhh1-dependent translational repression of certain poorly translated mRNAs by Dcp2. Finally, we provide evidence that Dcp2 helps to repress the abundance or translation of many mRNAs encoding proteins whose functions are dispensable during growth on glucose-replete rich medium, involved in catabolism of non-preferred carbon or nitrogen sources, mitochondrial respiration, or cell filamentation and invasive growth. These last findings support the emerging model that regulators of mRNA turnover add a layer of post-transcriptional control to well-established transcriptional repression mechanisms that control these various responses to nutrient limitation.

## RESULTS

### Dcp2 controls mRNA abundance via both Dhh1-dependent and -independent pathways

To determine the role of Dcp2 and Dhh1 in regulating mRNA abundance and translation in nutrient-replete cells, we interrogated our previous ribosome profiling and RNA-Seq datasets obtained from isogenic WT, *dcp2*Δ, *dhh1*Δ and *dcp2*Δ*dhh1*Δ strains cultured in nutrient-rich YPD medium at 30°C (Zeidan et al. 2018). RNA-Seq analysis identified numerous mRNAs differentially expressed in *dcp2*Δ vs. WT cells, including 1376 derepressed mRNAs (Figure 1A, blue) exhibiting a median increase of 1.96-fold (Figure 1B, blue) (dubbed mRNA_up_*dcp2*Δ), and 1281 repressed transcripts (Figure 1A, red) showing a median decrease of 0.57-fold in mRNA abundance conferred by *dcp2*Δ (Figure 1B, red) (mRNA_dn_*dcp2*Δ) (Note in Figure 1B that the median change for all mRNAs is unity (log_2_ = 0) owing to normalization of RNA reads for library depth for both mutant and WT strains. In this and all subsequent box-plots, if the notches in two boxes do not overlap, their median values differ with 95% confidence). In an effort to validate the RNA-Seq results, we performed qRT-PCR analysis of selected up- or down-regulated transcripts in total mRNA from *dcp2*Δ and WT cells, normalizing their abundance to a luciferase mRNA spiked into all samples. We found a strong positive correlation between the mRNA changes identified in RNA-Seq or qRT-PCR (Spearman’s correlation coefficient ρ = 0.86) (Figure 1C and Figure 1 – figure supplement 1A).

**Figure 1.**
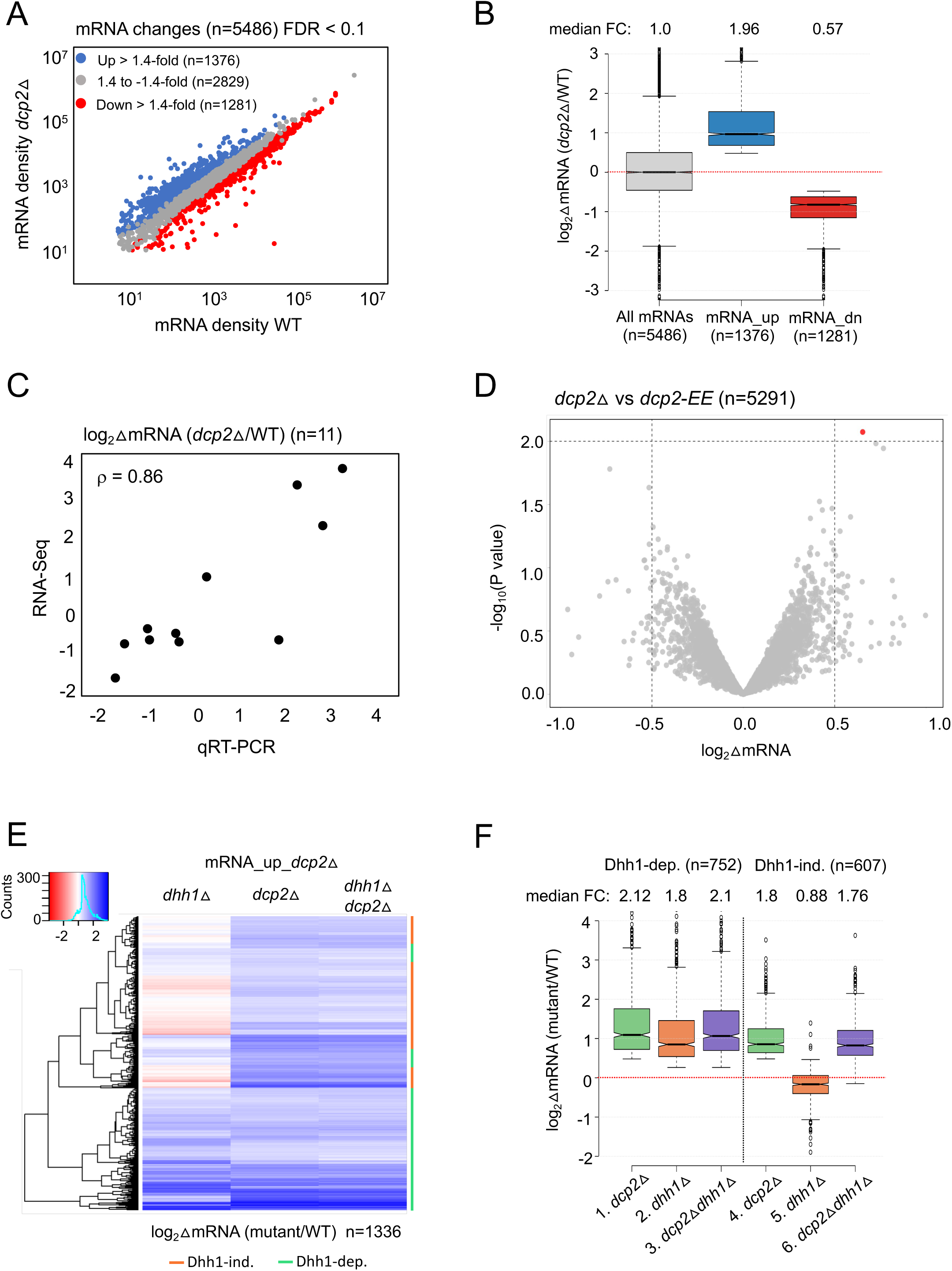
Dhh1-dependent and -independent regulation of mRNA abundance by decapping factor Dcp2. **(A)** Scatterplot of normalized mRNA densities in RPKM (Reads Per Kilobase of transcript, per Million mapped reads) for 5486 genes in *dcp2*Δ vs. WT cells determined by RNA-Seq. Genes showing significant changes in mRNA abundance of >1.4-fold in *dcp2*Δ vs. WT cells (at FDR < 0.1) as determined by DESeq2 analysis are shown as blue or red dots. **(B)** Notched box-plot of the log_2_ fold-changes in mRNA abundance (from RNA-Seq) in *dcp2*Δ vs. WT cells for all mRNAs (n=5486, median FC = 1) or subsets of transcripts either derepressed (n=1376, median FC = 1.96) or repressed (n=1281, median FC = 0.57). Median values for each group are indicated at the top, and the numbers of mRNAs for which data were obtained for each group is indicated at the bottom. A few outliers were omitted to expand the y-axis scale. **(C)** Scatterplot showing correlation between the log_2_ fold-changes in mRNA abundance determined by RNA-Seq vs. qRT-PCR for 11 different genes in *dcp2*Δ vs. WT cells, with the Spearman correlation coefficient (ρ) indicated. **(D)** Evidence that mRNA changes conferred by *dcp2*Δ result from loss of Dcp2 catalytic activity. A Volcano-plot showing log_2_ fold-changes in mRNA abundance (from RNA-Seq) in *dcp2*Δ vs. *dcp2*-*E149Q,E153Q* (*dcp2-EE*) (x-axis) vs. -log_10_P values for the mRNA changes (y-axis) cells for all mRNAs (n=5291). **(E)** Hierarchical clustering analysis of the log_2_ fold-changes in mRNA abundance conferred by *dhh1*Δ, *dcp2*Δ or *dhh1*Δ*dcp2*Δ vs. WT for 1336 of the mRNA_up_*dcp2*Δ transcripts that are derepressed in abundance in *dcp2*Δ vs. WT cells. Transcripts annotated on the right with green or red bars require Dhh1 or are independent of Dhh1, respectively, for their repression by Dcp2 in WT cells. The color scale indicating log_2_ΔmRNA values ranges from 4 (strong derepression, dark blue) to -4 (strong repression, dark red). A few outlier mRNAs with log_2_ΔmRNA values of > 4.0 or < -4.0 were excluded to enhance the color differences among the remaining mRNAs. **(F)** Notched box-plot of log_2_ΔmRNA values in the mutants indicated at the bottom vs. WT for the two sets of mRNAs repressed in abundance by Dcp2 in a manner dependent (cols. 1-3) or independent of Dhh1 (cols. 4-6). Note that 17 of the 1376 mRNA_up_*dcp2*Δ transcripts were not detected in the *dhh1*Δ vs. WT experiment and, hence, were excluded from consideration. A few outliers were omitted from the plots to expand the y-axis scale. Horizontal red dotted lines indicate log_2_ΔmRNA values of zero, for a median fold-change of 1.

To identify the importance of Dcp2 catalytic activity in the observed mRNA abundance changes conferred by *dcp2*Δ, we conducted RNA-Seq on an isogenic *dcp2*-*E149Q,E153Q* mutant (*dcp2-EE*) with Gln substitutions of two key conserved Glu residues in the catalytic domain expected to abolish decapping activity (Aglietti et al. 2013; He et al. 2018). The RNA-Seq results for the WT, *dcp2*Δ, and *dcp2-EE* strains showed high reproducibility among biological replicates (ρ = 0.99) (Figure 1 – figure supplement 2A-C), and the mRNA densities for all expressed genes were also highly correlated between the *dcp2*Δ and *dcp2-EE* strains (ρ = 0.99) (Figure 1 – figure supplement 2G), with all but one mRNAs showing no significant differences between *dcp2*Δ vs. *dcp2-EE* strains (Figure 1D). Consistent with these results, the mRNA changes conferred by the catalytically defective *dcp2-E153Q-N245* and *dcp2-E198Q-N245* alleles (encoding only the N-terminal 245 residues of Dcp2) were found previously to be highly correlated with those conferred by *dcp2*Δ (correlation coefficients of 0.7-0.8) (He et al. 2018). Together these results indicate that the decapping activity of Dcp2 mediates its effects on the majority of yeast mRNAs whose abundance is controlled by Dcp2 in vivo.

To evaluate the contribution of decapping activator Dhh1 in controlling mRNA abundance by Dcp2, we compared the mRNA expression changes conferred by *dcp2*Δ, *dhh1*Δ, and the *dcp2*Δ*dhh1*Δ double mutation for the aforementioned 1376 transcripts derepressed by the *dcp2*Δ single mutation (mRNA_up_*dcp2*Δ group). A subset of 752 transcripts (dubbed Dhh1-dependent) show similar derepression in all three mutants (Figure 1E, groups labeled with green bars), with median increases in abundance compared to WT cells ranging from 1.8-to 2.1-fold in the three mutants (Figure 1F, cols. 1-3). The fact that combining *dcp2*Δ and *dhh1*Δ confers similar derepression ratios as those given by each single mutation would be expected if Dhh1 and Dcp2 function in the same pathway to control the abundance of these mRNAs, with Dhh1 activating decapping by Dcp2 and attendant 5’ to 3’ degradation. In contrast, the remaining 607 transcripts in the mRNA_up_*dcp2*Δ group showed little change in abundance in the *dhh1*Δ single mutant, despite derepression in the *dcp2*Δ mutant *on par* with that observed for the Dhh1-dependent group (Figure 1E, groups labeled with red bars). Whereas *dcp2*Δ conferred a median derepression of 1.8-fold, *dhh1*Δ produced a small repression of 0.88-fold for these mRNAs compared to WT (Figure 1F, cols. 4-5). As would expected if these mRNAs are decapped independently of Dhh1, they display essentially the same derepression in response to the *dcp2*Δ and *dhh1*Δ*dcp2*Δ mutations vs. WT (Figure 1E, cols. 2-3; Figure 1F, cols. 4 & 6). Supporting our analysis above, we interrogated RNA-Seq data for *dhh1*Δ vs. WT comparisons from three published studies, two of which involved a different strain background from ours (Radhakrishnan et al. 2016; Jungfleisch et al. 2017) (He et al. 2018). Importantly, the transcripts derepressed by *dcp2*Δ in a Dhh1-dependent manner in our study, showed significant increases in median mRNA abundance in *dhh1*Δ vs. WT comparisons in all three datasets (Figure 1 – figure supplement 1B, cols. 5-8); whereas our Dhh1-independent group of transcripts showed either little change or reduced median abundance in the published datasets (Figure 1 – figure supplement 1B, cols. 9-12). Together, the results suggest that only about one-half of the mRNAs targeted by Dcp2 for degradation are dependent on Dhh1 in WT cells for activation of decapping.

### Multiple decapping activators target a common subset of transcripts for Dcp2-mediated repression of mRNA abundance

Apart from Dhh1, other decapping activators include enhancers of decapping factors (Edcs), Scd6, Pat1/Lsm1-7 complex and the Upf factors of the NMD pathway. To evaluate whether Dhh1-independent degradation by Dcp2 involves other decapping activators that function in place of Dhh1, we evaluated RNA-Seq data we obtained recently comparing isogenic *pat1*Δ, *pat1*Δ*dhh1*Δ or *edc3*Δ*scd6*Δ mutants to WT as well as published RNA-Seq data on an isogenic *upf1*Δ mutant lacking a key NMD factor (Celik et al. 2017). Interestingly, *pat1*Δ, *pat1*Δ*dhh1*Δ and *edc3*Δ*scd6*Δ mutants exhibit increased median abundance for the Dhh1-dependent mRNA_up_*dcp2*Δ group of transcripts compared to WT, as observed for the *dhh1*Δ mutant (Figure 2A, cols. 2-5). Consistent with this, cluster analysis of individual transcripts reveals increases of similar magnitude in the four decapping activator mutants and the *dcp2*Δ strain for many of the Dhh1-dependent transcripts (Figure 2B); although, there are also numerous transcripts derepressed to a greater or lesser extent in the *pat1*Δ, *pat1*Δ*dhh1*Δ or *edc3*Δ*scd6*Δ mutants compared to the *dhh1*Δ single mutant. (An in depth analysis of this transcript specificity is being presented elsewhere). In contrast, *upf1*Δ did not increase the median mRNA abundance of this group of transcripts (Figure 2A, col. 6) and cluster analysis revealed derepression by *upf1*Δ of only a small subset of the Dhh1-dependent trasnscripts elevated in the other mutants (Figure 2B). Thus, the mRNAs dependent on Dhh1 for repression of their abundance by Dcp2 generally also require Pat1 and/or Edc3/Scd6, but not Upf1, for full repression in WT cells.

**Figure 2.**
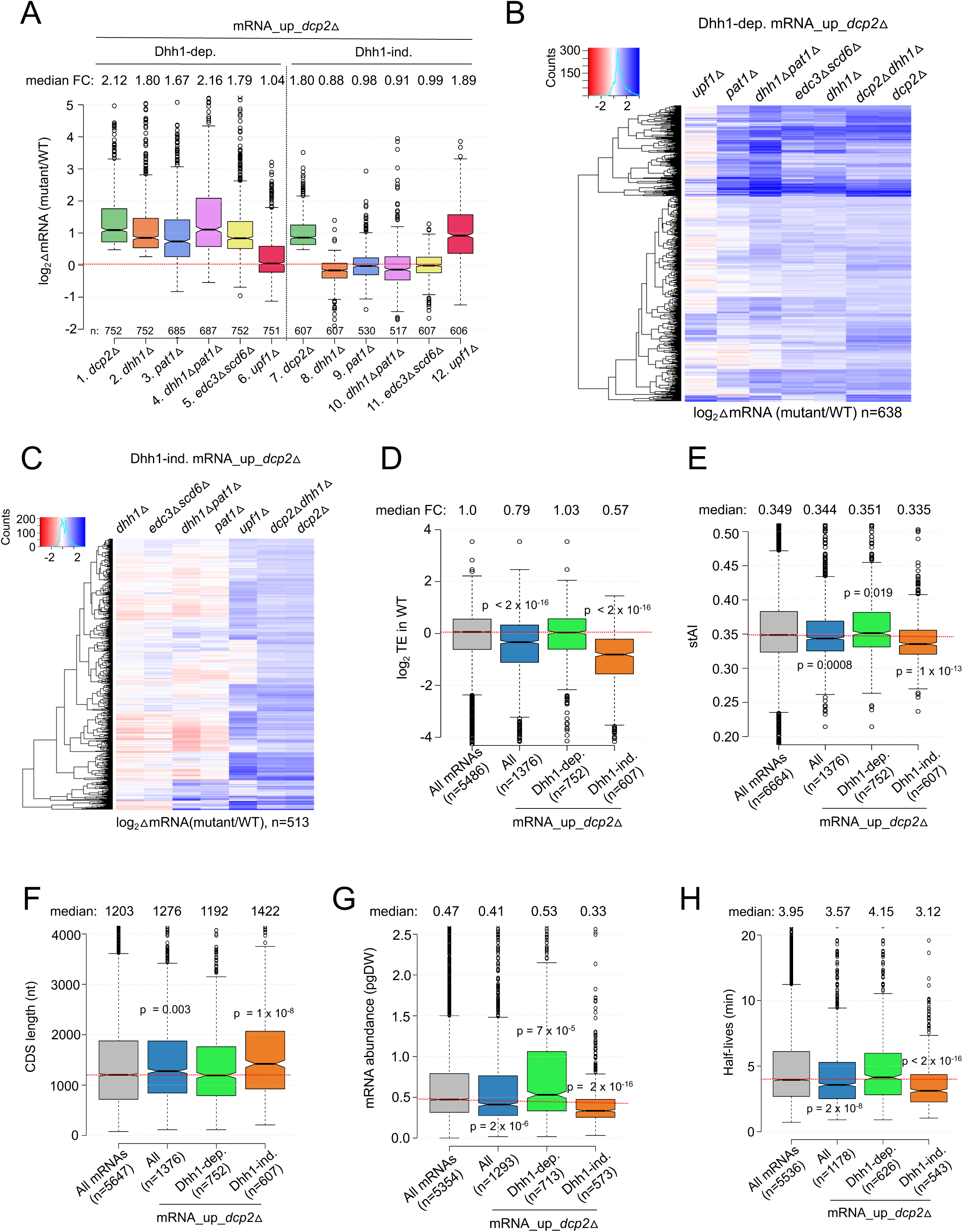
Multiple decapping activators function in unison to repress a subset of Dcp2-repressed mRNAs. **(A)** Notched box-plot as in Figgure 1B showing log_2_ fold-changes in mRNA abundance in the mutants indicated at the bottom vs. WT for the Dhh1-dependent (cols. 1-6) or Dhh1-independent (cols. 7-12) subsets of the mRNA_up_*dcp2*Δ set of transcripts. RNA-Seq data for the *pat1*Δ, *dhh1*Δ*pat1*Δ and *edc3*Δ*scd6*Δ mutants were from unpublished datasets deposited at GEO (GSE220578). **(B & C)** Hierarchical clustering analyses as in Figure 1E of the log_2_ fold-changes in mRNA abundance conferred by the mutations listed across the top vs. WT for the Dhh1-dependent (panel B, n=638) or Dhh1-independent (panel C, n=513) subset of the mRNA_up_*dcp2*Δ transcripts (excluding a few outliers with log_2_ΔmRNA values >4 or <-4). **(D-H)** Notched box-plot as in Figure 1B showing the log_2_TE in WT cells (D), stAI values (E), CDS length (F), mRNA abundance expressed as pg per dry cellular weight (G) and half-lives (H) for All mRNAs, all mRNA_up_*dcp2*Δ transcripts, or the Dhh1-dependent or -independent subsets of mRNA_up_*dcp2*Δ transcripts. P-values calculated using the Mann-Whitney U test for the differences between all mRNAs and the indicated groups are shown. For (D) WT TE values were calculated as the ratio of mRNA reads protected from RNAse digestion by association with translating 80S ribosomes (ribosome-protected fragments, RPFs) to the total mRNA reads measured by RNA-Seq for the same transcript in WT cells.

A markedly different outcome was observed for the Dhh1-independent group of mRNA_up_*dcp2*Δ transcripts, which showed no increase in median mRNA abundance in the *pat1*Δ, *pat1*Δ*dhh1*Δ, or *edc3*Δ*scd6*Δ mutants relative to WT, as observed for *dhh1*Δ (Figure 2A, cols. 8-11). Cluster analysis confirmed that most transcripts in this group showed little change in abundance in all four of these decapping activator mutants; although, a subset of exceptional mRNAs were derepressed in the single and double mutant lacking *PAT1* (Figure 2C). Thus, the majority of Dhh1-independent mRNAs also appear to be independent of Pat1, Scd6 or Edc3 for their repression by Dcp2. The fact that expression of these mRNAs is generally unaffected even in the *pat1*Δ*dhh1*Δ and *edc3*Δ*scd6*Δ double mutants eliminates the possibility that they exhibit redundant requirements for either Edc3 or Scd6, or either Pat1 or Dhh1. Interestingly, *upf1*Δ increased the median abundance of this group of transcripts comparably to *dcp2*Δ (Figure 2A, col. 12 vs. col. 7) and derepressed a much larger fraction than did any of the other decapping activator mutations (Figure 2C). All three mutants lacking NMD factors Upf1, Upf2, or Upf3 exhibited similar marked derepression of the Dhh1-independent transcripts with little effect on the Dhh1-dependent group (Figure 2 – figure supplement 1). These results are significant in revealing two groups of mRNAs whose abundance is repressed by Dcp2 in a manner that (i) requires the concerted action of Dhh1, Pat1 and Edc3/Scd6 (Dhh1-dependent group) or (ii) frequently requires the NMD factors but none of the other four decapping activators, nor even one of the Edc3/Scd6 and Dhh1/Pat1 pairs of activators, for repression by Dcp2 (Dhh1-independent group).

As noted above, previous findings have suggested that a key determinant of decapping and mRNA decay is the rate of translation at either the initiation or elongation stages (Chan et al. 2018; Hanson et al. 2018). In particular, Dhh1 has been implicated in targeting mRNAs enriched for slowly decoded non-optimal codons for decapping and decay (Radhakrishnan et al. 2016). Consistent with previous findings (He et al. 2018), we observed that the entire group of transcripts derepressed in abundance by *dcp2*Δ (mRNA_up_*dcp2*Δ) exhibit lower than average translational efficiencies (TEs) in WT cells, as determined by ribosome profiling of our *dcp2*Δ and WT strains conducted in parallel with RNA-Seq (Zeidan et al. 2018) (Figure 2D, col. 2). However, these mRNAs have only a slightly lower species-specific tRNA adaptation index (stAI) (Figure 2E, col. 2), a measure of codon optimality that quantifies the relative cellular supply of cognate and near-cognate tRNAs for a given codon (Radhakrishnan et al. 2016), suggesting that their lower than average TEs generally reflect slower rates of initiation versusvs. elongation. The Dhh1-dependent subset of Dcp2-repressed mRNAs exhibits average TEs and sTAI values (Figures 2D-E, col. 3), ostensibly at odds with the notion that an abundance of suboptimal codons is a key attribute of most mRNAs targeted by Dhh1 for enhanced decapping and decay (Radhakrishnan et al. 2016). It is possible, however, that these mRNAs often contain short runs of suboptimal codons that trigger Dhh1-dependent decapping despite an average frequency of such codons over the entire CDS.

Consistent with previous results on NMD substrates (Celik et al. 2017), the Dhh1-independent transcripts exhibit lower median TE and stAI values compared to all Dcp2-repressed mRNAs (Figures 2D-E, cols. 4 vs. 2). They also exhibit other features of poorly translated mRNAs (Pelechano et al. 2013; Radhakrishnan et al. 2016; Lahtvee et al. 2017; Chan et al. 2018), including a greater median length of coding sequences (CDS), lower median transcript abundance in WT cells, and lower median mRNA half-life, compared to all mRNAs and all Dcp2-repressed mRNAs, none of which are characteristic of the Dhh1-dependent group of transcripts (Figure 2F-H, cols. 3-4 vs.1-2). It is unclear whether any of these attributes of Dhh1-independent mRNAs are instrumental in enhancing decapping/degradation or targeting NMD factors to the Upf1-dependent members of the group. It has been suggested that low codon optimality may increase frameshifting errors that lead to premature termination events recognized by the Upf proteins (Celik et al. 2017).

### Both Dhh1-dependent and -independent repression of mRNA abundance by Dcp2 appears to involve decapping and degradation rather than reduced transcription

It was shown previously that *dcp2*Δ and many other slow-growing yeast mutants exhibit changes in gene expression (O’Duibhir et al. 2014) that correspond to the Environmental Stress Response (ESR), wherein hundreds of mRNAs are either repressed (rESR, 545 genes) or induced (iESR, 283 genes) stereotypically in response to diverse stress or starvation conditions (Gasch et al. 2000). Induction of iESR genes involves paralogous transcription factors, Msn2 and Msn4, that bind stress response elements (STREs) in promoters (Gorner et al. 1998); whereas most rESR gene products are involved in ribosome biogenesis or translation. Consistent with the previous findings, gene ontology (GO) analysis of the mRNAs dysregulated by *dcp2*Δ identified stress response genes for the mRNA_up_*dcp2*Δ group, and genes involved in ribosome production and translation for the mRNA_dn_*dcp2*Δ group of genes (Figure 7 – figure supplement 1A-B). Accordingly, it was possible that a large fraction of the mRNAs derepressed by *dcp2*Δ are induced at the transcriptional level as a manifestation of the iESR. At odds with this idea however, the Dhh1-independent mRNA_up_*dcp2*Δ transcripts are ∼2-fold underrepresented in iESR mRNAs (p=5.3 x 10^-3^) (Figure 3B); and although the Dhh1-dependent group is significantly overrepresented for iESR mRNAs (p=1.9 x 10^-90^) (Figure 3A), 76% of them are not iESR transcripts. Thus, the bulk of mRNA derepression conferred by *dcp2*Δ does not involve the ESR.

**Figure 3:**
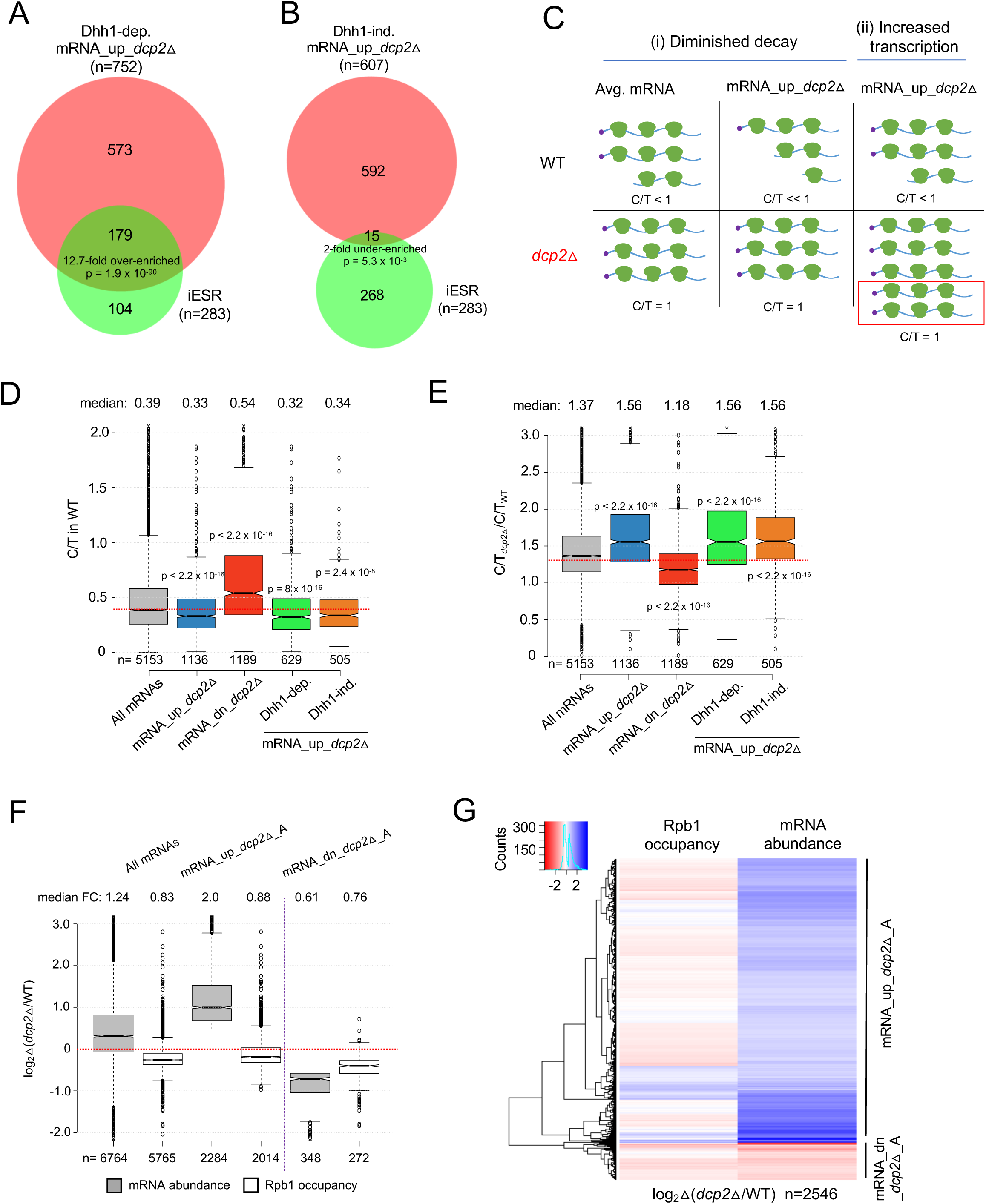
Evidence that Dhh1-independent changes in mRNA abundance conferred by *dcp2*Δ do not result from altered transcription. **(A-B)** The Dhh1-independent subset of Dcp2-repressed mRNAs are not enriched for iESR transcripts. Proportional Venn diagrams showing overlap between the Dhh1-dependent (A) and -independent (B) subsets of mRNA_up_*dcp2*Δ transcripts with induced ESR (iESR) mRNAs. Hypergeometric distribution p-values are displayed for significant changes. **(C)** Schematics depicting the predicted effects of *dcp2*Δ on levels of the capped proportion of mRNAs for the mRNAs preferentially repressed by Dcp2 compared to the average mRNA, according to the two different derepression mechanisms of (i) diminished decapping/degradation, and (ii) increased transcription. *Mechanism (i).* In WT cells (upper panel), mRNAs preferentially targeted by Dcp2 for decapping and degradation (mRNA_up_*dcp2*Δ) have a smaller proportion of capped transcripts (C/T << 1) compared to mRNAs with average susceptibility to Dcp2 (Avg. mRNA, C/T < 1). In *dcp2*Δ cells (lower panel, red), the C/T ratios for both groups of mRNAs increase to unity, which confers a relatively larger increase in C/T ratio in *dcp2*Δ vs. WT cells for the mRNA_up_*dcp2*Δ group. *Mechanism (ii)*. The mRNA_up_*dcp2*Δ group is preferentially induced at the transcriptional level and thus resembles the average mRNA both in C/T ratio in WT cells and the increase in C/T ratio in *dcp2*Δ vs. WT cells. The red box depicts the increase in number of transcripts in *dcp2*Δ by this mechanism. **(D-E)** Dcp2-repressed mRNAs exhibit greater than average decapping in WT cells that is reversed by *dcp2*Δ. Notched box-plots of capped to total mRNA ratios in WT cells (D) or in *dcp2*Δ relative to WT cells (E) for all mRNAs (grey) or the following sets of mRNAs: mRNA_up_*dcp2*Δ (blue), mRNA_dn_*dcp2*Δ (red), Dhh1-dependent mRNA_up_*dcp2*Δ (green), Dhh1-independent mRNA_up_*dcp2*Δ (orange). **(F-G)** Quantification of absolute Rpb1 occupancies and mRNA abundance by spike-in normalization reveals reduced Pol II occupancies of most genes showing mRNA derepression in *dcp2*Δ cells. (F) Notched box-plot analysis of changes in Rpb1 occupancies or mRNA abundance in *dcp2*Δ vs. WT cells for the mRNA_up_*dcp2*Δ_A and mRNA_dn_*dcp2*Δ_A groups identified by ERCC-normalized RNA-Seq for all mRNAs (cols. 1-2), mRNA_up_*dcp2*Δ_A (cols. 3-4), and mRNA_dn_*dcp2*Δ_A (cols. 5-6). (G) Hierarchical clustering analysis of the same data from (F) (excluding a few outliers with log_2_Δ values > +4 or < -4).

Because eliminating Dcp2 might provoke other transcriptional responses besides the ESR, we sought additional evidence that reduced mRNA degradation owing to diminished decapping is a major driver of mRNA derepression in *dcp2*Δ cells. Recent evidence indicates that many transcripts decapped by Dcp1/Dcp2 undergo 5’ to 3’ degradation while associated with translating ribosomes (Hu et al. 2009), with Xrn1 following the last translating ribosomes loaded on mRNAs prior to decapping (Pelechano et al. 2015). Such decapped degradation intermediates may account for ∼12% of the polyadenylated mRNA population in WT yeast cells (Pelechano et al. 2013). We reasoned that mRNAs targeted preferentially for decapping and degradation by Dcp2 should be enriched for such uncapped degradation intermediates and hence exhibit a greater than average proportion of uncapped transcripts in WT cells, which will be eliminated in the *dcp2*Δ mutant. In contrast, mRNAs that are derepressed indirectly as the result of elevated transcription in *dcp2*Δ cells should exhibit an average proportion of uncapped isoforms in WT cells. These predictions are illustrated in Figure 3C for an idealized scenario of enhanced mRNA decapping/degradation (model (i)) vs. elevated transcription (model (ii)) to explain the derepression of mRNA_up_*dcp2*Δ transcripts. To evaluate these predictions, we measured the abundance of capped isoforms for all expressed transcripts by CAGE (cap analysis of gene expression), using a revised no-amplification-nontagging technique (nAnT-iCAGE) that eliminates certain artifacts during cDNA library preparation (Murata et al. 2014). We also subjected the same total RNA from WT and *dcp2*Δ cells to standard RNA-Seq and normalized the capped transcripts per million (TPM) determined by nAnT-iCAGE to the TPMs in total mRNA determined by RNA-Seq for each transcript in the two strains. To validate this approach, we similarly compared WT and isogenic *xrn1*Δ cells, reasoning that eliminating 5’ to 3’ degradation of decapped mRNAs by Xrn1 should have the opposite effect from eliminating decapping by *dcp2*Δ and confer higher vs. lower proportions of uncapped isoforms for the set of transcripts preferentially targeted for degradation by Dcp1/Dcp2 and Xrn1.

We observed strong correlations (ρ = 0.99) between biological replicates for WT, *dcp2*Δ and *xrn1*Δ cells, for both nAnT-iCAGE-Seq and parallel RNA-Seq analysis (Figure 3 – figure supplement 1A-F). There is also a strong positive correlation between changes in TPMs for total RNA from RNA-Seq (△mRNA_T) and TPMs of capped RNA from nAnT-iCAGE-Seq (△mRNA_C) in *dcp2*Δ vs. WT (ρ= 0.83) (Figure 3 – figure supplement 2A), and strong overlap between genes showing significant derepression of total mRNAs vs. capped mRNAs in *dcp2*Δ vs. WT cells as determined by DESeq2 analysis (Figure 3 – figure supplement 2C)—all as expected if transcripts derepressed by *dcp2*Δ accumulate as capped isoforms in the mutant cells. In contrast, we found a marked negative correlation between total and capped mRNA changes conferred by *xrn1*Δ (ρ = -0.64) (Figure 3 – figure supplement 2B), and a corresponding under-enrichment of mRNAs significantly derepressed in total vs. capped transcripts (Figure 3 – figure supplement 2D), as expected if the mRNAs derepressed by *xrn1*Δ accumulate as decapped isoforms. The opposite effects of *dcp2*Δ and *xrn1*Δ in these comparisons provide strong validation of this approach to measuring the proportion of capped molecules for each mRNA in different strains.

Calculating the capped to total TPM ratio (C/T) for each mRNA revealed that the mRNA_up_*dcp2*Δ transcripts (derepressed in total RNA by *dcp2*Δ) exhibit a lower proportion of capped transcripts in WT cells compared to all mRNAs (Figure 3D, cols. 1-2), consistent with the “Diminished decay” model (Figure 3C, WT). Furthermore, their C/T ratios are increased by *dcp2*Δ to a greater extent than observed for all mRNAs (Figure 3E, cols. 1-2), as expected if their relatively low C/T ratios in WT result from enhanced decapping (Figure 3C(i), *dcp2Δ* vs. WT). The mRNA_dn_*dcp2*Δ group of transcripts (repressed in abundance by *dcp2*Δ) exhibit the opposite features of a greater than average proportion of capped transcripts in WT (Figure 3D, col. 3 vs. 1) and a less than average increase in the fraction of capped transcripts in *dcp2*Δ vs. WT cells (Figure 3E, col. 3 vs. 1). Extending this analysis to include all mRNAs, which were binned according to their changes in total mRNA abundance between *dcp2*Δ and WT cells, we observed that greater increases in total mRNA levels are generally associated with greater increases in the proportion of capped mRNAs (C/T) conferred by *dcp2*Δ vs. WT (Figure 3 – figure supplement 3B). These findings support the “Diminished decay” model in Figure 3C(i) in the manner expected if loss of decapping and attendant increased mRNA stability is a major driver of increased mRNA abundance in *dcp2*Δ vs. WT cells.

Further supporting this last conclusion, the proportion of capped mRNAs (C/T) for the mRNA_up_*dcp2*Δ transcripts is decreased by *xrn1*Δ to a greater extent than observed for all mRNAs (Figure 3 – figure supplement 3A, col. 4 vs. 2), as predicted from reduced 5’-3’ degradation of the uncapped isoforms generated by Dcp1/Dcp2 in the absence of Xrn1. Also as expected, these mRNAs exhibit changes in C/T ratios in opposite directions in response to *xrn1*Δ vs. *dcp2*Δ, decreasing more than observed for all mRNAs in *xrn1*Δ cells (Figure 3 – figure supplement 3A, col. 4 vs. 2), while increasing more than do all mRNAs in *dcp2*Δ cells (Figure 3 – figure supplement 3A, col. 3 vs. 1). Moreover, examining all mRNAs revealed an indirect correlation between the changes in C/T ratios in *xrn1*Δ vs. WT cells and the changes in total mRNA abundance conferred by *xrn1*Δ (Figure 3 – figure supplement 3C), opposite to the trend observed for *dcp2*Δ (Figure 3 – figure supplement 3B).

The foregoing analysis was also applied to the Dhh1-independent and Dhh1-dependent subsets of the mRNAs derepressed by *dcp2*Δ (described above in Figure 1E-F). Importantly, both groups resemble the entire group of mRNA_up_*dcp2*Δ transcripts in showing lower than average proportions of capped transcripts in WT cells (Figure 3D, cols. 4-5 vs. 2), and a corresponding greater than average increase in C/T ratios in response to *dcp2*Δ (Figure 3E, cols. 4-5 vs. 2). Thus, the cohort of Dhh1-independent mRNAs, which are generally repressed by NMD factors, appear to be targeted for decapping by Dcp2 in WT cells to the same extent as the Dhh1-dependent set of transcripts that are controlled by Dcp2 dependent on Dhh1/Pat1/Edc3 or Scd6.

As an independent assessment of whether the depression of mRNAs by *dcp2*Δ is driven by decapping and 5’-3’ degradation, we determined the codon protection index (CPI) of the mRNA_up_*dcp2*Δ transcripts, previously identified as a measure of co-translational decay for each gene. Decapped mRNA degradation intermediates exhibit three-nucleotide periodicity generated by Xrn1 exonucleolytic cleavage behind the last translating ribosomes, and the CPI metric captures the prevalence of such intermediates for each mRNA (Pelechano et al. 2015). Importantly, the mRNA_up_*dcp2*Δ group exhibits higher than average CPIs, indicating a greater than average involvement of decapping and co-translational degradation by Xrn1 in their decay, whereas the mRNA_dn_*dcp2*Δ transcripts exhibit lower than average CPI values (Figure 3 – figure supplement 4A), consistent with the involvement of an alternative degradation pathway controlling their abundance.

Finally, to investigate the possible contribution of increased transcription to derepression of mRNA abundance by *dcp2*Δ, we performed ChIP-Seq on Rpb1 (a subunit of RNA polymerase II (Pol II)) in WT and *dcp2*Δ cells to measure the relative Pol II occupancies averaged across the coding sequences of each expressed gene, observing excellent correlations between biological replicates (r = 0.99, Figure 3 – source data 3). We first analyzed Rpb1 occupancies for stress-induced genes expected to be derepressed by *dcp2*Δ at the level of mRNA synthesis owing to transcriptional activation by stress-activated transcription factors Msn2/Msn4 (Elfving et al. 2014). Indeed, we observed increased median Rpb1 occupancies for the group of 270 iESR genes, which was substantially greater for the subset found to bind Msn2 in their promoter regions following a shift from glucose to glycerol as carbon source (Msn2-iESR, n=41) (Elfving et al. 2014) compared to the remaining genes lacking detectable Msn2 binding (nonMsn2-iESR, n=225) (Figure 3 – figure supplement 4B). A representative Msn2-iESR transcript, *TPS2*, displays elevated Rpb1 occupancy across the CDS in *dcp2*Δ vs. WT cells that parallels the increased mRNA abundance conferred by *dcp2*Δ determined by RNA-Seq (Figure 3 – figure supplement 4C). In contrast to iESR transcripts, the majority of transcripts derepressed by *dcp2*Δ do not exhibit increased Rpb1 occupancies in the CDS (Figure 3 – figure supplement 4D, mRNA_up_*dcp2*Δ). Although *dcp2*Δ confers a slight increase in median Rpb1 occupancy for the entire group (Figure 3 – figure supplement 4E, cols. 1-2), the fold-change of 1.03 is far below the 1.96-fold increase in mRNA abundance shown in Figure 1B. Furthermore, removal of iESR mRNAs from the group eliminated a significant change in Rpb1 occupancy for the remaining derepressed transcripts (Figure 3 – figure supplement 4E, col. 6). Thus, as illustrated for the transcripts *ATG8, FLO5*, and *CAT8* (Figure 3 – figure supplement 4F), the increased mRNA abundance of most mRNA_up_*dcp2*Δ transcripts appears to result from reduced mRNA turnover rather than increased transcription in *dcp2*Δ cells. Supporting this, the mRNA_up_*dcp2*Δ group is 7-fold depleted for a group of 1907 transcripts that show similar changes in Rpb1 occupancies and mRNA levels conferred by *dcp2*Δ (Figure 3 – figure supplement 4G). Analysis of mRNAs repressed by *dcp2*Δ likewise reveals smaller reductions in Rpb1 occupancies vs. mRNA abundance (Figure 3 – figure supplement 4E, mRNA_dn_*dcp2*Δ), with a reduction in median Rpb1 occupancy of only 0.93-fold (Figure 3 – figure supplement 4E, col. 3) compared to 0.57 for mRNA abundance (Figure 1B), suggesting increased mRNA turnover vs. decreased transcription for these transcripts in *dcp2*Δ cells; which applies to the group of rESR transcripts as well (Figure 3 – figure supplement 4E, col. 5).

The Rpb1 occupancies described above were normalized to the average occupancy in each strain and hence correspond to relative values. To examine absolute changes in Pol II occupancies, we added a spike-in of *S. pombe* chromatin to each *S. cerevisiae* chromatin sample, at 1:10 DNA mass ratio, prior to immunoprecipitation of both *S.cerevisiae* and *S. pombe* Rpb1 with the same antibodies. To measure absolute vs. relative changes in mRNA abundance, we repeated the RNA-Seq experiments by adding a fixed amount of External RNA Controls Consortium (ERCC) transcripts to equal amounts of total RNA from WT and *dcp2*Δ cells prior to preparation of cDNA libraries. Importantly, the spike-in normalized RNA-Seq data revealed a median increase of 24% in bulk mRNA in *dcp2*Δ vs. WT cells (Figure 3F, col. 1), indicating elevated total mRNA levels in cells lacking decapping enzyme. We identified groups of transcripts with >1.4-fold absolute changes in *dcp2*Δ vs. WT cells (FDR < 0.01), dubbed mRNA_up_*dcp2*Δ_A and mRNA_dn_*dcp2*Δ_A, respectively (Figure 3F, cols. 3, 5). (ERCC spike-in shrank the group of repressed mRNAs while expanding the group of derepressed mRNAs because many mRNAs showing reduced abundance relative to the average transcript exhibit increased absolute abundance with spike-in normalization). Analyzing the absolute changes in spike-in normalized Rpb1 occupancies revealed a 1.2-fold reduced median Rpb1 occupancy for all expressed transcripts, indicating a global reduction in transcription rate (Figure 3F, col. 2). This is consistent with previous results indicating that decreased mRNA turnover in mutants lacking mRNA degradation enzymes is buffered by decreased rates of transcription (Sun et al. 2013). Importantly, the derepressed mRNA_up_*dcp2*Δ_A transcripts show reduced absolute Rpb1 occupancies in *dcp2*Δ vs. WT cells (Figure 4F, cols. 3-4), which applies broadly to individual transcripts within the mRNA_up_*dcp2*Δ_A group (Figure 3G). These results confirm our conclusion that mRNAs are derepressed by *dcp2*Δ primarily owing to decreased mRNA turnover rather than increased transcription. The much smaller mRNA_dn_*dcp2*Δ_A group also shows a lower median Rpb1 occupancy, similar in magnitude to its decreased median mRNA abundance (0.76-fold vs. 0.61-fold, Figure 3F, cols. 5-6), suggesting that decreased transcription contributes to the decreased abundance of these repressed transcripts in *dcp2*Δ cells.

**Figure 4.**
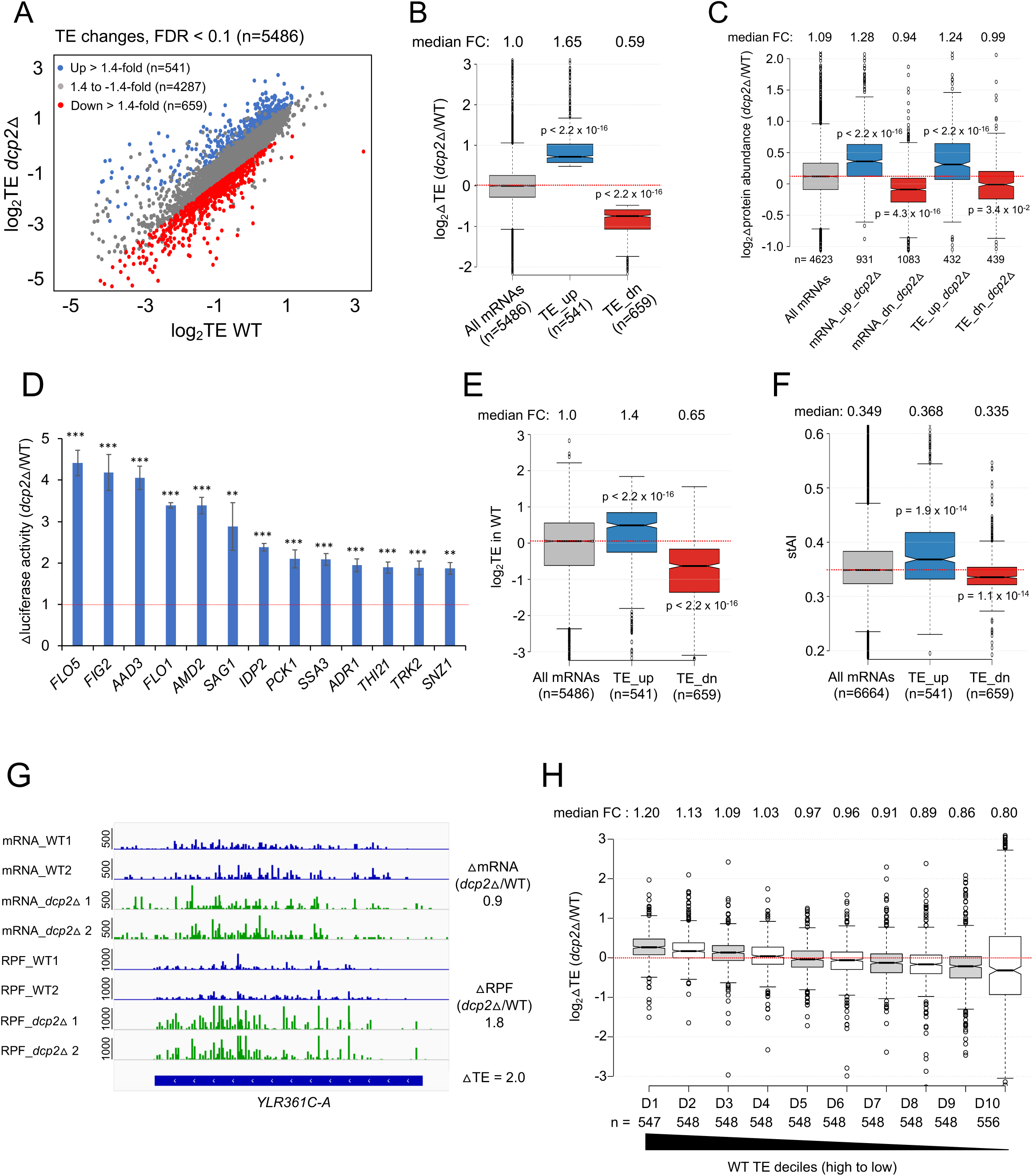
(A) Dcp2 regulates translational efficiencies (TEs) of hundreds of transcripts. **(A)** Scatterplot as in Figure 1A except displaying significant log_2_ fold-changes in TE in *dcp2*Δ vs. WT cells, determined by ribosome profiling. **(B)** Notched box-plot of log_2_ fold-changes in TE (from ribosome profiling) in *dcp2*Δ vs. WT cells for all mRNAs (with median FC of 1.0) and for the 2 sets of TE_up_*dcp2*Δ or TE_dn_*dcp2*Δ transcripts showing translational repression or stimulation, respectively by Dcp2. **(C)** Notched box-plot of log_2_ fold-changes in protein abundance determined by TMT-MS/MS in *dcp2*Δ vs. WT cells for all mRNAs or for the four indicated groups showing Dcp2-mediated repression or stimulation of mRNA abundance or TE. **(D)** Changes in Nano-luciferase activity in *dcp2*Δ vs. WT cells expressed from the indicated 13 *nLUC* reporters. Average values (+/- S.E.M) from at least three biological replicates are shown. Results of t-tests are indicated as: ***, P <0.001, **, P<0.01. **(E-F)** Notched box-plot of log_2_TE in WT (E) and stAI values (F) for all mRNAs or for the two groups showing repression or stimulation of translation by Dcp2. **(G)** Representative gene exhibiting increased TE in *dcp2*Δ vs. WT cells. Integrated Genomics Viewer (IGV, Broad Institute) display of mRNA reads and ribosome-protected fragments (RPFs) across the *YLR361-A* gene from two biological replicates each for WT, and *dcp2*Δ strains, shown in units of RPKM (reads per 1000 million mapped reads). Position of the CDS (blue) is at the bottom with the scale in bp; scales of RPKM for each track are on the left, and calculated ΔmRNA, ΔRPF and ΔTE values between each mutant and WT are on the right. **(F)** Notched box-plots of log_2_ fold-changes in TE in *dcp2*Δ vs. WT across ten deciles of transcripts binned according to TE in WT cells, progressing left to right from highest to lowest TEs.

### Dcp2 modulates the translational efficiencies of many mRNAs

Given its role in decapping-mediated mRNA degradation, most studies on Dcp2 have focused on mRNA decay, and its possible role in translational control is largely unexplored. Hence, we determined the changes in translation for all mRNAs conferred by *dcp2*Δ using our ribosome profiling analysis of the same WT and *dcp2*Δ strains subjected to RNA-Seq (Zeidan et al. 2018). Ribosome profiling (Ribo-Seq) entails deep sequencing of mRNA fragments protected from RNase cleavage by translating 80S ribosomes (RPFs), which when normalized to total mRNA levels, reveals the ribosome density on each mRNA, a measure of relative translational efficiency (TE). Interestingly, DESeq2 analysis revealed that Dcp2 differentially controls the TEs of hundreds of mRNAs, with 541 mRNAs showing higher TEs in *dcp2*Δ vs. WT (TE_up_*dcp2*Δ) (Figure 4A, blue), with a median increase of 1.65-fold (Figure 4B, col. 2); and a similar number (n=659) displaying reduced TEs in *dcp2*Δ cells (TE_dn_*dcp2*Δ) (Figure 4A, red) with a median decrease of 0.59-fold (Figure 4B, col. 3). These results suggest that Dcp2 broadly controls gene expression at the translational level in addition to regulating mRNA stability.

In an effort to establish that changes in TE are generally associated with changes in the synthesis and abundance of the encoded proteins, we measured the changes in steady-state levels of individual proteins by Tandem Mass Tag Mass spectroscopy (TMT-MS/MS) of total protein samples isolated from the *dcp2*Δ and WT strains. In this approach, all peptides in the mutant and WT extracts are covalently labeled with different isobaric mass tags, which have the same mass but yield reporter ions of different masses in the tandem MS mode during peptide fragmentation. Analyzing a mixture of the differentially labeled mutant and WT samples yields the ratios of peptide abundance in the two strains. Using TMT-MS/MS data from three biological replicates of each strain, we determined changes in abundance for ∼4600 different proteins, with the three replicates showing similar distributions of peptide abundance and highly reproducible levels across all detected proteins (Figure 4 – figure supplement 1A-C). Importantly, a positive correlation exists between the relative changes in protein abundance from TMT-MS/MS and changes in RPFs from Ribo-Seq in *dcp2*Δ vs. WT cells (ρ = 0.6) (Figure 4 – figure supplement 1D). Furthermore, the groups of mRNAs defined above showing significant changes in mRNA abundance or TEs conferred by *dcp2*Δ also displayed changes in protein levels in the same directions (Figure 4C). Considering that steady state protein abundance is controlled by rates of degradation in addition to rates of synthesis, the substantial correspondence found between Ribo-Seq and TMT/MS-MS data indicates that the changes in ribosome occupancies (RPFs) measured by Ribo-Seq generally signify corresponding changes in translation rates between *dcp2*Δ and WT cells. To provide additional support for this last conclusion, we analyzed the expression of Nano-luciferase (*nLUC*) reporters constructed for particular genes by inserting the *nLUC* coding sequences immediately preceding the stop codon, in-frame with the main ORF of each gene, preserving the native 5’UTR and 3’UTR sequences. We observed increased luciferase expression in cell extracts of *dcp2*Δ vs. WT transformants harboring the reporter plasmids constructed for 13 different genes that showed increased RPFs in *dcp2*Δ vs. WT cells (Figure 4D).

Examining the properties of mRNAs displaying increased translation in *dcp2*Δ vs. WT cells, we found that the TE_up_*dcp2*Δ transcripts tend to have shorter CDS lengths, longer half-lives, and greater mRNA abundance compared to all mRNAs (Figure 4 – figure supplement 2A-C). These features are associated with efficiently translated mRNAs and, indeed, the TE_up_*dcp2*Δ mRNAs have greater than average TEs in WT cells (Figure 4E) and are enriched for optimal codons (Figure 4F). Consistent with this, the TE_dn_*dcp2*Δ mRNAs have properties opposite of those exhibited by the TE_up_*dcp2*Δ group, including longer CDS, shorter half-lives and lower transcript abundance compared to all mRNAs (Figure 4 – figure supplement 2A-C), and lower than average TE in WT (Figure 4E) and frequency of non-optimal codons (Figure 4F). Extending this analysis to include all expressed mRNAs, which were sorted into 10 bins on the basis of their TEs in WT cells, revealed a direct correlation between TE changes conferred by *dcp2*Δ and TE in WT (Figure 4H), supporting the notion that Dcp2-translationally repressed mRNAs tend to be well-translated, whereas Dcp2-translationally activated transcripts are generally poorly-translated in WT cells.

### Dcp2-translationally repressed transcripts generally do not accumulate as decapped low-TE species in WT cells

The mRNAs encoded by *YLR361-A* and *YLR297A* are representative transcripts exhibiting TE increases conferred by *dcp2*Δ of 2.0- and 2.7-fold, respectively, but display either no significant change or a considerably smaller increase in mRNA abundance in *dcp2*Δ vs. WT cells (Figure 4G; Figure 4 – figure supplement 2D). This suggests that Dcp2 represses their translation without targeting these mRNAs for degradation. In fact, *dcp2*Δ generally confers the opposite effects on mRNA abundance and TE for the cohort of mRNAs it regulates translationally, as the TE_up_*dcp2*Δ group shows a decreased median mRNA level, while the TE_dn_*dcp2*Δ group of mRNAs shows an increased median mRNA level in *dcp2*Δ vs. WT cells (Figure 5A). The finding that mRNAs translationally repressed by Dcp2 appear to have a less than average dependence on Dcp2 for mRNA decay led us to consider whether the TE_up_*dcp2*Δ transcripts are in fact preferentially decapped by Dcp2 but are not targeted for degradation, such that their overall mRNA abundance is not repressed by Dcp2. If so, the uncapped mRNAs persisting in WT cells would have a low TE in WT cells owing to their inability to bind eIF4F and become activated for translation initiation, and their TEs would increase in *dcp2*Δ cells as they would remain fully capped and capable of binding eIF4F (see model in Figure 5B).

**Figure 5:**
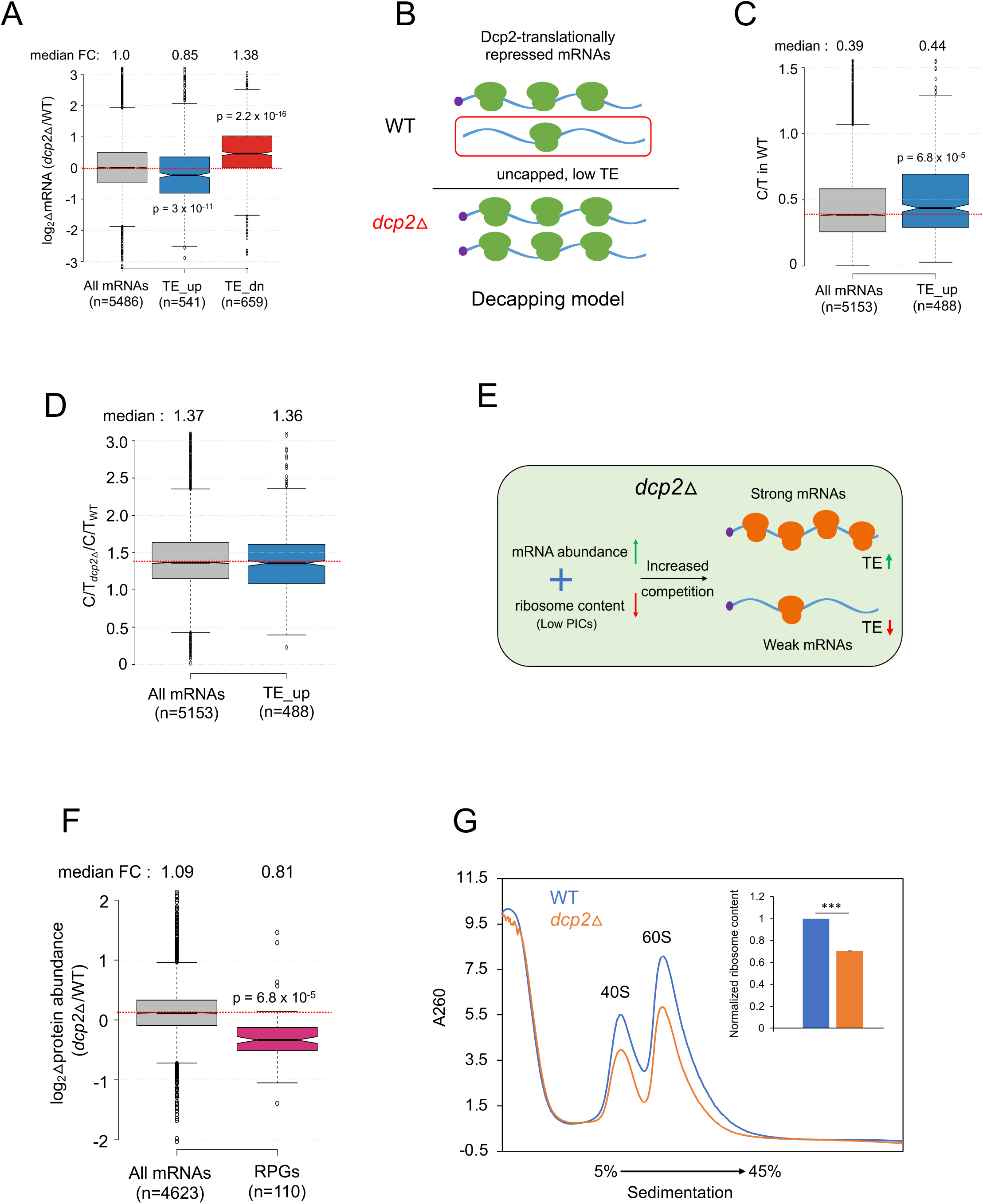
Evidence that the majority of TE changes conferred by *dcp2*Δ result from increased competition for limiting PICs owing to diminished ribosome production and elevated mRNA levels. **(A)** Notched box-plot showing log_2_ fold-changes in mRNA abundance in *dcp2*Δ relative to WT cells for all mRNAs, or the mRNAs translationally repressed (Blue) or stimulated (red) by Dcp2 in WT cells. **(B)** Hypothetical schematic model to explain TE increases conferred by *dcp2*Δ resulting from the persistence of translationally inert, decapped intermediates in WT cells. In WT (upper), the TE_up_*dcp2*Δ group of mRNAs is preferentially targeted by Dcp2 for decapping and these uncapped species cannot bind eIF4F and thus exhibit low TEs. In *dcp2*Δ cells (lower), decapping is eliminated and the low-TE, decapped fraction no longer exists, which increases the overall TE of the transcript pool. **(C-D**) Notched box-plots of C/T ratios in WT cells (C) and C/T ratios in *dcp2*Δ vs. WT cells (D) for all mRNAs and the group translationally repressed by Dcp2. (E) Schematic of the PIC competition model proposed to explain the broad reprogramming of TEs conferred by *dcp2*Δ. A combination of diminished ribosome production resulting from down-regulation of rESR transcripts and elevated bulk capped mRNAs resulting from loss of decapping-mediated mRNA turnover evokes increased competition among all mRNAs for limiting PICs, producing relatively greater translation of “strong” efficiently translated mRNAs at the expense of “weak” poorly translated mRNAs. **(F)** Notched box-plot showing log_2_ fold-changes in protein abundance determined by TMT-MS/MS in *dcp2*Δ vs. WT cells for all mRNAs or those from 110 genes encoding ribosomal proteins (RPGs). **(G)** Quantification of total 40S and 60S ribosomal subunits in *dcp2*Δ vs. WT cells. Representative A_260_ profiles of equal proportions of cell extracts obtained from WT (blue) and *dcp2*Δ (orange) cultures are shown. The inset summarizes the combined areas under the 40S and 60S peaks normalized to the OD_600_ of the cell cultures calculated from three biological replicates of *dcp2*Δ and WT cells, setting the mean WT value to unity. An unpaired Student’s t-test indicates a highly significant difference in the means calculated from the biological replicates of the two different strains (***, p < 0.001).

This model makes testable prediction that the TE_up_*dcp2*Δ mRNAs should exhibit a lower than average proportion of capped mRNA (C/T ratio) in WT cells, and a greater than average increase in C/T ratios in *dcp2*Δ vs. WT cells owing to loss of decapping in the mutant (Figure 5B). Instead, the TE_up_*dcp2*Δ mRNAs have somewhat higher, not lower, than average C/T ratios in WT cells (Figure 5C), and they show an increase in C/T ratio in *dcp2*Δ vs. WT cells indistinguishable from that seen for all mRNAs (Figure 5D). This finding do not support the hypothesis that most TE_up_*dcp2*Δ transcripts have a higher than average proportion of uncapped transcripts of low TE in WT cells, whose translation is rescued by loss of decapping in *dcp2*Δ cells.

### Evidence that competition for limiting ribosomes reprograms translation in *dcp2*Δ cells

We considered an alternative possibility that *dcp2*Δ confers TE changes as an indirect consequence of elevated mRNA levels resulting from loss of the major pathway for mRNA degradation. This idea was prompted by our finding above that the mRNAs showing TE increases in *dcp2*Δ cells tend to be efficiently translated in WT cells, whereas mRNAs poorly translated in WT tend to show TE reductions in the mutant (Figures 4E & H). Previously, we observed this same pattern of translational reprogramming in yeast cells impaired in different ways for assembly of 43S preinitiation complexes (PICs), including: (i) increased phosphorylation of eIF2*α*in WT cells induced by isoleucine/valine starvation using the drug sulfometuron (SM), which decreases formation of the eIF2-GTP-Met-tRNA_i_ ternary complex required to assemble 43S PIC; (ii) deletion of genes *TMA64* and *TMA20* encoding factors that recycle 40S subunits from termination complexes at stop codons to provide free 40S subunits for PIC assembly; and (iii) a reduction in free 40S subunits by depleting a 40S subunit protein. This recurrent pattern of translational reprogramming could be explained by proposing that increased competition for limiting PICs allows “strong” mRNAs, highly efficient in recruiting PICs, to outcompete weak mRNAs that recruit PICs less efficiently (Gaikwad et al. 2021). Supporting the possibility that a similar competition exists among mRNAs translationally altered by *dcp2*Δ, we found that the TE_up_*dcp2*Δ group of mRNAs also shows an increased median TE in response to increased phosphorylation of eIF2*α* induced by SM (WT_SM) and by deletion of *TMA64/TMA20* (*tma*ΔΔ), whereas the TE_dn_*dcp2*Δ mRNAs exhibit the opposite changes in median TE in response to these two conditions (Figure 5 – figure supplement 1A). Hierarchical clustering of TE changes for individual mRNAs reveals that the majority of mRNAs showing significant TE changes in *dcp2*Δ vs. WT cells exhibit changes in the same direction (same color) in response to the *tma*ΔΔ mutations or SM treatment of WT cells (Figure 5 – figure supplement 1B). There are numerous exceptions to this trend, however, suggesting that perturbations specific to each condition can differentially affect the translation of specific subsets of mRNAs and alter their responses to increased competition for limiting PICs.

We considered that increased competition among mRNAs for limiting PICs might exist in *dcp2*Δ cells for two main reasons: i) impairing decapping-mediated mRNA decay will stabilize most transcripts and elevate bulk cellular mRNA abundance; and ii) reduced ribosome content arising from the repression of rESR mRNAs, which encode primarily ribosomal proteins and ribosome biogenesis factors (Gasch et al. 2000). Indeed, a recent study demonstrated reduced ribosome content in mutant cells undergoing the ESR response (Terhorst et al. 2020). We hypothesized that the combination of increased mRNA abundance and reduced ribosomal content in *dcp2*Δ cells will increase competition among mRNAs for limiting PICs and confer the observed translational reprogramming that favors strong over weak mRNAs (Figure 5E).

As mentioned above, our spike-in normalized RNA-Seq data revealed a median increase of 24% in bulk mRNA in *dcp2*Δ vs. WT cells (Figure 3F), indicating elevated mRNA abundance in cells lacking Dcp2. Considering that all mRNAs should be capped in *dcp2*Δ cells, whereas a fraction of mRNAs are uncapped in WT, the increase in capped mRNA abundance conferred by *dcp2Δ* should be even greater than 24%. To explore whether *dcp2*Δ reduces ribosome content, we first interrogated the ribosome occupancies measured by Ribo-Seq, a measure of relative protein synthesis rates, and found that the group of 139 ribosomal proteins exhibits ∼2-fold lower RPFs in *dcp2*Δ vs. WT (Figure 5 – figure supplement 1C). Consistent with this, TMT-MS/MS analysis revealed a ∼25% reduction in median steady-state abundance of ribosomal proteins in *dcp2*Δ relative to WT cells compared to all proteins (Figure 5F). We then measured the abundance of assembled ribosomal subunits by resolving total 40S and 60S subunits in whole cell extracts of mutant and WT cells by sedimentation through sucrose density gradients, using an extraction buffer lacking Mg^+2^ to dissociate 80S ribosomes into free 40S and 60S subunits. Normalizing the A_260_ absorbance tracings of ribosomal subunits to the number of OD_600_ units (cellular volume) of extracted cells, revealed ∼30% lower levels of both 40S and 60S subunits in *dcp2*Δ vs. WT cells (Figure 5G). Thus, all three measurements of ribosome synthesis or abundance indicate a significant reduction in ribosome levels in *dcp2*Δ vs. WT cells that when coupled to the increased capped mRNA content in the mutant, should increase competition among mRNAs for limiting PICs. Based on our previous studies (Gaikwad et al. 2021), this can be expected to reprogram translation to favor well-translated mRNAs and, hence, may account for most of the TE increases or decreases conferred by *dcp2*Δ.

### Dhh1 is required for translational repression of a subset of transcripts by Dcp2

Having found evidence that the bulk of TE changes conferred by *dcp2*Δ could arise from increased competition for limiting ribosomes, we wondered whether Dhh1 contributes to this indirect mechanism of translational reprogramming. To address this question, we asked first whether mRNAs translationally dysregulated by *dcp2*Δ are dependent on Dhh1 for their translational reprogramming. Cluster analysis of the TE changes observed in the *dcp2*Δ, *dhh1*Δ, and *dhh1*Δ*dcp2*Δ mutants relative to WT revealed that most transcripts showing increased TEs in the *dcp2*Δ single mutant exhibit little increase, or even decreases, in TE in the *dhh1*Δ single mutant, and also show similar TE increases in the *dhh1*Δ*dcp2*Δ double mutant compared to the *dcp2*Δ single mutant (Figure 6A, groups with orange bars), all indicating that they are translationally controlled by Dcp2 independently of Dhh1. Indeed, examining the TE changes for this subset of Dcp2-translationally repressed mRNAs reveals nearly identical increases in median TE in the *dhh1*Δ*dcp2*Δ double and *dcp2*Δ single mutant of ∼1.6-fold, but only a slight increase in median TE of ∼1.1-fold in the *dhh1*Δ single mutant (Figure 6B, cols. 4-6). Interestingly, the remaining one-fourth of TE_up_*dcp2*Δ transcripts exhibit similar TE increases in all three deletion mutants (Figure 6A, groups with green bars), with nearly identical increases in median TE of ∼2.3-fold relative to WT (Figure 6B, cols. 1-3). Thus, this latter group of mRNAs appears to be dependent on Dhh1 for translational repression by Dcp2, such that eliminating either factor individually or in combination confers nearly the same derepression of TEs. Interestingly, the Dhh1-dependent subset shows considerably greater translational repression by Dcp2 compared to the Dhh1-independent subset, with *dcp2*Δ conferring median TE increases of 2.3-fold compared to only 1.6-fold for the Dhh1-independent group of transcripts (Figure 6B, compare cols. 1 and 4).

**Figure 6:**
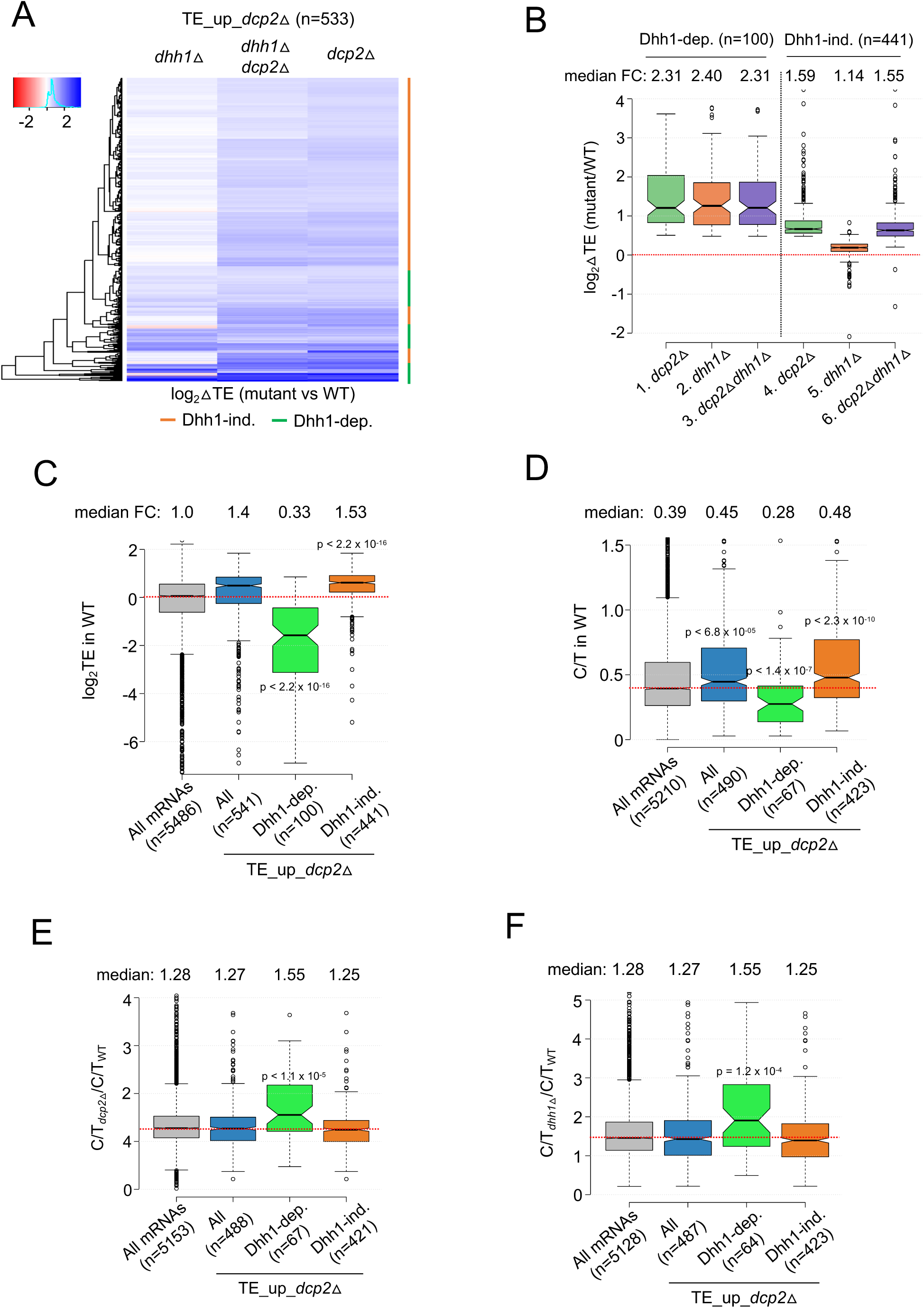
mRNAs exhibiting Dhh1-dependent translational repression by Dcp2 are poorly translated mRNAs preferentially targeted for decapping by Dcp2 and Dhh1. **(A)** Hierarchical clustering analysis of log_2_ fold-changes in TE conferred by the mutations listed across the top vs. WT for the TE_up_*dcp2*Δ mRNAs (excluding a few outliers with log_2_ΔTE values > +4 or < -4). Dhh1-dependence or Dhh1-independence for the TE changes is indicated on the right with green or red bars, respectively. **(B)** Notched box-plot of log_2_ fold-changes in TE conferred by the mutations listed across the bottom vs. WT for the Dhh1-dependent TE_up_*dcp2*Δ (col. 1-3) or Dhh1-independent TE_up_*dcp2*Δ (col. 4-6) groups of mRNAs **(C-F)** Notched box-plot of log_2_ TE in WT (C), C/T ratios in WT (D), C/T ratios in *dcp2*Δ vs. WT (E), and C/T ratios in *dhh1*Δ vs. WT (F) for all mRNAs, all TE_up_*dcp2*Δ mRNAs, or the Dhh1-dependent or Dhh1-independent subsets of the TE_up_*dcp2*Δ mRNAs.

The Dhh1-independent subset of TE_up_*dcp2*Δ transcripts exhibit higher than average median TEs in WT cells (Figure 6C, col. 4 vs. 1), suggesting that their TE increases in *dcp2*Δ cells conform to the PIC competition model, wherein efficiently translated mRNAs outcompete poorly-translated mRNAs for limiting ribosomes in *dcp2*Δ cells (Figure 5E). The Dhh1-dependent subset, by contrast, have a much lower than average median TE in WT cells (Figure 6C, col. 3 vs. 1), suggesting that their translation is repressed by Dcp2 by a different mechanism that involves Dhh1.

We considered that Dhh1-dependent translational repression might involve Dhh1-stimulated decapping and persistence of the decapped mRNAs in a low-TE state envisioned in the decapping model of Figure 5B. Supporting this idea, the Dhh1-dependent subset of TE_up_*dcp2*Δ transcripts exhibit a lower proportion of capped transcripts (C/T ratio) in WT cells compared to both all mRNAs and the Dhh1-independent group of Dcp2-translationally repressed mRNAs (Figure 6D, col. 3 vs. 1 & 4). Moreover, they exhibit a greater than average increases in C/T ratios in both *dcp2*Δ and *dhh1*Δ vs. WT cells (Figure 6E-F, col. 3 vs. 1 & 4), as expected if Dhh1-stimulated decapping by Dcp2 underlies their preponderance of decapped isoforms in WT cells. The levels of total mRNA for this group are also increased by *dcp2*Δ and *dhh1*Δ (Figure 6 – figure supplement 1B & C, col. 3 vs. 1-2), in the manner expected if they are preferentially targeted for mRNA degradation by decapping and 5’-3’ decay in addition to being translationally repressed by Dcp2/Dhh1. The Dhh1-independent group of translationally repressed transcripts, by contrast, exhibit lower than average or average repression of their abundance by Dcp2 and Dhh1 (Figure 6 – figure supplement 1B & C, cols. 4 vs. 1). These 100 mRNAs are not enriched for non-optimal codons, exhibiting a median stAI indistinguishable from all mRNAs (Figure 6 – figure supplement 1A, col. 3 vs. 1), suggesting that targeting of stalled elongating ribosomes by Dhh1 (Radhakrishnan et al. 2016) also does not underlie their translational repression.

### Dcp2 represses the abundance or translation of mRNAs encoding proteins involved in catabolism of alternative carbon sources or respiration

The mRNA_up_*dcp2*Δ transcripts derepressed in abundance by *dcp2*Δ are functionally enriched for stress response genes, reflecting mobilization of the iESR (Gasch et al. 2000) in this slow-growing mutant. Interestingly, these mRNAs are also enriched for genes involved in metabolism of the energy reserves glycogen and trehalose (Figure 7 – figure supplement 1A), the tricarboxylic acid (TCA) cycle (involved in respiration in mitochondria), or meiotic recombination (Figure 7 – figure supplement 1A), of which only two genes belong to the iESR (Figure 7 – figure supplement 3A-B). The subset showing Dhh1-dependent repression of mRNA abundance by Dcp2 show enrichment for the same functional categories except for meiotic recombination, which is enriched among the Dhh1-independent subset of mRNA_up_*dcp2*Δ transcripts instead, along with genes involved in DNA repair and cell-cell adhesion (Figure 7 – figure supplement 1C-D). This suggests a functional bifurcation among Dcp2-repressed mRNAs based on involvement of Dhh1 in the degradation mechanism. Interestingly, the transcripts showing increased TEs in *dcp2*Δ cells (TE_up_*dcp2*Δ) are also enriched for genes involved in respiration, and for ribosomal protein genes (RPGs) (Figure 7 – figure supplement 1E). As most of these mRNAs are well-translated in WT cells, their TE increases can be attributed to the competition mechanism of translational reprogramming in *dcp2*Δ cells (Figure 5E).

To integrate the outcome of altered mRNA abundance and TEs and obtain a measure of altered protein synthesis rates, we identified the groups of mRNAs showing significantly changed ribosome occupancies in *dcp2*Δ vs. WT cells and subjected them to GO analysis. The mRNAs exhibiting increased RPFs (Ribo_up_*dcp2*Δ) are enriched for genes involved in the same categories mentioned above, including stress response factors or proteins involved in metabolism of energy reserves or respiration, but also for metabolism of vitamins or cofactors or for glutamate biosynthesis (Figure 7 – figure supplement 1G). Derepression by *dcp2*Δ of the stress response gene *SSA3* and two genes involved in vitamin biosynthesis, *THI21* (thiamine) and *SNZ1* (pyridoxine), was recapitulated with the corresponding *nLUC* reporters (Figure 4D). Examining 37 genes involved in metabolism of energy reserves reveals derepression by *dcp2*Δ almost entirely at the level of mRNA abundance (Figure 7A), epitomized by the trehalose biosynthetic gene *TPS2* (Figure 7 – figure supplement 2A). In contrast, 52 genes whose products function directly in respiration as components of the electron transport chain, TCA cycle or mitochondrial ATPase, are derepressed primarily at the level of TE (Figure 7B), as exemplified by the TCA cycle gene *LSC1* (Figure 7 – figure supplement 2B). Supporting the latter, Western blot analysis revealed increased steady-state levels in *dcp2*Δ cells of five mitochondrial proteins involved in respiration, Aco1, Atp20, Cox14, Idh1, and Pet10, relative to the glycolytic enzyme GAPDH (Figure 7C).

**Figure 7:**
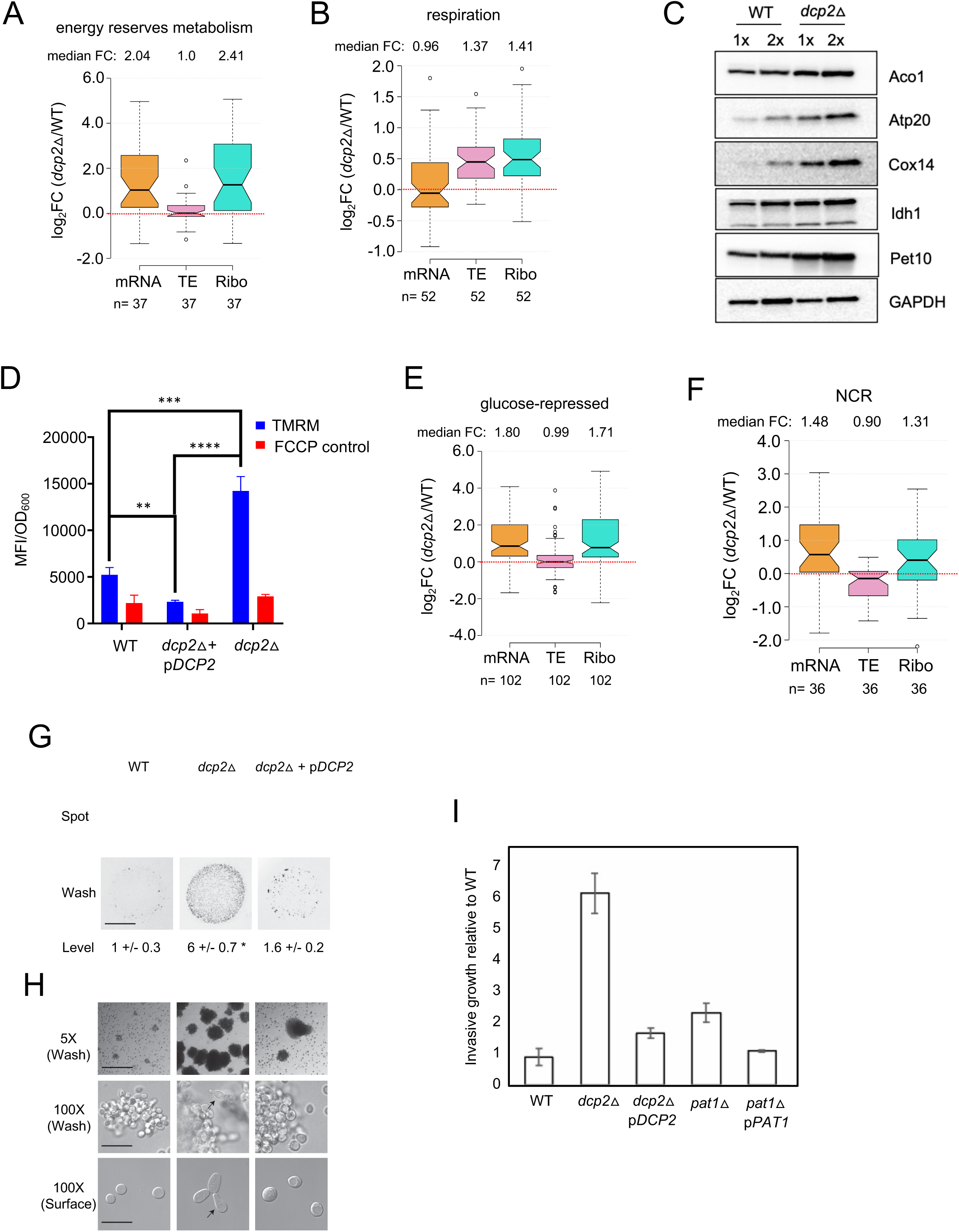
Dcp2 represses genes involved in respiration, catabolism of non-preferred carbon or nitrogen sources, or invasive growth, on rich medium (A-B, and E-H). Notched box-plot showing log_2_ fold-changes in mRNA, TE or RPFs (Ribo) in *dcp2*Δ vs. WT cells for 37 genes involved in metabolism of energy reserves (identified by KEGG pathway) (A), 52 genes encoding mitochondrial proteins with direct roles in oxidative phosphorylation (B), 102 genes subject to glucose repression or induced by Adr1 or Cat8 in low glucose (Young et al. 2003; Tachibana et al. 2005) (E), and 36 genes subject to nitrogen-catabolite repression (Godard et al. 2007) (F). **(C)** Western blot analyses of five mitochondrial proteins indicated on the right involved in respiration (and GAPDH examined as a loading control) in WCEs from WT or *dcp2*Δ cells, with adjacent lanes differing 2-fold in amount of extract. Immune complexes were visualized using enhanced chemiluminescence. The results shown here are representative of three biological replicates that gave highly similar results, presented in Figure 7 – source data 2. **(D)** *dcp2*Δ confers increased mitochondrial ΔΨ_m_. Cells were cultured to mid-log phase. TMRM (500 nM) was added and incubated for 30 min before samples were collected and washed once with deionized water. ΔΨ_m_ was determined by measuring TMRM fluorescence intensity using flow cytometry. Data are presented in arbitrary fluorescence intensity units per OD_600_. 2-way ANOVA was used for statistical analysis and data are given as mean values ± SD (n=3) (****p<0.0001) **(G-H)** *dcp2*Δ confers increased invasive growth. (E) The top and bottom panels show cells spotted on YPD agar medium and grown to confluence before or after washing under water, respectively, revealing increased invasive growth in the agar for the *dcp2*Δ strain compared to WT or the *dcp2*Δ strain complemented with WT *DCP2* on a plasmid (p*DCP2*). The levels of invasive growth were quantified and indicated below the images. (F) The three panels show colony or cell morphology at 5X (after wash), 10X (after wash) and 100X (surface) magnification, respectively, for the strains analyzed in (E). **(I)** Fold-change in invasive growth in *dcp2*Δ, *pat1*Δ and respective complemented strains with WT copy of gene relative to WT cells. Error represents the standard deviation. Significance was determined by Student’s t-test, p-value < 0.05.

To obtain independent evidence that *dcp2*Δ increases respiratory function, we measured the mitochondrial membrane potential (ΔΨ_m_), produced by the electron transport chain, using the probe tetramethylrhodamine (TMRM)—a cationic fluorescent dye that accumulates in mitochondria as a function of ΔΨ_m_. Quantifying dye fluorescence by flow cytometry revealed a marked increase in TMRM fluorescence in the *dcp2*Δ mutant at levels substantially greater than the background signals observed when ΔΨ_m_ was dissipated by addition of the uncoupler FCCP (carbonylcyanide p-trifluoromethoxyphenylhydrazone). The increased TMRM fluorescence was fully complemented by plasmid-borne WT *DCP2* (Figure 7D), providing evidence that *dcp2*Δ increases mitochondrial ΔΨ_m_.

In WT cells growing on medium replete with glucose, such as YPD, respiration is suppressed and energy is produced by fermentation. Proteins involved in catabolism of alternative carbon sources are also repressed on YPD medium. Interestingly, *dcp2*Δ confers increased median translation (RPFs) of a group of 102 mRNAs shown to be glucose-repressed in WT cells and/or activated by the transcriptional activators Adr1 or Cat8 that function in catabolism of non-glucose carbon sources (Young et al. 2003; Tachibana et al. 2005), which is achieved primarily via increased transcript levels (Figure 7E). In fact, *ADR1* (Figure 7 – figure supplement 2C) and *CAT8* (Figure 7 – figure supplement 2D) are themselves derepressed by *dcp2*Δ, which might contribute to the induction of their target gene transcripts in high-glucose medium observed here. The derepression by *dcp2*Δ of *ADR1*, as well as of *IDP2* and *PCK1*, encoding glucose-repressed enzymes of the glyoxylate cycle and gluconeogenesis, respectively, was recapitulated with the corresponding *nLUC* reporters (Figure 4D).

### Dcp2 represses the expression of genes involved in catabolism of alternative or nitrogen sources, in autophagy, and in invasive growth on rich medium

In addition to derepressing genes involved in catabolism of non-glucose carbon sources, we observed increased mRNA levels and translation in *dcp2*Δ cells for a group of 36 nitrogen-catabolite repressed (NCR) genes, which are transcriptionally down-regulated by the presence of the preferred nitrogen sources present in YPD medium (Godard et al. 2007) (Figure 7F). Related to this finding, a group of 24 *ATG* genes directly involved in autophagy show elevated median mRNA levels and translation in *dcp2*Δ cells (as exemplified by *ATG8;* Figure 7 – figure supplement 2F-G). This last observation is in line with previous observations indicating a role for mRNA decapping and degradation in suppressing autophagy in non-starvation conditions (Hu et al. 2015), where salvaging amino acids from extraneous proteins is not adaptive.

Interestingly, *dcp2*Δ cells exhibit a flocculation phenotype, wherein cells stick together and settle to the bottom of a liquid culture. Flocculation typically results from increased cell adhesion due to the up-regulation of *FLO* genes, whose products promote cell adhesion. Indeed, *dcp2*Δ confers increased mRNA levels and translation for a group of 16 genes encoding cell wall proteins, which include multiple *FLO* gene products and other agglutinins (exemplified by *FLO5;* Figure 7 – figure supplement 2E & H). Supporting these results, four of these genes displayed increased expression of the corresponding *nLUC* reporters in *dcp2*Δ cells (*FLO5*, *FLO1, FIG2 and SAG1*, Figure 4D). Cell adhesion is critical for filamentous growth wherein cells switch from separated round cells to adhesion-linked “chains” of elongated cells, which allows them to penetrate substrates (invasive growth). The *dcp2*Δ mutant showed elevated invasive growth compared to WT in a plate-washing assay, which was complemented by WT *DCP2* on a plasmid (Figure 7G). Quantitation of invasive growth across separate trials showed that *dcp2*Δ conferred a 6-fold increase compared to WT, which was largely diminished by plasmid-borne *DCP2* (Figure 7I). Microscopic examination revealed that the *dcp2*Δ mutant formed clumps of cells that were larger and more abundant than WT; and that *dcp2*Δ cells exhibit an elongated morphology, which generally results from enhanced apical growth due to a delay in the cell cycle during filamentous growth (Kron et al. 1994) (Loeb et al. 1999). This last phenotype was observed both in cells scraped from colonies before washing (Figure 7H, 100X, Surface) and in cells excised from the invasive scar after washing (Figure 7H, 100X, Wash). These cellular phenotypes, all of which were diminished by plasmid-borne *DCP2*, suggest that Dcp2 controls other aspects of filamentous/invasive growth besides cell adhesion. Importantly, we observed similar albeit less pronounced invasive growth phenotypes in the mutant lacking decapping activator Pat1 (Figure 7I and Figure 7 – figure supplement 2I-J). Thus, mRNA decapping by Dcp1/Dcp2 and its activation by Pat1 contributes to the control of filamentous growth by the regulation of cell adhesion genes and other associated mechanisms. Only small fractions of the sets of agglutinin genes, autophagy genes, and the NCR or glucose-repressed genes that are derepressed by *dcp2*Δ belong to the iESR (Figure 7 – figure supplement 3C-F).

### Dcp2 stimulates expression of genes involved in protein synthesis, glycosylation, and the unfolded protein response on rich medium

In addition to reducing production of ribosomes, *dcp2*Δ also diminishes mRNA and RPF levels of genes encoding various translation initiation and elongation factors, and of proteins involved in sulfate assimilation into methionine and cysteine (Figure 7 – figure supplement 1B & H), which together should reduce rates of protein synthesis in the mutant cells. The lower expression of sulfur assimilation genes involves reductions in both transcript abundance and TE (Figure 7 – figure supplement 4A), as exemplified for *MET5* (Figure 7 – figure supplement 4B). The abundance of mRNAs and RPFs for numerous genes involved in protein glycosylation also are reduced by *dcp2*Δ (Figure 7 – figure supplement 1B & H; Figure 7 – figure supplement 4C). The suppression of both synthesis and glycosylation of trans-membrane and secreted proteins in the endoplasmic reticulum predicted by these reductions might explain the observed down-regulation by *dcp2*Δ of genes involved in the unfolded protein response (UPR) (Figure 7 – figure supplement 1B & H, Figure 7 – figure supplement 4D). These include the protein kinase encoded by *IRE1* that promotes splicing and activation of the UPR transcription factor Hac1, which is down-regulated at the levels of mRNA and translation (Figure 7 – figure supplement 4E). Figure 8 contains a summary of the major pathways that are elevated or suppressed on elimination of mRNA decapping in *dcp2*Δ cells, as described above.

**Figure 8:**
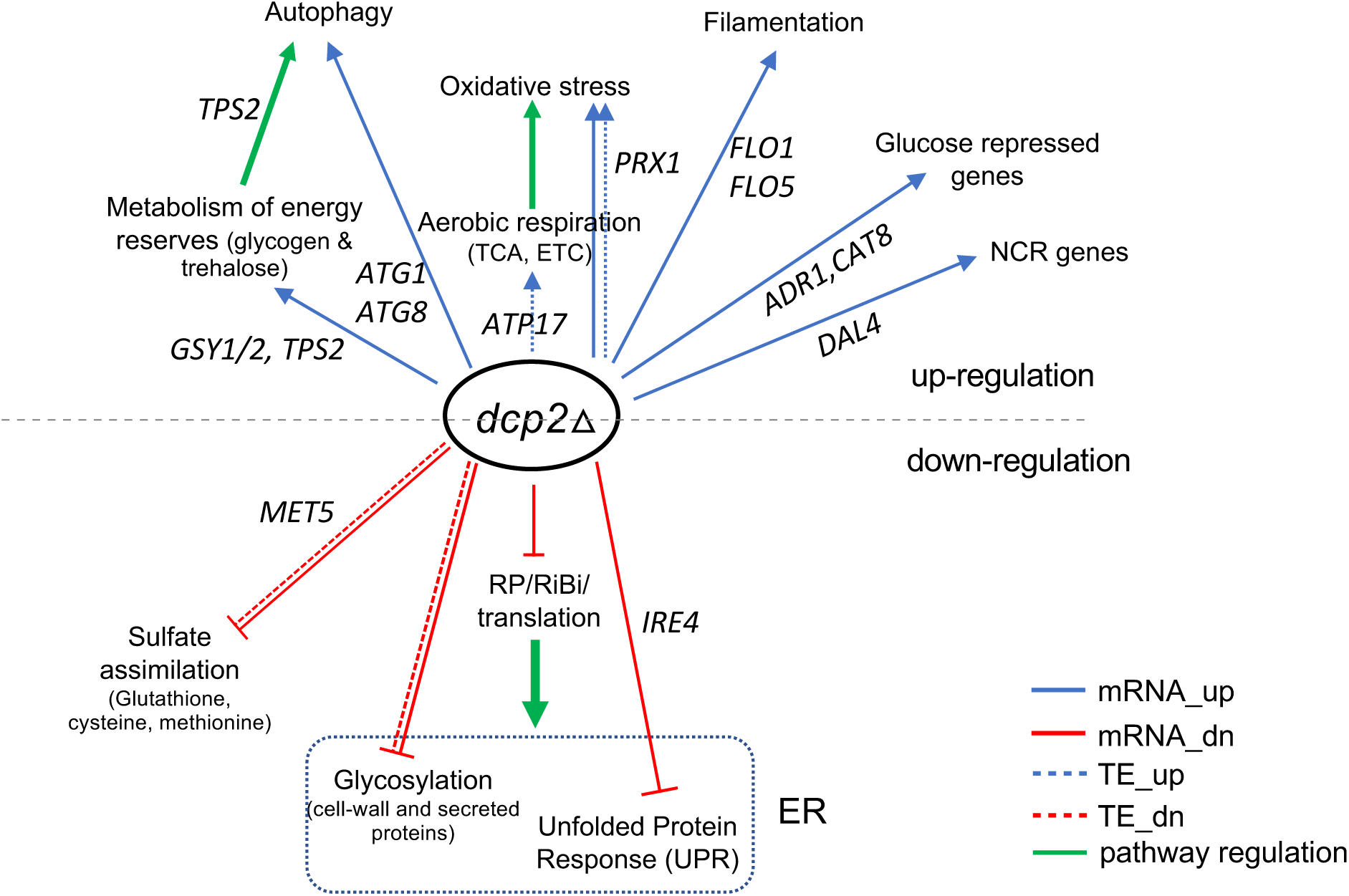
Summary of genes and pathways dysregulated by *dcp2*Δ. Categories above the dotted line are genes or pathways up-regulated by *dcp2*Δ, while those at the bottom are down-regulated. Straight and dotted lines represent changes in mRNA abundance and TE, respectively. Green arrows indicate pathway regulation. Important genes in each functional category are shown. ER, endoplasmic reticulum.

## DISCUSSION

In this study we analyzed the changes in mRNA abundance and translation in the yeast translatome conferred by eliminating or impairing the activity of the catalytic subunit of the mRNA decapping enzyme, Dcp2, in both WT cells and mutant cells lacking decapping activator and translational repressor Dhh1. Our analysis of mRNA changes revealed that roughly one-half of the mRNAs derepressed in abundance by *dcp2*Δ were similarly derepressed by *dhh1*Δ, with no further derepression seen in the *dcp2*Δ*dhh1*Δ double mutant (Figure 1E-F). These are the results expected if these mRNAs are dependent on Dhh1 to stimulate their decapping by Dcp1/Dcp2 and attendant 5’-3’ degradation by Xrn1, such that eliminating Dhh1, Dcp2, or both proteins together, stabilizes these transcripts to similar degrees. Interestingly, these Dhh1-dependent mRNAs were also derepressed on elimination of the decapping activators Pat1 or the combination of Edc3 and Scd6 (Figure 2A-B), suggesting that these four factors act together to stimulate decapping, such that eliminating any one is sufficient to stabilize the whole group of transcripts. One way to account for this behavior is to propose that the autoinhibition of Dcp2 activity exerted by different modules in its C-terminal IDR must be overcome simultaneously by independent interactions with each of these decapping activators. In this view, an active decapping complex comprised of the Dcp1/Dcp2 holoenzyme and Dhh1, Pat1, and Edc3 or Scd6 would be assembled on these mRNAs to accelerate decapping and degradation in WT cells (Figure 8 – figure supplement 1, *left*). This interpretation is consistent with recent findings indicating that repression of certain Dhh1-target mRNAs is mediated by Edc3 and Scd6 acting interchangeably to recruit Dhh1 to the same segment of the Dcp2 IDR, and with yeast two-hybrid data indicating that Dcp2 can interact with multiple decapping activators bound simultaneously to different modules in the IDR (He et al. 2022). The remaining one-half of the mRNAs appears to be degraded independently of Dhh1, Pat1, and Edc3 or Scd6. Interestingly, most of these transcripts are derepressed in mutants lacking Upf proteins to the same extent as in *dcp2*Δ cells, suggesting that most of them belong to the cohort of natural NMD substrates defined previously (Celik et al. 2017). Presumably, on such transcripts, aberrant splicing or transcription initiation, or non-canonical translation initiation or elongation, leads to out-of-frame translation and premature termination, resulting in activation of Dcp1/Dcp2 by Upf proteins (Figure 8 – figure supplement 1, *right*).

As would be expected if their abundance is controlled by decapping, Dcp2-repressed transcripts exhibit highly similar derepression in response to *dcp2* point mutations that inactivate Dcp2 catalytic activity compared to the *dcp2*Δ mutation (Figure 1D). Moreover, combining CAGE sequencing with RNA-Seq revealed that these mRNAs also exhibit greater than average proportions of uncapped molecules in WT cells, regardless of whether they are dependent or independent of Dhh1 for Dcp2 control over their abundance (Figure 3D). The accumulation of such uncapped degradation intermediates presumably reflects the fact that 5’-3’ degradation by Xrn1 is frequently delayed, eg. by the presence of elongating ribosomes that initiated translation prior to decapping (Pelechano et al. 2015). Hence, mRNAs preferentially targeted for degradation by decapping should exhibit a greater than average proportion of uncapped transcripts in WT cells, which is eliminated by deleting *DCP2*, just as we observed for both the Dhh1-dependent and -independent groups of mRNAs repressed by Dcp2 (Figure 3E).

Our parallel analysis of spike-in normalized Pol II occupancies in chromatin by Rpb1 ChIP-Seq and mRNA abundance by ERCC-normalized RNA-Seq revealed that the great majority of transcripts derepressed in *dcp2*Δ cells exhibit reduced, rather than elevated Pol II occupancies in the cognate CDSs (Figure 3G), confirming that decreased mRNA turnover rather than elevated transcription underlies their increased mRNA levels. An exception to this generalization are the iESR genes, especially those activated by transcription factor Msn2, which exhibit significantly elevated Rpb1 occupancies in *dcp2*Δ cells. Even for these Msn2-activated iESR genes, however, the increases in mRNA levels are much greater than the increases in Rpb1 occupancy conferred by *dcp2*Δ (3.71-fold vs. 1.47-fold increases in median values; cf. col.3 in Figure 8 – figure supplement 2A vs. Figure 3 – figure supplement 4B), suggesting substantial contributions from reduced decapping/degradation in addition to increased transcription. Supporting this, iESR transcripts, whether Msn2-activated or not, exhibit greater than average proportions of uncapped mRNAs (lower C/T ratios) in WT cells that are reversed by *dcp2*Δ (Figure 8 – figure supplement 2B, cols. 3-4 vs. 1), indicating preferential targeting by Dcp2 for decapping/decay; whereas most of the 199 Msn2 target genes do not show such evidence for enhanced decapping (Figure 8 – figure supplement 2B, col. 5 vs. 1). The fact that the Msn2-targeted iESR genes show greater increases both in mRNA abundance and Rpb1 occupancy compared to non-Msn2-targeted iESR genes in *dcp2*Δ cells (Figure 8 – figure supplement 2A and Figure 3 – figure supplement 4B, cols. 3-4 vs. 1) provides further evidence that a combination of Msn2-dependent transcriptional activation in response to slow cell growth and eliminating Dcp2-mediated mRNA decay underlies derepression of this subset of iESR genes in *dcp2*Δ cells.

The mRNAs dependent on Dhh1 for Dcp2-repression of abundance generally exhibit average translational efficiencies (TEs) in WT cells and also average CDS lengths, mRNA abundance, and mRNA stabilities (Figure 2D & F-H). The association of these properties with efficient translation (Thompson et al. 2016; Weinberg et al. 2016) seems inconsistent with the possibility that their enhanced decapping and degradation results from low rates of translation initiation with attendant greater access of the cap structure to Dcp1/Dcp2. These mRNAs also exhibit an average proportion of non-preferred codons (Figure 2E), suggesting that pausing during translation elongation is not a key driver of their enhanced decapping and degradation. As discussed further below, a third possibility is that Dcp1/Dcp2 are recruited to these mRNAs by one or more RNA binding proteins that interact with specific sequences in the 3’UTRs of these transcripts. The Dhh1-independent group of mRNAs, by contrast, tend to have lower than average TEs and codon optimality (Figure 2D-E), as noted previously for NMD substrates (Celik et al. 2017). Interestingly, this latter group is enriched for transcripts involved in DNA repair or recombination (Figure 7 – figure supplement 1D), distinct from the cellular functions involving Dhh1-dependent mRNAs, which include stress responses, metabolism of energy reserves, and the TCA cycle (Figure 7 – figure supplement 1C).

Notwithstanding the established function of Dcp2 in targeting mRNAs for degradation by decapping, we observed widespread changes in translational efficiencies in *dcp2*Δ cells (Figure 4A-B). The increased ribosome density (higher RPFs per mRNA) measured by ribosome profiling for the Dcp2-translationally repressed mRNAs was generally associated with increased steady-state protein abundance measured by TMT-MS/MS analysis in *dcp2*Δ vs. WT cells (Figure 4C), which was supported for particular mRNAs by increased expression of *nLUC* reporters constructed for the corresponding genes (Figure 4D). As a group, the Dcp2-translationally repressed mRNAs do not appear to be preferentially targeted by Dcp2 for mRNA turnover, as they exhibit decreased rather than increased relative abundance in *dcp2*Δ vs. WT cells in comparison to all mRNAs (Figure 5A).

We considered the possibility that the increased TEs conferred by *dcp2*Δ arise from enhanced decapping of these mRNAs and attendant accumulation of uncapped intermediates with a low rate of translation initiation in WT cells resulting from their inability to bind eIF4F. By abolishing decapping, *dcp2*Δ would eliminate this pool of low-TE intermediates and thereby increase the TEs relative to WT cells calculated for the total pools of such mRNAs (Figure 5B). At odds with this model, however, the Dcp2-translationally repressed mRNAs exhibit somewhat higher, not lower, than average proportions of capped mRNAs in WT cells (Figure 5C), inconsistent with translational repression by decapping.

Searching for a different mechanism, we noted that the mRNAs translationally repressed by Dcp2 tend to be efficiently translated in WT cells, showing higher than average calculated TEs, shorter than average CDS lengths, lower than average proportions of poor codons, and greater than average mRNA stability and abundance (Figures 4E-F & Figure 4 – figure supplement 2A-C). Two notable features of *dcp2*Δ cells suggested that the TE increases observed for most such mRNAs could arise indirectly from increased competition for limiting ribosomes in cells lacking Dcp2. First, *dcp2*Δ cells exhibit ∼30% lower than WT levels of mature ribosomes (Figure 5F-G), which probably reflects reduced expression of the rESR transcripts—highly enriched for mRNAs encoding ribosomal proteins or ribosome biogenesis factors (Gasch et al. 2000) (Figure 5 – figure supplement 1C & E). Although the RPG transcripts show increased TEs in *dcp2*Δ cells, this effect is outweighed by their reduced transcript abundance, to yield a net decrease in translation levels (RPF occupancies) in *dcp2*Δ vs. WT cells (Figure 5 – figure supplement 1E), which is consistent with the reduced ribosome content in *dcp2*Δ cells (Figure 5G). Second, *dcp2*Δ cells exhibit ∼24% higher than WT levels of total mRNA, determined by RNA-Seq analysis with spike-in normalization (Figure 3F), as might be expected from decreased decapping and mRNA turnover. It has been reported that reduced mRNA turnover in mutants lacking mRNA decay factors is buffered by decreased transcription (Sun et al. 2013), which likely limits the increase in mRNA levels we observed in *dcp2*Δ cells. However, the fact that all mRNAs should be capped in *dcp2*Δ cells implies that the increase in capped mRNA levels will be even larger than the 1.25-fold determined for all mRNAs. We reasoned that that the decrease in ribosome content coupled with the increase in capped mRNA abundance should increase competition among mRNAs for limiting 43S PICs, which in turn will favor the mRNAs that are most efficiently translated in WT cells (Figure 5E). Supporting this idea, which was first predicted by mathematical modeling (Lodish 1976), the TE changes conferred by *dcp2*Δ are strongly mimicked by two other conditions that limit 43S PIC assembly and also appear to increase competition among mRNAs to favor strongly translated mRNAs (Gaikwad et al. 2021): (i) phosphorylation of eIF2α in amino acid-starved cells and (ii) impairment of 40S recycling at stop codons by eliminating recycling factors Tma64/Tma20 (Figure 5 – figure supplement 1A-B). While other explanations are possible, this indirect mechanism is a plausible way to account for the bulk of translational programming conferred by eliminating the decapping enzyme.

Just as we found for changes in mRNA abundance, the alterations in TE conferred by *dcp2*Δ can occur either dependently or independently of Dhh1, with the minority, Dhh1-dependent group showing greater translational repression by Dcp2 compared to the Dhh1-independent group (Figure 6A-B). The Dhh1-independent group exhibits the key features associated with the indirect mechanism of translational repression involving increased competition for limiting PICs: efficient translation and greater than average proportions of capped transcripts in WT cells (Figure 6C-D). The ∼100 mRNAs belonging to the Dhh1-dependent group, by contrast, exhibits much lower than average TEs in WT (Figure 6C), which seems incompatible with the competition mechanism. The latter are also preferentially targeted by Dcp2 and Dhh1 for degradation (Figure 6 – figure supplement 1B-C) and exhibit higher than average proportions of uncapped degradation intermediates mRNAs in WT, which are reversed by *dcp2*Δ or *dhh1*Δ (Figure 6D-F).

An alternative mechanism to account for the ∼100 Dhh1-dependent Dcp2-translationally repressed mRNAs would be to propose that Dcp2 and Dhh1 sequester them in P-bodies, where translationally silent mRNAs can be enriched (Luo et al. 2018), without stimulating their decapping. In this model, association with Dcp2/Dhh1 would be sufficient for translational repression without a requirement for Dcp2 catalytic activity. At odds with this model, however, our ribosome profiling analysis of the *dcp2-EE* catalytic mutant indicates highly similar TE changes conferred by this point mutation and by the *dcp2*Δ deletion (Figure 5 – figure supplement 1D). We cannot rule out the scenario that *dcp2-EE* impairs its association with mRNAs to reduce P body formation; nor the possibility that the increased total mRNA abundance conferred by *dcp2-EE* would interfere with the ability of Dcp2 to sequester this specific subset of translationally-repressed mRNAs in P bodies. Additional work is required to elucidate the mechanism of Dhh1-dependent translational repression by Dcp2.

*S. cerevisiae* preferentially derives energy from glucose via fermentation, producing ethanol, and represses aerobic respiration in glucose-rich medium such as YPD. Importantly, we found that Dcp2-translationally repressed mRNAs are enriched for transcripts encoding mitochondrial proteins involved in respiration, including a majority of those encoding components of the electron transport chain (ETC) and mitochondrial ATPase; and that 8 mRNAs encoding TCA cycle enzymes and three encoding ETC components are also repressed in abundance by Dcp2 (Figure 7 – figure supplement 1A-B). Indeed, a group of ∼50 such mitochondrial proteins directly involved in respiration shows increased median ribosome occupancies (Figure 7B) and steady-state protein abundance (Figure 7 – figure supplement 2K) in *dcp2*Δ vs. WT cells, which we confirmed for several proteins by Western blotting (Figure 7C). These findings suggest that oxidative phosphorylation is up-regulated in the *dcp2*Δ mutant on YPD medium, which was supported by our demonstration of increased mitochondrial membrane potential in *dcp2*Δ vs. WT cells (Figure 7D).

*S. cerevisiae* also preferentially utilizes glucose and represses catabolism of alternative carbon sources on glucose-replete medium, in part by deactivating transcriptional activators Adr1 and Cat8 (Zaman et al. 2008). We observed increased mRNA abundance and translation of *ADR1, CAT8* and a group of ∼100 glucose-repressed or Adr1/Cat8 target genes (Figure 7 – figure supplement 1C-D & Figure 7E) in *dcp2*Δ cells. Similarly, a group of ∼36 genes that are transcriptionally repressed in the presence of preferred nitrogen sources (NCR genes) showed elevated median mRNA levels and translation in *dcp2*Δ cells on YPD medium (Figure 7F). These findings suggest that Dcp2 post-transcriptionally enhances fermentation of glucose and catabolism of preferred nitrogen sources on rich medium. Consistent with this, Dcp2 helps to suppress the generation of amino acids from proteins via autophagy on rich medium by repressing levels of certain *ATG* mRNAs encoding proteins required specifically for this process (Hu et al. 2015), which we confirmed here (Figure 7 – figure supplement 2F-G). Interestingly, *dcp2*Δ also derepressed the abundance of multiple cell adhesion genes and conferred increased cell filamentation and invasive growth on rich medium (Figure 7 – figure supplement 2H & Figure 7G-I), processes that are normally suppressed when nutrients are plentiful (Roberts and Fink 1994). The same phenotypes are also displayed to a lesser degree by elimination of Pat1 (Figure 7 – figure supplement 2I-J). If filamentation and invasive growth are regarded as strategies for nutrient foraging, then the repression of these processes by Dcp2 and Pat1 on rich medium supports the model that mRNA decapping and attendant translational repression by Dcp1/Dcp2 participates in repressing multiple pathways that are dispensable in cells growing on nutrient-rich medium.

Previously, it was shown that the yeast Pumilio family RNA binding protein Puf3 coordinately represses the expression of multiple mRNAs encoding mitochondrial proteins that support the synthesis or assembly of the complexes that carry out oxidative phosphorylation, during fermentative growth on glucose where respiration is repressed. The Puf3-targeted mRNAs include many protein components of mitochondrial ribosomes needed for translation of the small number of mitochondrially-encoded proteins, and other factors involved in import, folding, or maturation of nuclear-encoded mitochondrial proteins, but only two proteins (Atp1 and Atp18) directly involved in oxidative phosphorylation (Lapointe et al. 2018). It appears that Puf3 reduces the abundance of these target mRNAs by enhancing their deadenylation, decapping and degradation in glucose-grown cells (Olivas and Parker 2000; Miller et al. 2014). We found that Dcp2 represses a sizeable group of 52 transcripts directly involved in oxidative phosphorylation, primarily at the level of translation (Figure 7B) and in a manner that appears to conform to the indirect ribosome competition model for translational reprograming (Figure 5E), as they exhibit a greater than average median TE in WT cells. About 10 of these transcripts are also repressed by Dcp2 at the level of mRNA abundance, but the two Puf3 targets *ATP1* and *ATP18* are not among them. In fact, *ATP1* is the only Puf3 target among this group of 52 transcripts whose translation is derepressed by *dcp2*Δ. It seems unlikely therefore that Puf3 plays a direct role in regulating the abundance or translation of these mRNAs by Dcp2. Interrogating the mitochondrial protein mRNAs that do bind Puf3, and focusing on a subset of 91 that also exhibit increased expression of the encoded proteins in *puf3*Δ cells (dubbed Puf3 “cis” targets by Lapointe et al.), we found derepression of translation at the TE but not mRNA level in *dcp2*Δ (Figure 8 – figure supplement 3). Further, *dcp2*Δ conferred derepressed translation (increased RPF occupancies) by >1.5-fold for 19 of them, but it conferred a comparable increase in mRNA abundance for only five (*IMG2, SDH5, MPM1, CIR1, MSC6*), with the increased translation of the remaining 14 mRNAs resulting from lesser increases in both mRNA levels and TE. It is possible that Puf3 is involved in the Dcp2-mediated repression of mRNA abundance of the aforementioned five Puf3 target mRNAs. In addition to repressing transcript abundance in fermentative growth, Puf3 functions in localizing its direct mRNA targets to mitochondria (Saint-Georges et al. 2008), including at least ∼3/4^th^ of the 91 Puf3 “cis” transcripts described above. Although this Puf3 function is thought to enhance co-translational import of mitochondrial proteins during respiratory growth, it could be proposed that Puf3 represses the translation as well as stability of its target mRNAs during fermentative growth. It is possible therefore that Puf3 also plays a role in repressing the TEs of the subset of the aforementioned 19 transcripts among the group of 91 “cis” targets whose translation is derepressed by *dcp2*Δ; although these mRNAs are well-translated in WT cells and could be translationally repressed by the ribosome competition mechanism instead. It seems plausible that other RNA binding proteins besides Puf3 mediate the derepression of mRNA levels by *dcp2*Δ for many of the ∼1330 other transcripts in the mRNA_up_*dcp2*Δ group or members of the 100 mRNAs translationally repressed by Dcp2 in a manner requiring Dhh1, which were identified in our study.

## MATERIALS AND METHODS

### Strains, plasmids, and culture conditions

Yeast strains, plasmids, and primers used in the study are listed in Supplementary Tables 1 to 3, respectively. Yeast strain VAK022 harboring *dcp2-E149Q,E153Q* (*dcp2-EE*) was constructed in two steps. First, mutations G to C at positions 445 and 457 were introduced into *DCP2* in plasmid pQZ145 (Zeidan et al. 2018) using primers AKV005/AKV006 and the Quick-change Site-Directed mutagenesis kit (Agilent, 200519), generating plasmid pAV008 containing *dcp2-E149Q,E153Q*. Next, a 3.3-kb fragment containing *dcp2-E149Q,E153Q* with 400 bp upstream (have restriction site for Pf1N1 in the upstream region) of the start codon was PCR-amplified using primers AKV088/AKV090, and the resultant fragment was inserted between the BamHI and SacI sites of integrative vector YIplac211 to produce plasmid pAKV013. The DNA fragment generated by digestion of pAKV013 with Pf1N1 was used to transform WT strain W303 to Ura^+^. Finally, strain VAK022 was obtained from one such Ura^+^ transformant by selecting for loss of the *URA3* marker via homologous recombination by counter-selection on medium containing 5-fluoro-orotic acid. The replacement of *DCP2* with *dcp2-E149Q, E153Q* was verified by sequencing the PCR product obtained from chromosomal DNA amplified with the primer pairs AKV010/AKV015.

The 13 reporter plasmids containing *nLUC* reporters listed in Supplementary Table 2 were constructed to fuse the nanoLUC coding sequences (codon-optimized for *S. cerevisiae*, Supplementary Table 5), preceded by the GGG glycine codon, to the final codon of the complete CDS of each gene, preserving the native stop codon, 3’UTR sequences, and segment of 3’-noncoding sequences of the gene, as well as a segment of 5’-noncoding sequences including the native promoter and 5’UTR sequences, of the gene of interest (fragment lengths of 5’ and 3’ region is listed in Supplementary Table 4), by the following three-step procedure. First, DNA fragments were synthesized containing a SmaI site, the nLUC CDS (including the ATG and stop codon), the native stop codon and 3’ non-coding sequences for each gene of interest, and an EcoRI site, and inserted between the SmaI and EcoRI sites of pRS316 to produce an intermediate plasmid for each gene of interest. Second, PCR-amplification from WT yeast genomic DNA was conducted to generate a fragment for each gene of interest containing 20 nt of pRS316 adjacent to the SmaI site, the CCC nucleotides of the SmaI site, the 5’-noncoding region, 5’UTR, and CDS of the relevant gene (excluding the stop codon), the CCC complement of the GGG glycine codon, and the first 20nt of the *nLUC* CDS. Third, Gibson assembly was used to insert the PCR-amplified fragments from the second step at the SmaI sites of the intermediate plasmids, using 2x ExSembly Cloning Master mix (LifeSct LLC, M0005) and following the vender’s instructions except that a 15 min incubation at 25°C was included prior to the 37°C incubation to allow for optimal SmaI digestion.

Plasmid pNG158 contain the *PAT1* CDS, with 500 bp upstream and 192 bp downstream, on a 3083 bp fragment amplified by PCR from yeast genomic DNA, and inserted between the SalI and XmaI sites of pRS315, respectively. The plasmids were constructed by NEBuilder HiFi DNA assembly (New England Biolabs) according to the manufacturer’s protocol.

Unless mentioned otherwise, strains were cultured in YPD medium to mid-exponential growth phase (OD_600_ ∼0.6) before harvesting.

### Ribosome profiling and parallel RNA sequencing

Ribosome profiling and RNA-seq analysis were conducted in parallel, essentially as described previously (McGlincy and Ingolia 2017), using isogenic strains W303 (WT), CFY1016 (*dcp2*Δ), and VAK023 (*dcp2-EE*) with two biological replicates performed for each genotype. Cells were harvested by vacuum filtration and flash-frozen in liquid nitrogen. Cells were lysed in a freezer mill in the presence of lysis buffer (20 mM Tris (pH 8), 140 mM KCl, 1.5 mM MgCl2, 1% Triton X-100) supplemented with 500 mg/mL cycloheximide. Lysates were cleared by centrifugation at 3000 x g for 5 min at 4°C, and the resulting supernatant was subjected to centrifugation at 15,000 x g for 10 min at 4°C, flash-frozen in liquid nitrogen, and stored at -80°C.

For preparation of libraries of ribosome-protected mRNA fragments (RPFs), 50 A_260_ units of cell lysates were digested with 450 U of RNase I (Ambion, AM2294) for 1 h at room temperature (RT, ∼25°C) on a Thermomixer at 700 rpm, and extracts were resolved on 10–50% sucrose gradients by centrifugation for 160 min at 39,000 rpm, 4°C in a Beckman SW41Ti rotor. Gradients were fractionated at 0.75 ml/min with continuous monitoring of A_260_ values using a Biocomp Instruments Gradient Station. RNA was purified from the 80S monosome fractions using an RNA Clean and Concentrator kit (Zymo, R1018), resolved by electrophoresis on a 15% TBE-Urea gel, and 25-34 nt fragments isolated from the gel were dephosphorylated using T4 Polynucleotide kinase (New England Biolabs, M0201L). For each library sample, barcoded 5’ -pre-adenylated linkers were added to the 3’ ends of footprints using T4 Rnl2(tr) K227Q (New England Biolabs, M0351S), and excess unligated linker was removed using 10 U/µl 5’ deadenylase/RecJ exonuclease (Epicentre, RJ411250), followed by pooling and purification of ligated footprints using an Oligo Clean and Concentrator column (Zymo Research, D4060). Ribosomal RNA contamination was removed by biotinylated primers (McGlincy and Ingolia 2017), followed by reverse transcription using Protoscript II (New England Biolabs, M0368L) and circularization of cDNA using CircLigase ssDNA Ligase (Epicenter, CL4111K). Each pooled library was PCR amplified using Phusion polymerase (F-530) (New England Biolabs, M0530S). Quality of the libraries was assessed with a Bioanalyzer using the High Sensitivity DNA Kit (Agilent, 5067–4626) and quantified by Qubit. Single-end 50 bp sequencing was done on an Illumina HiSeq system at the NHLBI DNA Sequencing and Genomics Core at NIH (Bethesda, MD).

For RNA-Seq library preparation, total RNA was extracted and purified from aliquots of the same snapped-frozen cells described above using the Zymo RNA Clean and Concentrator kit. Five mg of total RNA was randomly fragmented by incubating with Ambion Fragmentation Reagent (Ambion, AM8740) at 70°C for 12 min, the reaction was stopped by adding Stop Solution (Ambion, AM8740) and precipitated by adding one part of isopropanol followed by precipitation with 70% ethanol. Fragment size selection, library generation, and sequencing were carried out using the same protocol described above for RPF library preparation, except that Ribo-Zero Gold rRNA Removal Kit (Illumina, MRZ11124C) was employed to remove rRNA after linker-ligation.

As described earlier (Martin-Marcos et al. 2017), Illumina sequencing reads were trimmed to remove the constant adapter sequence, mixed sample sequences were separated by the sample barcodes followed by removal of PCR duplicates using a custom Python (3.7) script. The sequences aligned to yeast non-coding RNAs were removed using bowtie (Langmead et al., 2009) and non-rRNA reads (unaligned reads) were then mapped to the *S. cerevisiae* genome (R64-1-1 S288C SacCer3 Genome Assembly) using TopHat (Trapnell et al., 2009). Only uniquely mapped reads from the final genomic alignment were used for subsequent analyses. Statistical analysis of changes in mRNA, RPFs, or TE values between two replicates each of any two strains being compared was conducted using DESeq2 (Love et al. 2014) excluding any genes with less than 10 total mRNA reads in the 4 samples (of two replicates each) combined. DESeq2 is well-suited to identifying changes in mRNA or RPF expression, or TEs, with very low incidence of false positives using results from only two highly correlated biological replicates for each of the strains/conditions being compared (Zhang et al. 2014; Lamarre et al. 2018). The R script employed for DESeq2 analysis of TE changes can be found on Github (https://github.com/hzhanghenry/RiboProR; (Kim et al. 2019). Wiggle files were generated as described previously (Zeidan et al. 2018) and visualized using Integrative Genomics Viewer (IGV 2.4.14, http://software.broadinstitute.org/software/igv/) (Robinson et al. 2011). Wiggle tracks shown are normalized according to the total number of mapped reads.

### Parallel CAGE and RNA sequencing

Total RNA was prepared by hot-phenol extraction (Schmitt et al. 1990) from two biological replicates of each strain. RNA integrity was determined using the Agilent RNA 6000 Nano kit (5067-1511) and the concentrations were measured by Nanodrop spectroscopy. CAGE libraries were constructed and sequenced following the nAnT-iCAGE protocol (Murata et al. 2014) by K.K. DNAFORM of Japan. Briefly, cDNAs were transcribed to the 5’ ends of capped RNAs, ligated at the 5’ and 3’ ends with barcoded linkers, followed by 2^nd^ strand synthesis, and the resulting DNA libraries were sequenced using the Illumina NextSeq500 (Single-end, 75 bp reads) platform. Between 20-25 million mapped CAGE tags were obtained for each sample (Figure 3 – figure supplement 1 – source data 1).

The sequenced CAGE reads of each sample were aligned to the reference genome of *S. cerevisiae* S288C (Assembly version: sacCer3) using HISAT2 (Kim et al. 2019). For read alignment, we disabled the soft clipping option in HISAT2 by using “--no-softclip” to avoid false-positive transcription start sites (TSSs). CAGE reads mapped to rRNA genes were identified using rRNAdust (http://fantom.gsc.riken.jp/5/sstar/Protocols:rRNAdust), and were excluded from subsequent TSS analyses (Figure 3 – figure supplement 1 – source data 1).

TSS identification, inference of TSS clusters (TC, representing putative core promoters), and assigning TCs to their downstream genes were carried out by using TSSr (Lu et al. 2021). CAGE reads with a mapping quality score (MAPQ) > 20 were considered uniquely mapped reads, which were used for subsequent analyses. CAGE signals of biological replicates were then merged as a single sample. The transcription abundance of each TSS was quantified as the numbers of CAGE tags/reads supporting the TSS per million mapped reads (TPM). Only TSSs with TPM ≥ 0.1 were used to infer TSS clusters (TCs), representing putative core promoters.

The “peakclu” method (Lu et al. 2021) was used to infer TCs for each sample, with the following options “peakDistance=50, extensionDistance=25, localThreshold = 0.01”. A set of Consensus TCs of all samples were generated by using the “consensusCluster” function in TSSr with an option of “dis = 100”. Consensus TCs were then assigned to their downstream genes if they are within 1000 bp upstream and 50 bp downstream of the start codon of annotated ORFs. The TPM value of a consensus TC in a sample is the sum of TPM values of all TSSs within its range. The TPM value of a gene was calculated as the sum of all consensus TC assigned to the gene.

RNA sequencing libraries were produced in parallel from the same RNA samples subjected to CAGE sequencing by the NHLBI DNA sequencing Core at NIH (Bethesda, MD) using the TruSeq Stranded mRNA Library Prep Kit (Illumina, Paired-end 50 bp reads) and sequenced using the NovaSeq6000 Illumina platform. Sequencing reads were mapped to the S288C genome (R64-1-1 S288C SacCer3) using STAR aligner (Dobin et al. 2013) and PCR duplicates were removed by Samtools (Figure 3 – figure supplement 1 – source data 1).

Identification of differentially expressed genes in total RNA (△mRNA_T) or capped RNA (△mRNA_C) in *dcp2*Δ vs. WT cells was conducted by DESeq2 analysis (Love et al. 2014) using raw read counts from the RNA-seq or CAGE sequencing experiments conducted in parallel on the same RNA samples. To calculate C/T ratios, RNA-Seq reads assigned to each gene were normalized as transcripts per million reads (TPM) by dividing the read counts by the length of each gene in kilobases (reads per kilobase, RPK), summing all the RPK values in a sample and normalizing, to generate “per million” scaling factors. Further, RPK values were normalized by the “per million” scaling factor to give TPM. Because a single read/tag is generated for each transcript in CAGE, its TPM (tags per million mapped tags) is equivalent to the TPM value obtained from RNA-seq, allowing comparisons of the two types of TPM values. The C/T ratio of each gene was calculated by dividing CAGE TPMs by RNA-Seq TPMs in each strain after first removing all genes with zero CAGE reads in either the *dcp2*Δ or WT samples being compared. In interrogating C/T ratios for the groups of mRNAs derepressed or repressed by *dcp2*Δ, the same two groups defined using the RNA-Seq data obtained in parallel with ribosome profiling were examined to allow changes in mRNA abundance, C/T ratios, ribosome occupancies, and TEs to be compared for the same two sets of mRNAs. The conclusions reached regarding changes in C/T ratios were not altered if the groups were defined using the RNA-seq data obtained in parallel with CAGE-sequencing instead.

### RNA sequencing with spike-in normalization

ERCC ExFold RNA spike-In Mixes (Ambion, part no 4456739) consisting of spike-In Mix I and spike-In Mix II were equally added to WT and *dcp2*Δ total RNA (2.4 ul of 1:100 fold diluted spike-In to 1.2 µg of total RNA), respectively, according to the manufacturer instructions. STAR software was used to align mapped sequencing reads, including reads from the spike-In RNAs, to the S288C genome or ERCC RNA sequences. Reads obtained from the 23 spike-in transcripts belonging to subgroup B (present in equal concentrations in Mix I and Mix II) were used to calculate size factors for each library and reads corresponding to yeast genes were normalized by the size factors. DESeq2 was further employed to calculate the differential expression between strains by setting the size factor to unity.

### ChIP-Seq and data analysis

WT and *dcp2*Δ strains were cultured in triplicate in YPD medium to A_600_ of 0.6–0.8 and treated with formaldehyde as previously described (Qiu et al. 2016). ChIP-Seq was conducted as described (Qiu et al. 2016) using monoclonal antibody against Rpb1 (8WG16, Biolegend, 664906). DNA libraries for Illumina paired-end sequencing were prepared using the DNA Library Prep Kit for Illumina from New England Biolabs (E7370L). Paired-end sequencing (50 nt from each end) was conducted by the DNA Sequencing and Genomics core facility of the NHLBI, NIH. Sequence data were aligned to the SacCer3 version of the genome sequence using Bowtie2 (Langmead et al. 2009) with parameters -X 1000 -very-sensitive, to map sequences up to 1 kb with maximum accuracy. PCR duplicates from ChIP-Seq data were removed using the samtools rmdup package. Numbers of aligned paired reads from each ChIP-Seq experiment are summarized in Figure 3 – source data 3. Raw genome-wide occupancy profiles for Rpb1 were computed using the coverage function in R and relative occupancies were obtained by normalizing each profile to the average occupancy obtained for the relevant chromosome (https://github.com/rchereji/bamR). To visualize specific loci, BigWig files of samples were loaded in the Integrative Genomics Viewer (IGV) (Robinson et al. 2011).

For spike-in normalization of Rpb1 ChIP-Seq data, identical aliquots of *S. pombe* chromatin were added to each *S. cerevisiae* chromatin sample being analyzed in parallel, corresponding to 10% of the DNA in the *S. cerevisiae* chromatin samples, prior to immunoprecipitating with Rpb1. As described fully in Figure 3 – source data 3, a normalization factor for each sample was calculated by dividing the average number of total *S. pombe* reads obtained across all samples by the total *S.pombe* reads obtained for that sample. The observed reads mapping to the *S. cerevisiae* genome were multiplied by the normalization factor to yield the spike-in normalized reads for that sample. Raw genome wide occupancy profiles for Rpb1 were computed using the coverage function in R, wherein each profile was set with the same total “OCC” to allow the comparison between WT and *dcp2*Δ, using the custom R script (https://github.com/hzhanghenry/OccProR).

### qRT-PCR analysis of mRNA abundance

Total RNA was isolated by hot-phenol extraction as previously described (Schmitt et al. 1990) and the concentration was determined using the Nanodrop ND-1000 spectrophotometer (ThermoFisher). Ten µg of total RNA was treated with DNase I (Roche, 4716728001) and 40 picograms of Luciferase Control RNA (Promega L4561) was added to 1 ug of DNase I-treated total RNA and subjected to cDNA synthesis using a Superscript III First-Strand synthesis kit (Invitrogen 18080051). qRT-PCR was carried out using 10-fold diluted cDNA and Brilliant II SYBR Green qPCR Master Mix (Agilent, 600828) with the appropriate primer pairs (listed in Supplementary Table 3) at 200 nM. Expression of each transcript was normalized to that of the luciferase spike-in RNA from at least two biological replicates.

### TMT-MS/MS analysis

Three biological replicates of WT and *dcp2*Δ were cultured in YPD medium and harvested by centrifugation for 5 min at 3000 x *g*. Whole cell extracts (WCEs) were prepared using 8M Urea in 25 mM triethylammonium-bicarbonate (TEAB; Thermo Scientific, 90114) by washing the pellets once with the same buffer and vortexing with glass beads in the cold room for 2 min with intermittent cooling on ice water. Lysates were clarified by centrifugation at 13,000 x *g* for 30 min and the protein quality was assessed following SDS-PAGE using GelCode™ Blue Stain (Thermo Scientific, 24592) and quantified using Pierce™ BCA Protein Assay Kit (Thermo Scientific, 23225). Sample preparation and TMT-MS/MS (Zecha et al. 2019) was performed by the NHLBI Proteomics Core at NIH (Bethesda, MD). Briefly, 100 µg of WCEs was incubated for 1 h at 37°C with freshly prepared DTT (20 mM final) to reduce disulfide bridges. Alkylation was performed at RT for 1 h with freshly made 50 mM iodoacetamide (50 mM, final) in 25 mM ammonium bicarbonate and the reaction was quenched by adding DTT (50 mM, final). Lysates were diluted 10-fold with 25 mM ammonium bicarbonate and digested with 3 µg of trypsin (Promega, v5111) overnight at 37°C. Digests were acidified by adding formic acid (1%, final) and desalted with Waters Oasis HLB 1cc columns. Peptides were eluted from desalted samples with 1 ml of buffer E (0.1% formic acid in 50% acetonitrile) and dried in a SpeedVac. Samples were labelled with TMT reagents for multiplexing (TMT10plex label reagent set, Thermo Scientific) according to the manufacturer’s instructions. Briefly, resuspended TMT reagent was added to each sample, incubated for 1 h at RT and the reaction quenched with 8 µl of 5% hydroxylamine for 15 min. Equal amounts of each sample were combined and the pooled sample was dried in a SpeedVac. To increase the protein coverage, each set of pooled TMT samples was separated into 24 fractions using basic reverse phase liquid chromatography (bRPLC). Quantification of TMT-labelled peptides was conducted on an LTQ Orbitrap Lumos-based nanoLCMS system (Thermo Scientific) with a 2 h gradient at 120k resolution for MS1 and 50K for MS2 at 38% HCD energy.

Raw data was processed using Proteome Discoverer 2.4 (Thermo Scientific) and the MS2 spectra were searched in the SwissProt Yeast database (https://www.uniprot.org/proteomes/UP000002311) using the SEQUEST search engine (Eng et al. 1994). Peptide spectral matches (PSM) were validated using Percolator based on q-values at a 1% FDR (Brosch et al. 2009) (http://www.sanger.ac.uk/Software/analysis/MascotPercolator/). Relative abundance of each peptide in a strain is measured by normalizing to the total abundance of that peptide coming from all the strains used in the study. We determined the protein-level fold changes based on the median of peptide-level fold changes from the Proteome Discoverer-produced abundances.

### Measuring *nLUC* reporter expression

Nano-luciferase was assayed in WCEs as previously described (Masser et al. 2016). Briefly, WT and *dcp2*Δ transformants harboring the appropriate *nLUC* reporter plasmids (Supplementary Table 2) or empty vector pRS316 were cultured in synthetic complete medium lacking uracil (SC-Ura) to OD_600_ of ∼1.2. Cells were collected by centrifugation at 3000 x g and resuspended in 1X PBS lysis buffer containing 1mM PMSF and protease inhibitor cocktail (Roche, 5056489001). WCEs were prepared by vortexing with glass beads, and clarified by centrifugation at 13000 x g for 30 min at 4°C. Nano-Glo substrate (Promega, N1120) was diluted 1:50 with the supplied lysis buffer and mixed with 10 µl of WCE in a white 96-well plate. Bioluminescence was determined immediately using a Centro Microplate Luminometer (Berthold), and light units were normalized by the total protein concentrations of the corresponding WCEs determined using the Bradford reagent (BioRad, 5000006).

### Polysome profiling to measure ribosome content

Three biological replicates of WT and *dcp2*Δ were cultured in 300 ml of YPD medium at 30°C to OD_600_ of 1.2 to 1.5 and quick-chilled by pouring into centrifuge tubes filled with ice. After collecting the cells by centrifugation for 10 min at 7000 x g, the cell pellets were resuspended in an equal volume of Buffer A (20 mM Tris-HCl [pH 7.5], 50 mM NaCl, 1 mM DTT, 200 μM PMSF, 0.15 µM Aprotinin, 1 µM Leupeptin, 0.1 µM Pepstatin A) and WCEs were prepared by vortexing with glass beads in the cold room, followed by two cycles of centrifugation for 10 min at 3,000 rpm and 15,000 rpm at 4°C, respectively. (Cycloheximide and MgCl_2_ were omitted from the lysis buffer in order to separate 80S ribosomes into 40S and 60S subunits.) Equal volumes of cleared lysate from WT and *dcp2*Δ cultures were resolved on 5–47% (w/w) sucrose gradients by centrifugation at 39,000 rpm for 3 h at 4°C in a Beckman SW41Ti rotor. Gradient fractions were scanned at 260 nm using a gradient fractionator (Bio-comp Instruments, Triax), and the area under the 40S and 60S peaks were quantified using ImageJ software. To estimate the ribosomal content per cell volume, the combined areas under the 40S and 60S peaks were normalized by the OD_600_ values of the starting cultures.

### Western-blot analysis

For Western analysis of respiratory proteins, WCEs were prepared by trichloroacetic acid (TCA) extraction as previously described (Reid and Schatz 1982) and immunoblot analysis was conducted as described previously (Nanda et al. 2009). After electroblotting to PVDF membranes (Millipore, IPVH00010), membranes were probed with antibodies against Aco1, Atp20, Cox14, Pet10 (a kind gift from Dr. Nikolaus Pfanner), Idh1 (Abnova, PAB19472), and GAPDH (Proteintech, 60004). Secondary antibodies employed were HRP-conjugated anti-rabbit (GE, NA9340V), anti-mouse IgG (GE, NA931V) and anti-goat IgG (Abnova, PAB29101). Detection was performed using enhanced chemiluminescence (ECL, Cytiva, RPN2109) and the Azure 200 gel imaging biosystem.

### Measuring mitochondrial membrane potential

Precultures were grown in SC-Ura (to select for the *URA3* plasmids) to OD_600_ of ∼3.0 and used to inoculate YPD medium at OD_600_ of 0.2. Cells were grown to OD_600_ of ∼0.6-0.8 and incubated with 500 nM TMRM for 1 h. Cells were washed once with distilled H_2_O and fluorescence was measured using flow cytometry (BD LSR II) at the Microscopy, Imaging & Cytometry Resources (MICR) Core at Wayne State University. Dye fluorescence is proportional to mitochondrial membrane potential. Median fluorescence intensity (MFI) of single cells was analyzed with the FlowJo software, and normalized to the OD_600_ of the cultures. Data presented are in arbitrary fluorescence units normalized to OD_600_ of the cultures. In control samples, 50 µM FCCP was added to cells to dissipate the membrane potential and provide a measure of non-specific background fluorescence.

### Plate-washing assay of invasive cell growth

The plate-washing assay was performed as described (Roberts and Fink 1994). Briefly, plates were incubated for 4d at 30°C and photographed before washing using the ChemiDoc XRS+ molecular imager (Bio-Rad) under the blot/chemicoloric setting with no filter. Small amounts of cells were excised from colonies using a toothpick and resuspended in water for microscopic examination. Plates were also photographed after being washed in a stream of water. Invasive growth was measured by the Image Lab 6.0.1 program (Bio-Rad, https://www.bio-rad.com/en-us/product/image-lab-software?ID=KRE6P5E8Z) using the round volume tool. Invasive growth levels were normalized for colony size and reported as the average of three independent replicates (Vandermeulen and Cullen 2020; Vandermeulen and Cullen 2022). Error represents the standard deviation. Significance was determined by Student’s t-test, p-value < 0.05.

### Data visualization and statistical analysis

Notched box-plots were constructed using a web-based tool at http://shiny.chemgrid.org/boxplotr/. In all such plots, the upper and lower boxes contain the 2^nd^ and 3^rd^ quartiles and the band gives the median. If the notches in two plots do not overlap, there is roughly 95% confidence that their medians are different. For some plots, the significance of differences in medians was assessed independently using the Mann-Whitney U test computed using the R Stats package in R. Scatterplots displaying correlations between sequencing read counts from biological replicates were created using the scatterplot function in Microsoft Excel. Spearman’s correlation analysis and the Student’s t-test were conducted using Microsoft Excel. Venn diagrams were generated using the web-based tool https://www.biovenn.nl/ and the significance of gene set overlaps in Venn diagrams was evaluated with the hypergeometric distribution using the web-based tool https://systems.crump.ucla.edu/hypergeometric/index.php Hierarchical clustering analysis of mRNA or TE changes in mutant vs. WT strains was conducted with the R heatmap.2 function from the R ‘gplots’ library, using the default hclust hierarchical clustering algorithm. Volcano plots were created using the web-based tool https://huygens.science.uva.nl/VolcaNoseR/. Gene ontology (GO) analysis was conducted using the web-based tool at http://funspec.med.utoronto.ca/.

### Data resources for mRNA features

Analyses of mRNA features for different gene sets were conducted using the following published compilations: 5’UTR and CDS lengths (Pelechano et al. 2013), WT mRNA steady-state amounts (in molecules per dry cellular weight - pgDW) (Lahtvee et al. 2017), WT mRNA half-lives (Chan et al. 2018), and stAI values (Radhakrishnan et al. 2016).

### Data availability

Ribosome profiling, RNA-Seq, ChIP-Seq and CAGE-Seq data discussed in this publication have been deposited in NCBI’s Gene Expression Omnibus and are accessible through GEO Series accession numbers GSE220578 (https://www.ncbi.nlm.nih.gov/geo/query/acc.cgi?acc=GSE220578) and GSE216831 (https://www.ncbi.nlm.nih.gov/geo/query/acc.cgi?acc=GSE216831). Previously published datasets used in the study can be found at <u>10.1371/journal.pgen.1008299</u> (Zeidan et al. 2018), <u>10.1261/rna.060541.116</u> (Celik et al. 2017), <u>10.1101/gr.209015.116</u> (Jungfleisch et al. 2017) https://doi.org/10.7554/eLife.34409 (He et al. 2018), and <u>10.1016/j.cell.2016.08.053</u> (Radhakrishnan et al. 2016).

## ACKNOWLEDGEMENTS

We thank Feng He, Allan Jacobson, and Bertrand Seraphin for generous gifts of yeast strains, and Nikolaus Pfanner for gifts of antibodies. We thank Yong Chen and Marjan Gucek of the NHLBI Proteomics Core for guidance on performing TMT-MS. We are grateful to Thomas Dever, Nicholas Guydosh and Jon Lorsch for many helpful discussions about data analysis and interpretation of results, and all members of the Hinnebusch, Lorsch, Dever, and Guydosh labs, for useful comments. This work was supported in part by the Intramural Program of the NIH. C.O. and M.L.G. were supported by NIH grant R01 HL117880; M.D.V. and P.J.C. by NIGMS grant GM098629; and X.N. and Z.L. by NSF grant 1951332 and the Saint Louis University 2022 President’s Research Fund.

## LEGEND FOR SOURCE FILES

**Figure 1 – source data 1**

RNA-Seq analysis showing log_2_ fold-changes in mRNA abundance by *dcp2*Δ vs. WT for all transcripts, mRNA_up_*dcp2*Δ and mRNA_dn_*dcp2*Δ groups (Figure 1A-B).

**Figure 1 – source data 2**

Comparing quantitative PCR (qPCR) and RNA-Seq data for 11 selected transcripts in *dcp2*Δ vs. WT (Figure 1C).

**Figure 1 – source data 3**

RNA-Seq analysis showing log_2_ fold-changes in mRNA abundance by *dcp2*Δ vs. *dcp2*Δ-*EE* for all transcripts (Figure 1D).

**Figure 1 – source data 4**

RNA-Seq analysis showing log_2_ fold-changes in mRNA abundance by *dcp2*Δ, *dhh1*Δ, and *dcp2*Δ*dhh1*Δ all relative to WT for all transcripts, Dhh1-dependent mRNA_up_*dcp2*Δ and Dhh1-independent mRNA_up_*dcp2*Δ transcripts (Figure 1E-F).

**Figure 1 – figure supplement 1 – source data 1**

RNA-Seq analysis showing log_2_ fold-changes in mRNA abundance by *dhh1*Δ vs. WT for all transcripts from three published data sets (Figure 1 – figure supplement 1B).

**Figure 1 – figure supplement 2 – source data 1**

RPKM normalized reads from RNA-Seq and Ribo-Seq for biological replicates WT, *dcp2*Δ and *dcp2*-*EE* (Figure 1 – figure supplement 2A-F); normalized mRNA densities for all transcripts in *dcp2*Δ vs. *dcp2-EE* cells (Figure 1 – figure supplement 2G).

**Figure 2 – source data 1**

RNA-Seq analysis showing log_2_ fold-changes in mRNA abundance by *pat1*Δ, *dhh1*Δ*pat1*Δ, *edc3scd6*Δ and *upf1*Δ all relative to WT for all transcripts (Figure 2A-C).

**Figure 2 – source data 2**

mRNA properties for all transcripts including TE in WT cells, stAI, CDS lengths, half-lives and mRNA abundance (Figure 2D-H).

**Figure 2 – figure supplement 1 – source data 1**

RNA-Seq analysis showing log_2_ fold-changes in mRNA abundance by *upf1*Δ, *upf2*Δ, and *upf3*Δ all relative to WT for all transcripts (Figure 2 – figure supplement 1).

**Figure 3 – source data 1**

List of iESR and rESR transcripts (Figure 3A-B).

**Figure 3 – source data 2**

Capped to total mRNA ratios (C/T) for all mRNAs in WT, *dcp2*Δ, *xrn1*Δ, *dcp2*Δ vs. WT and *xrn1*Δ vs. WT (Figure 3D-E).

**Figure 3 – source data 3**

Calculation of size factor from Spike-in (S. pombe) reads and S. cerevisiae reads obtained from Rpb1 (Pol II subunit) ChIP-Seq. Pearson correlation for aligned reads was calculated between biological replicates. log_2_ fold-changes in spike-in normalized Rpb1 occupancy (ChIP-Seq) and spike-in normalized RNA-Seq data is shown for all transcripts in *dcp2*Δ vs. WT.

**Figure 3 – figure supplement 1 – source data 1**

Statistics for CAGE-Seq and parallel RNA-Seq data analysis. Data reproducibility between biological replicates for CAGE-Seq and RNA-Seq in WT, *dcp2*Δ and *xrn1*Δ (Figure 3 – figure supplement 1A-F).

**Figure 3 – figure supplement 2 – source data 1**

log_2_ fold-changes in capped mRNAs (CAGE-Seq) and total mRNA (RNA-Seq) for all transcripts in *dcp2*Δ vs. WT and *xrn1*Δ vs. WT (Figure 3 – figure supplement 2A-B). List of transcripts (total and capped) significantly derepressed in abundance by *dcp2*Δ and *xrn1*Δ (Figure 3 – figure supplement 2C-D).

**Figure 3 – figure supplement 4 – source data 1**

Codon-protection index (CPI) values for all mRNAs (Figure 3 – figure supplement 4A).

**Figure 3 – figure supplement 4 – source data 2**

log_2_ fold-changes in Rpb1 occupancies (ChIP-Seq) averaged over the CDSs for all transcripts in *dcp2*Δ vs. WT (Figure 3 – figure supplement 4B & E).

**Figure 4 – source data 1**

log_2_ fold-changes in translational efficiencies (TE) measured by Ribo-Seq and parallel RNA-Seq in *dcp2*Δ vs. WT for all transcripts, TE_up_*dcp2*Δ and TE_dn_*dcp2*Δ groups (Figure 4A-B).

**Figure 4 – source data 2**

log_2_ fold-changes in protein abundances measured by TMT-MS/MS in *dcp2*Δ vs. WT for all transcripts (Figure 4C).

**Figure 4 – source data 3**

Specific activity of Nano-Luciferase reporters in WT and *dcp2*Δ for three biological replicates and change in luciferase activity was calculated in *dcp2*Δ vs. WT with S.E.M (Figure 4D).

**Figure 4 – figure supplement 1 – source data 1**

Normalized protein abundances determined by TMT-MS/MS for biological replicates in WT and *dcp2*Δ (Figure 4 – figure supplement 1A-C).

**Figure 4 – figure supplement 1 – source data 2**

log_2_ fold-changes in protein abundances measured by TMT-MS/MS and RPFs, measured by ribosome profiling in *dcp2*Δ vs. WT for all transcripts (Figure 4 – figure supplement 1D).

**Figure 5 – source data 1**

Quantification of ribosome content relative to OD_600_ in *dcp2*Δ vs. WT cells (Figure 5G).

**Figure 5 – figure supplement 1 – source data 1**

log_2_ fold-changes in TE for all transcripts conferred by SM treatment of WT cells, or the *tma64*Δ/*tma20*Δ double mutation (Figure 5 – figure supplement 1A-B).

**Figure 5 – figure supplement 1 – source data 2**

log_2_ fold-changes in TE in *dcp2*Δ vs. *dcp2-EE* cells for all mRNAs measured by ribosome profiling (Figure 5 – figure supplement 1D).

**Figure 6 – source data 1**

log_2_ fold-changes in TE by *dcp2*Δ, *dhh1*Δ and *dcp2*Δ*dhh1*Δ all relative to WT for all transcripts, Dhh1-dependent TE_up_*dcp2*Δ and Dhh1-independent TE_up_*dcp2*Δ transcripts (Figure 6A-B).

**Figure 7 – source data 1**

List of genes in each functional groups analyzed including, Glucose repressed (119), NCR (41), metabolism of energy reserves (37), and respiratory genes (60), Glycosylation (56), Sulfur metabolism (16), Unfolded protein response (94), Agglutinin (16), Autophagy related genes (26), and ribosomal protein genes (148) (Figure 7A-B & E-F; Figure 7 – figure supplement 2G-H; Figure 7 – figure supplement 4A, C-D).

**Figure 7 – source data 2**

Raw files showing expression of mitochondrial proteins with GAPDH control for three biological replicates (Figure 7C).

**Figure 7 – source data 3**

Mitochondria membrane potential using flow cytometry in WT and *dcp2*Δ for three biological replicates measured with fluorescent dye TMRM (Figure 7D).

**Figure 7 – figure supplement 1 – source data 1**

GO (gene ontology) analysis (MIPS functional classification) for the significantly derepressed and repressed transcripts for mRNA, TE, and Ribo changes (Figure 7 – figure supplement 1).

**Figure 8 – figure supplement 2 – source data 1**

List of transcripts including iESR, Msn2-iESR, non-Msn2-iESR and Msn2-targets (Figure 8 – figure supplement 1A-B).

**Figure 8 – figure supplement 3 – source data 1**

Mitochondrial protein mRNAs that bind to Puf3 and also exhibit increased expression of the encoded proteins in *puf3*Δ cells (Puf3 “cis” targets) (Figure 8 – figure supplement 3).

## Supplementary Material

**Figure 1 – figure supplement 1.**
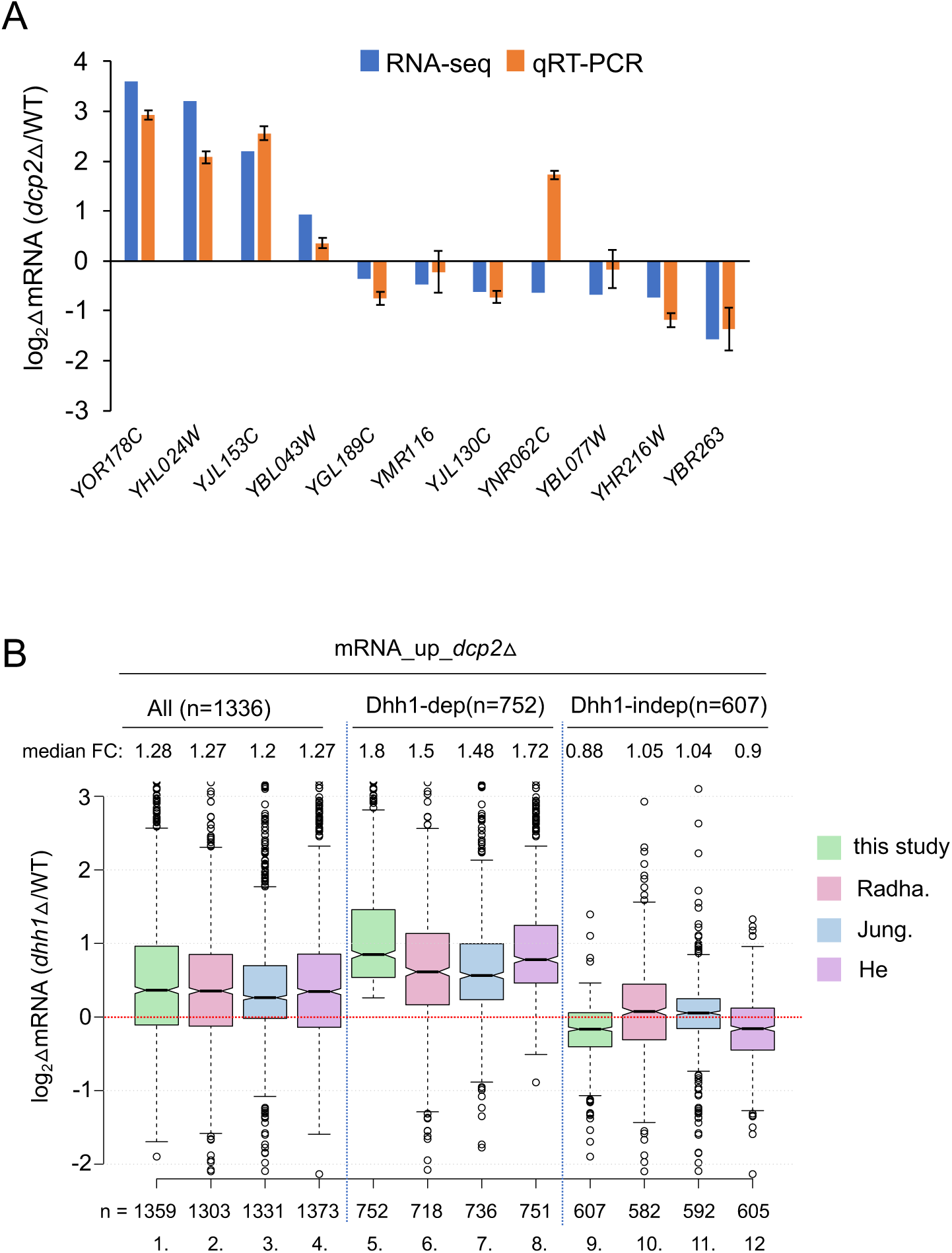
Supporting information that Dcp2 regulates mRNA abundance in a manner dependent or independent of Dhh1. **(A)** log_2_ fold-changes in mRNA abundance conferred by *dcp2*Δ vs. WT for 11 different genes determined by RNA-Seq (blue) or qRT-PCR (orange). Expression of each transcript measured by qRT-PCR was normalized to a luciferase RNA spike-in added to the total RNA. The results represent average values with standard deviations derived from at least three independent RNA preparations. **(B)** Notched box-plot as in Figure 1B showing log_2_ fold-changes in mRNA abundance in *dhh1*Δ vs. WT for all mRNA_up_*dcp2*Δ transcripts or for the Dhh1-dependent or -independent subsets of these mRNAs calculated from data obtained here (this study), in Jungfleisch et al. (2017) (Jung.), Radhakrishnan et al. (2016) (Radha.), or He et al. (2018) (He). The numbers of mRNAs for which data were obtained in each study are indicated at the bottom.

**Figure 1 – figure supplement 2.**
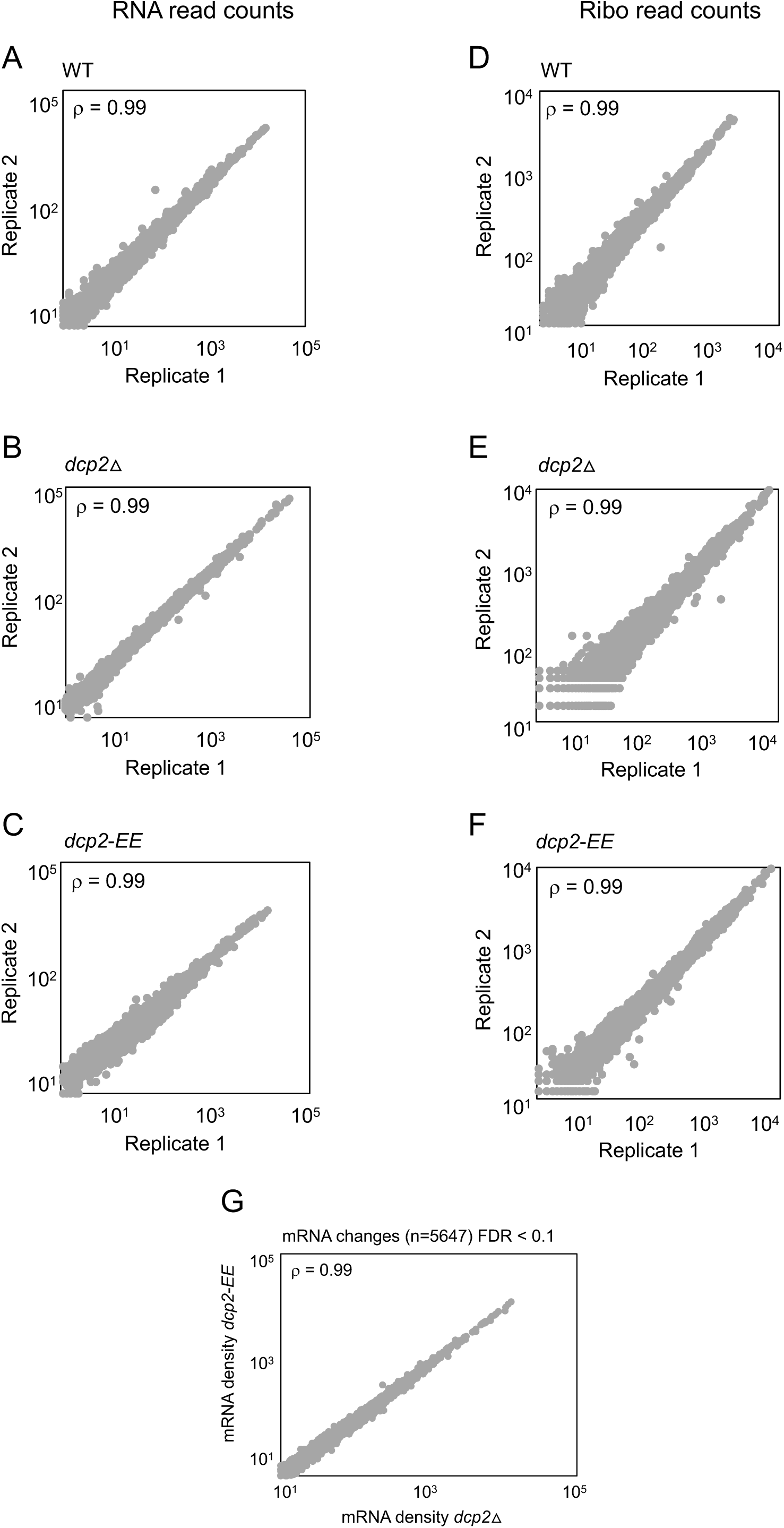
Reproducibility between biological replicates of ribosome footprint profiling and RNA-Seq analyses for WT, *dcp2*Δ and *dcp2-EE*. Scatterplots of RPKM-normalized RNA-Seq (A–C) and Ribo-Seq (D-F) read densities for all expressed mRNAs for biological replicates of the WT (A, D), *dcp2*Δ (B, E) and *dcp2*-*EE* (C, F) strains. Spearman correlation coefficients (ρ) for the plotted genes is indicated in each plot. **(G)** No significant changes in mRNA abundance between *dcp2*Δ and *dcp2-EE.* Scatterplot of normalized mRNA densities as in Figure 1A for *dcp2*Δ vs. *dcp2*-*E149Q,E153Q* (*dcp2-EE*) cells, showing a Spearman correlation coefficient of 0.99

**Figure 2 – figure supplement 1.**
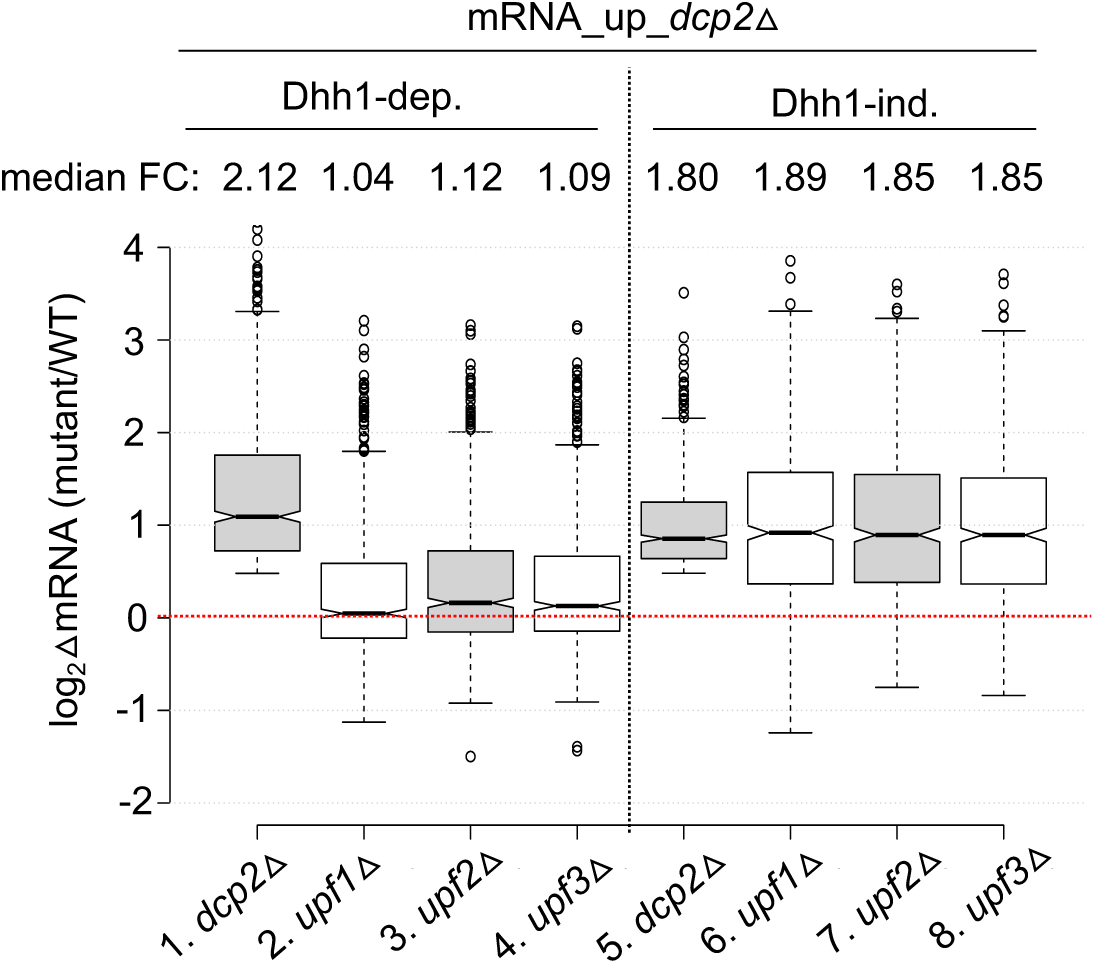
All three deletion strains lacking a Upf factor exhibit similar changes in mRNA abundances for the two groups of transcripts regulated by Dcp2. Notched box-plot as in Figure 1B showing log_2_ fold-changes in mRNA abundance in *dcp2*Δ, *upf1*Δ, *upf2*Δ, and *upf3*Δ vs. WT for the Dhh1-dependent or -independent subsets of these mRNAs. Data for *upf1*Δ, *upf2*Δ, and *upf3*Δ vs. WT were taken from Celik et al., 2017.

**Figure 3 – figure supplement 1.**
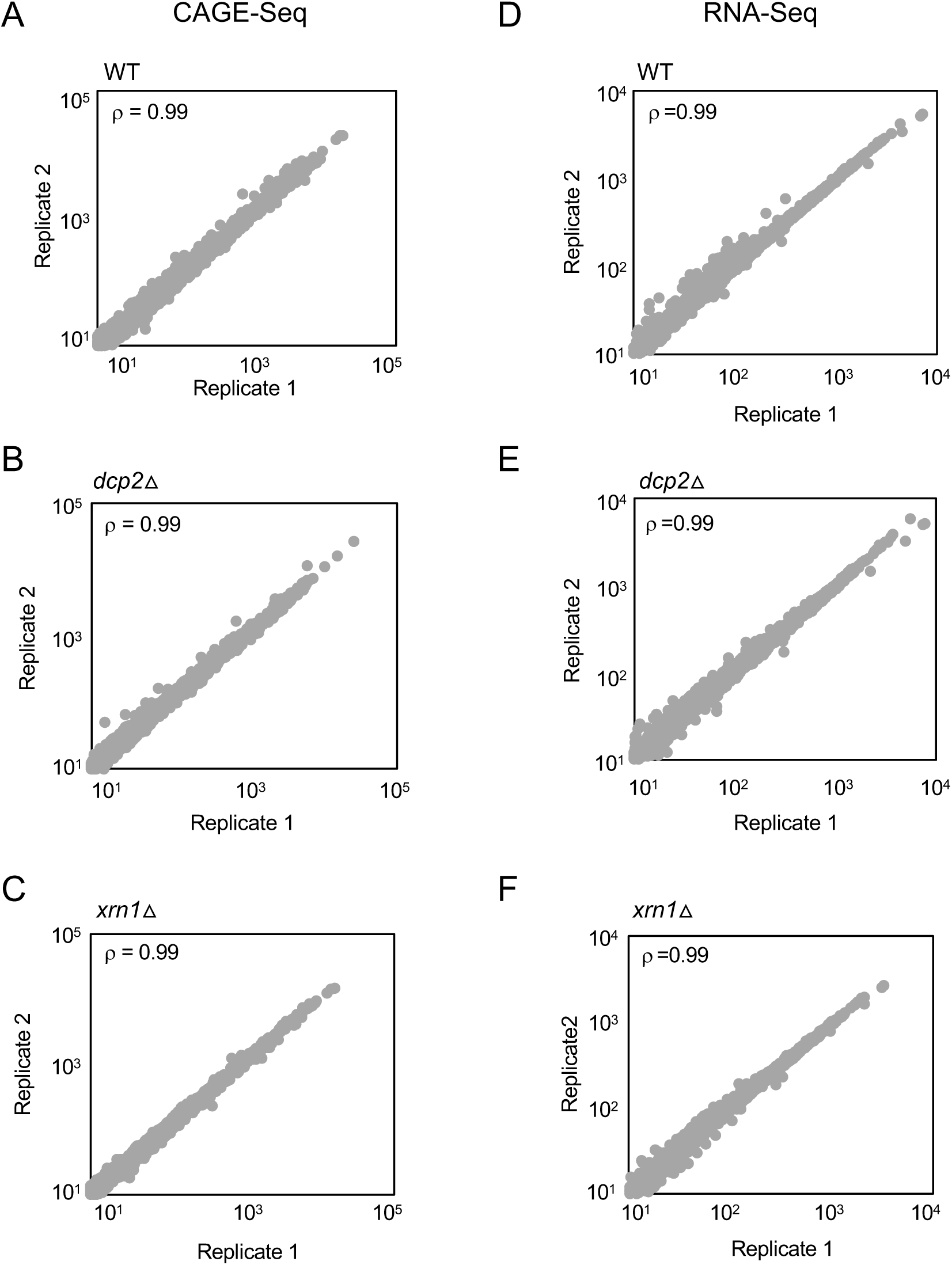
Reproducibility between biological replicates of CAGE-Seq and parallel RNA-Seq analyses for WT, *dcp2*Δ and *xrn1*Δ strains. Scatterplots of TPM-normalized CAGE reads (A-C) or TPM-normalized RNA-Seq (D-F) reads for all expressed mRNAs between biological replicates of the WT (A, D), *dcp2*Δ (B, E) and *xrn1*Δ (C, F). The Spearman correlation coefficient (ρ) is indicated in each plot.

**Figure 3 – figure supplement 2.**
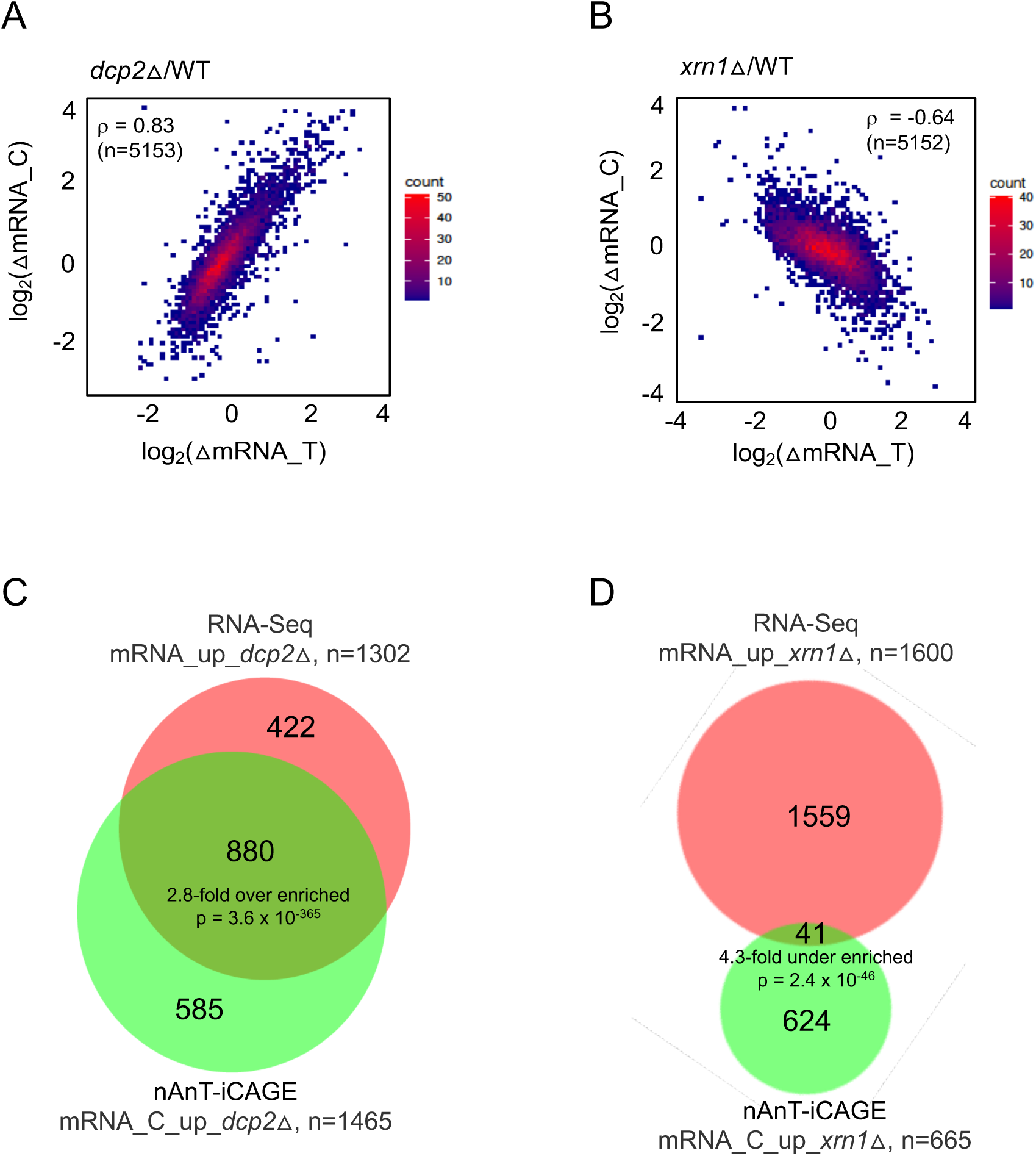
mRNAs derepressed in abundance by *dcp2*Δ or *xrn1*Δ differ in accumulating as capped (*dcp2*Δ) or uncapped (*xrn1*Δ) species. **(A-B)** Density scatterplots of log_2_ fold-changes in abundance of capped mRNA (from CAGE-Seq) vs. total mRNA (from RNA-Seq) in mutant vs. WT for (A) *dcp2*Δ or (B) *xrn1*Δ cells. Red and blue color represents higher and lower density of points, respectively **(C-D)** Proportional venn diagrams showing overlaps between total and capped mRNAs that are significantly derepressed in abundance in mutant vs. WT for *dcp2*Δ (C) or *xrn1*Δ (D) cells.

**Figure 3 – figure supplement 3.**
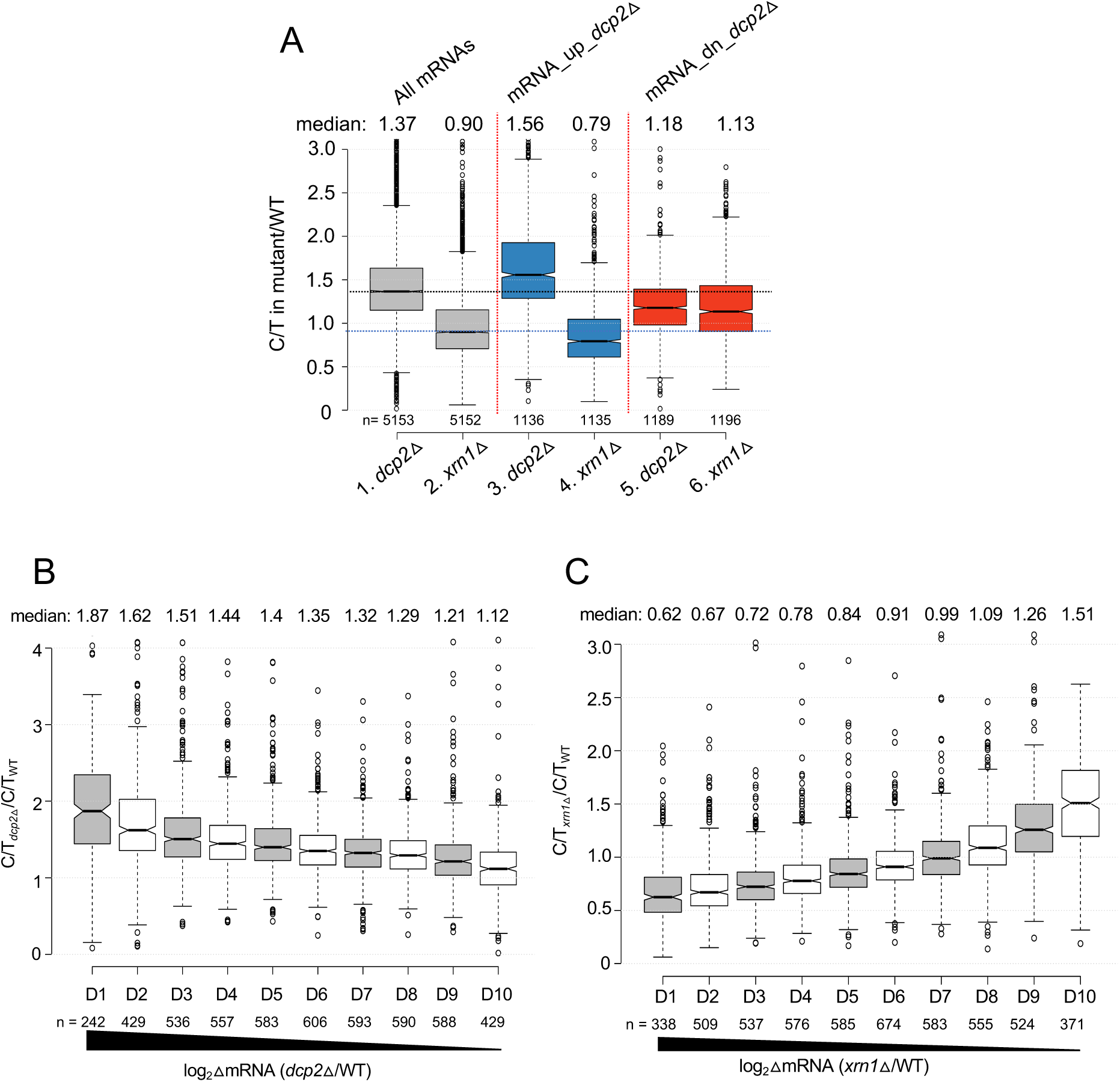
Supporting evidence that mRNAs derepressed by *dcp2*Δ exhibit greater than average levels of both decapping and degradation of uncapped intermediates by Xrn1. **(A)** Notched box-plot showing proportions of capped mRNAs (C/T ratio) in *dcp2*Δ or *xrn1*Δ vs. WT cells for all mRNAs and for the two sets of mRNA_up_*dcp2*Δ or mRNA_dn_*dcp2*Δ transcripts. **(B-C)** Notched box-plots of C/T ratios in *dcp2*Δ vs. WT (B) or *xrn1*Δ vs. WT (C) across 10 deciles of transcripts binned according to their derepression in the corresponding mutant relative to WT, progressing left to right from highest to lowest derepression ratios.

**Figure 3 – figure supplement 4.**
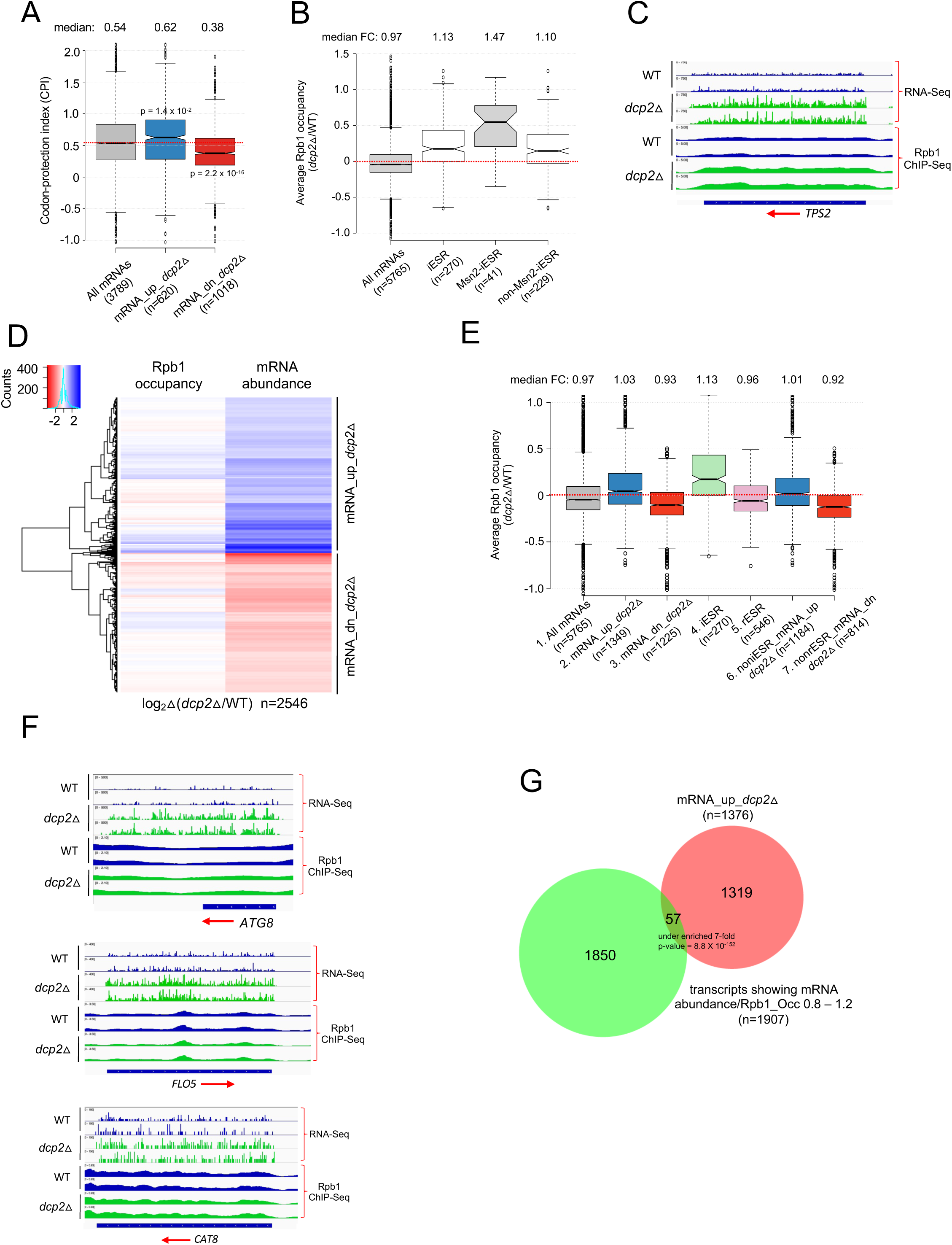
Analysis of the Codon-protection Index and Rpb1 occupancies determined by ChIP-Seq indicate that mRNA derepression by *dcp2*Δ is primarily due to loss of decapping-mediated cotranslational mRNA decay. **(A)** Dcp2-repressed mRNAs exhibit greater than average co-translational decay. Notched box-plot of the codon-protection index (CPI) for all mRNAs or the sets of mRNA_up_*dcp2*Δ or mRNA_dn_*dcp2*Δ transcripts. **(B)** Notched box-plots showing changes in relative Rpb1 occupancies averaged over the CDSs in *dcp2*Δ vs. WT cells for all mRNAs, the 270 iESR genes, the 41 iESR genes found to bind Msn2 in their promoter regions following a shift from glucose to glycerol as carbon source (Elfving, Chereji et al. 2014), and the remaining iESR genes lacking detectable Msn2 binding. **(C)** IGV depiction of relative mRNA abundance (RNA-Seq) or Pol II occupancy over the CDS (Rpb1 ChIP-Seq) for the *TPS2* gene in *dcp2*Δ vs. WT cells, showing two biological replicates for each. **(D)** Hierarchical clustering analysis of changes in relative Rpb1 occupancies averaged over the CDSs and changes in relative mRNA abundance in *dcp2*Δ vs. WT cells for the mRNA_up_*dcp2*Δ and mRNA_dn_*dcp2*Δ groups identified in Figure 1 (excluding a few outliers with log_2_Δ values > +4 or < -4). **(E)** Notched box-plots showing changes in relative Rpb1 occupancies averaged over the CDSs in *dcp2*Δ vs. WT for all mRNAs, mRNAs either derepressed or repressed by *dcp2*Δ defined in Figure 1A-B, iESR and rESR transcripts, and the non-ESR subsets of the mRNAs derepressed or repressed by *dcp2*Δ. **(F)** IGV depictions presented as in (C) for the indicated three genes. **(G)** Overlap between the 1376 derepressed mRNAs by *dcp2*Δ (RNA-Seq) and the transcripts showing similar changes in mRNA abundance and Rpb1 occupancy in *dcp2*Δ vs. WT i.e., ratio of mRNA abundance vs. Rpb1 occupancy is 0.8 to 1.2.

**Figure 4 – figure supplement 1.**
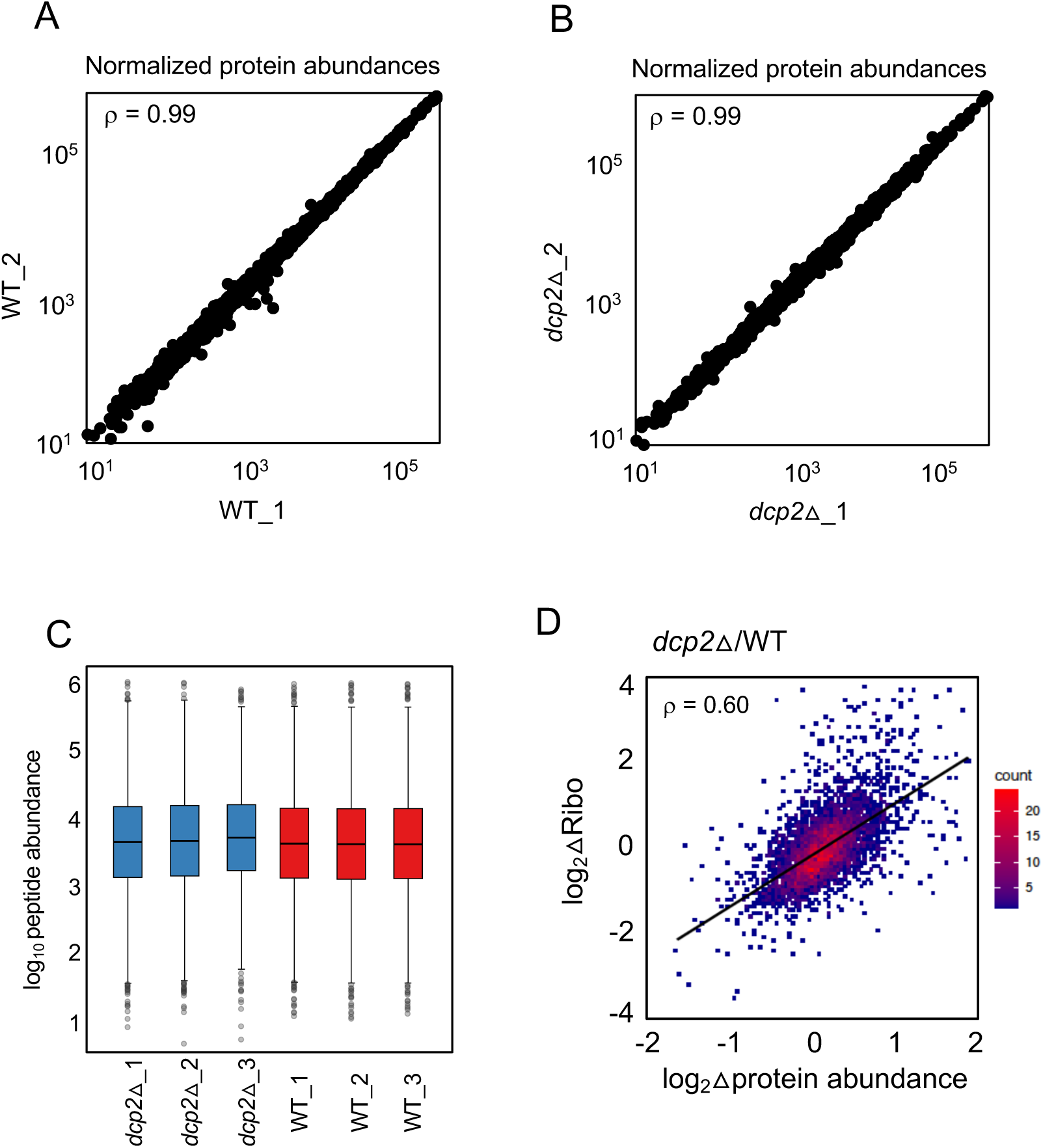
Marked correlation between RPF changes (from ribosome profiling) and protein abundance changes (from TMT-MS/MS) conferred by *dcp2*Δ. **(A-B)** Scatterplot displaying normalized protein abundances determined by TMT-MS/MS for biological replicates, with Spearman correlation coefficients indicated in each plot. **(C)** Box-plots showing the distribution of normalized peptide abundance for three biological replicates each for *dcp2*Δ and WT strains. **(D)** Density-scatterplot of log_2_ fold-changes of RPFs vs. protein abundance for all genes detected in both ribosome profiling and TMT-MS/MS experiments, respectively. Red and blue color represents higher and lower density of points, respectively and black line represents linear regression. Spearman correlation coefficient (ρ) is displayed in the plots.

**Figure 4 – figure supplement 2.**
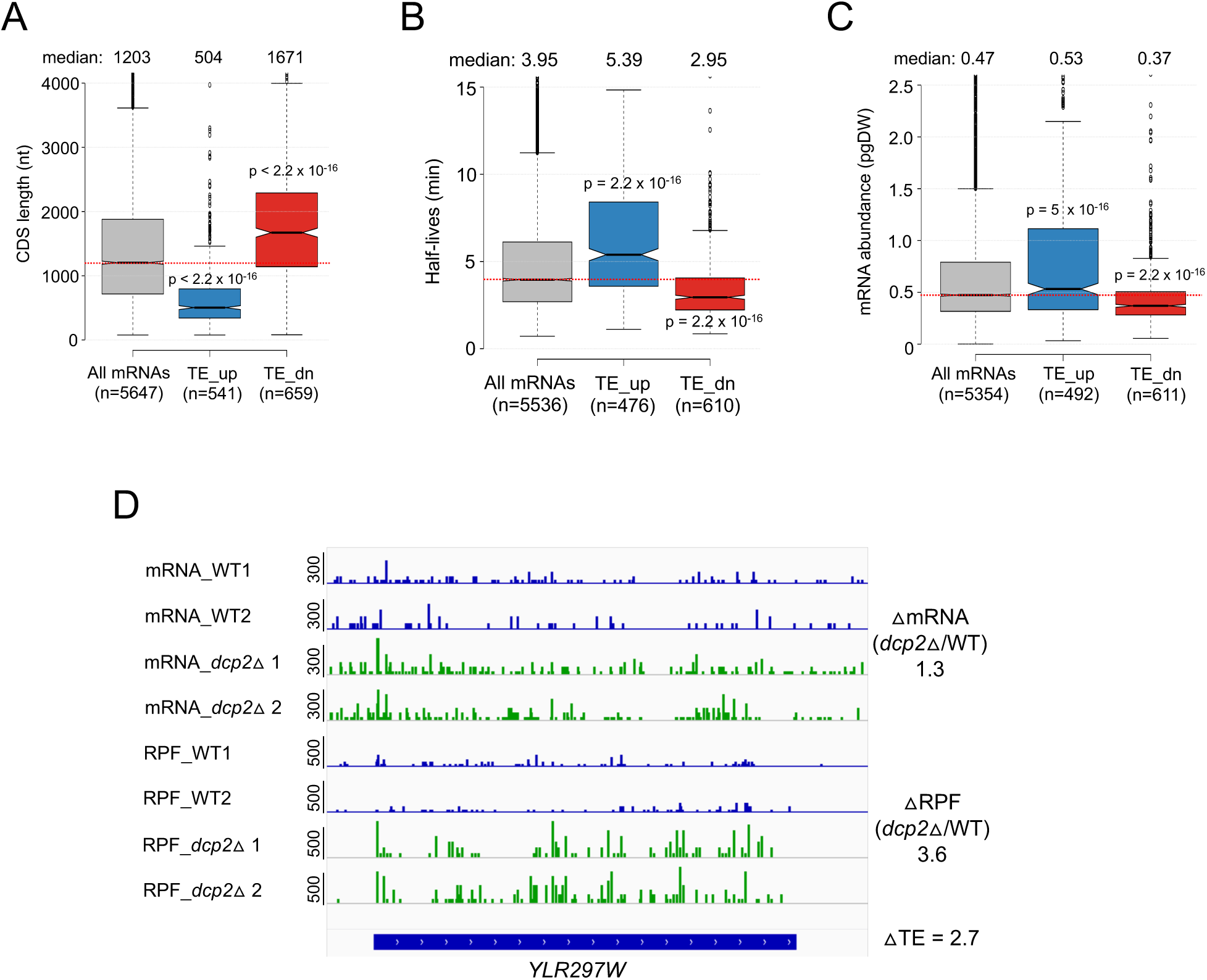
Dcp2-translationally repressed transcripts have properties associated with well-translated mRNAs. **(A-C)** Notched box-plots of CDS lengths (A), mRNA half-lives (B), mRNA abundances (C) for all mRNAs and the two sets of mRNAs translationally repressed or stimulated by Dcp2. **(D)** IGV depiction of a representative gene exhibiting increased TE in *dcp2*Δ vs. WT cells, presented as in Figure 4G.

**Figure 5 – figure supplement 1.**
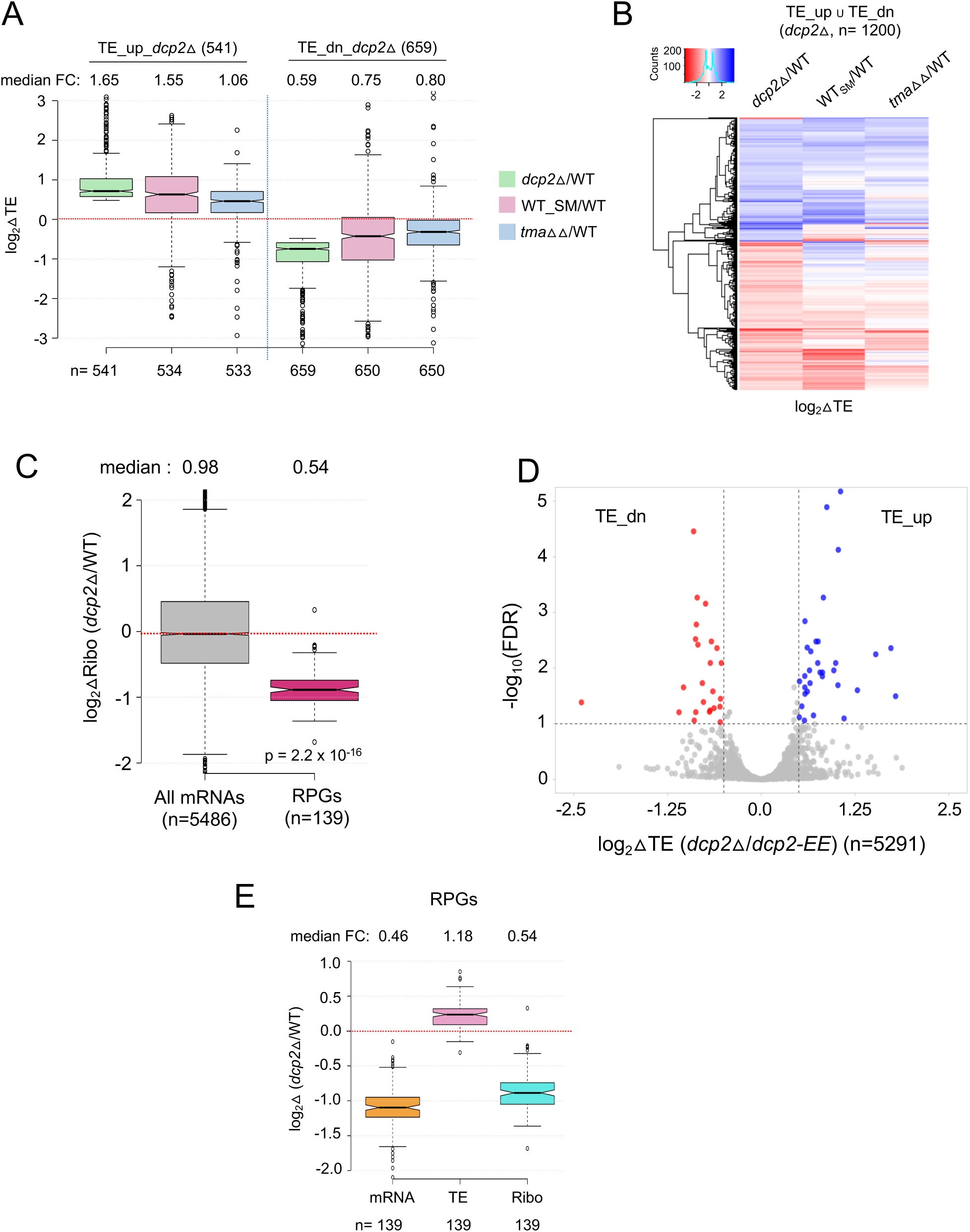
Supporting evidence that *dcp2*Δ evokes translational reprogramming by increasing competition for limiting PICs. **(A)** Notched box-plot of log_2_ fold-changes in TE conferred by *dcp2*Δ, SM treatment of WT cells, or the *tma64*Δ/*tma20*Δ double mutation, all determined by ribosome profiling, for the two groups of mRNAs that are translationally repressed (cols. 1-3) or translationally stimulated (cols. 4-6) by Dcp2. **(B)** Hierarchical clustering analysis of log_2_ fold-changes in TE conferred by *dcp2*Δ (col. 1), SM treatment of WT cells (col. 2), or the *tma64*Δ/*tma20*Δ double mutation (col. 3) for the 1200 mRNAs belonging to the same two groups analyzed in (B) for which data was available in all three analyses (excluding a few outliers with log_2_ΔTE values > +4 or < -4). In panels A-B, ribosome profiling data from (Gaikwad, Ghobakhlou et al. 2021) were interrogated. **(C)** Notched box-plot of log_2_ fold-changes in RPFs (measured by ribosome profiling) in *dcp2*Δ vs. WT cells for all mRNAs or RPG mRNAs. **(D)** Evidence that the majority of the TE changes conferred by *dcp2*Δ result from loss of Dcp2 catalytic activity. A Volcano-plot showing log_2_ fold-changes in TE in *dcp2*Δ vs. *dcp2-EE* cells (x-axis) vs. the -log_10_FDR values for the TE changes (y-axis) for all mRNAs. The dotted lines demarcate TE fold-changes of >1.4 at FDR < 0.1, such that 41 mRNAs (blue dots) and 30 mRNAs (red dots) display significant TE increases or decreases, respectively. **(E)** Notched box-plot showing log_2_ fold-changes in mRNA, TE or RPFs (Ribo) in *dcp2*Δ vs. WT cells for all RPGs. In accordance with the PIC competition model, the efficiently translated RPG mRNAs show increased TEs in *dcp2*Δ vs. WT cells despite strong reductions in abundance, (attributable to the rESR), for a net decrease in ribosome occupancies (RPFs/Ribo levels).

**Figure 6 – figure supplement 1.**
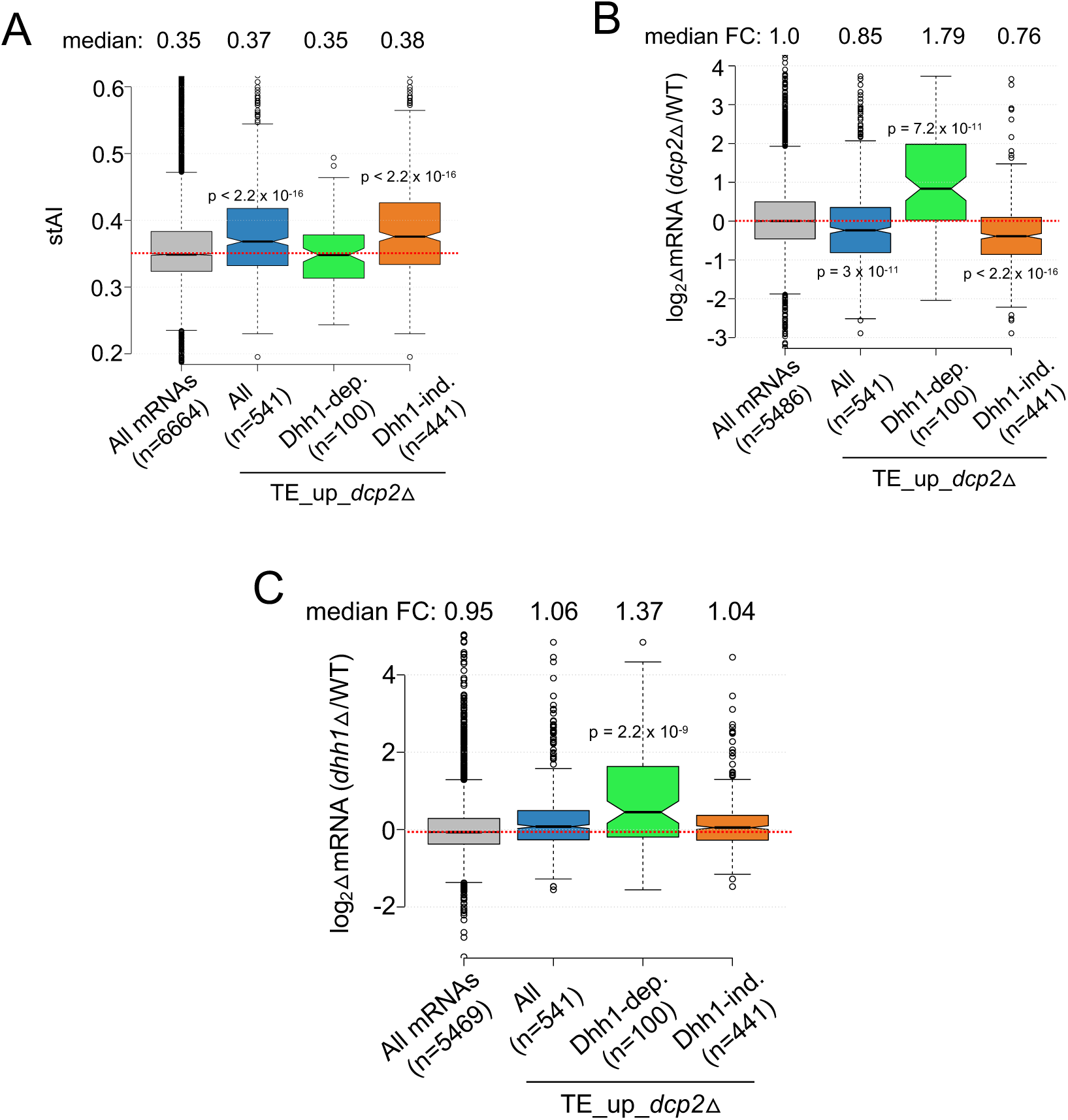
Additional evidence that mRNAs exhibiting Dhh1-dependent translational repression by Dcp2 are preferentially targeted for decapping by Dcp2 and Dhh1. **(A-C)** Notched box-plot of stAI values (A), log_2_ fold-changes in mRNA abundance in *dcp2*Δ vs. WT (B), and log_2_ fold-changes in mRNA abundance in *dhh1*Δ vs. WT (C) for all mRNAs, all TE_up_*dcp2*Δ mRNAs, or the Dhh1-dependent or Dhh1-independent subsets of the TE_up_*dcp2*Δ mRNAs.

**Figure 7 – figure supplement 1.**
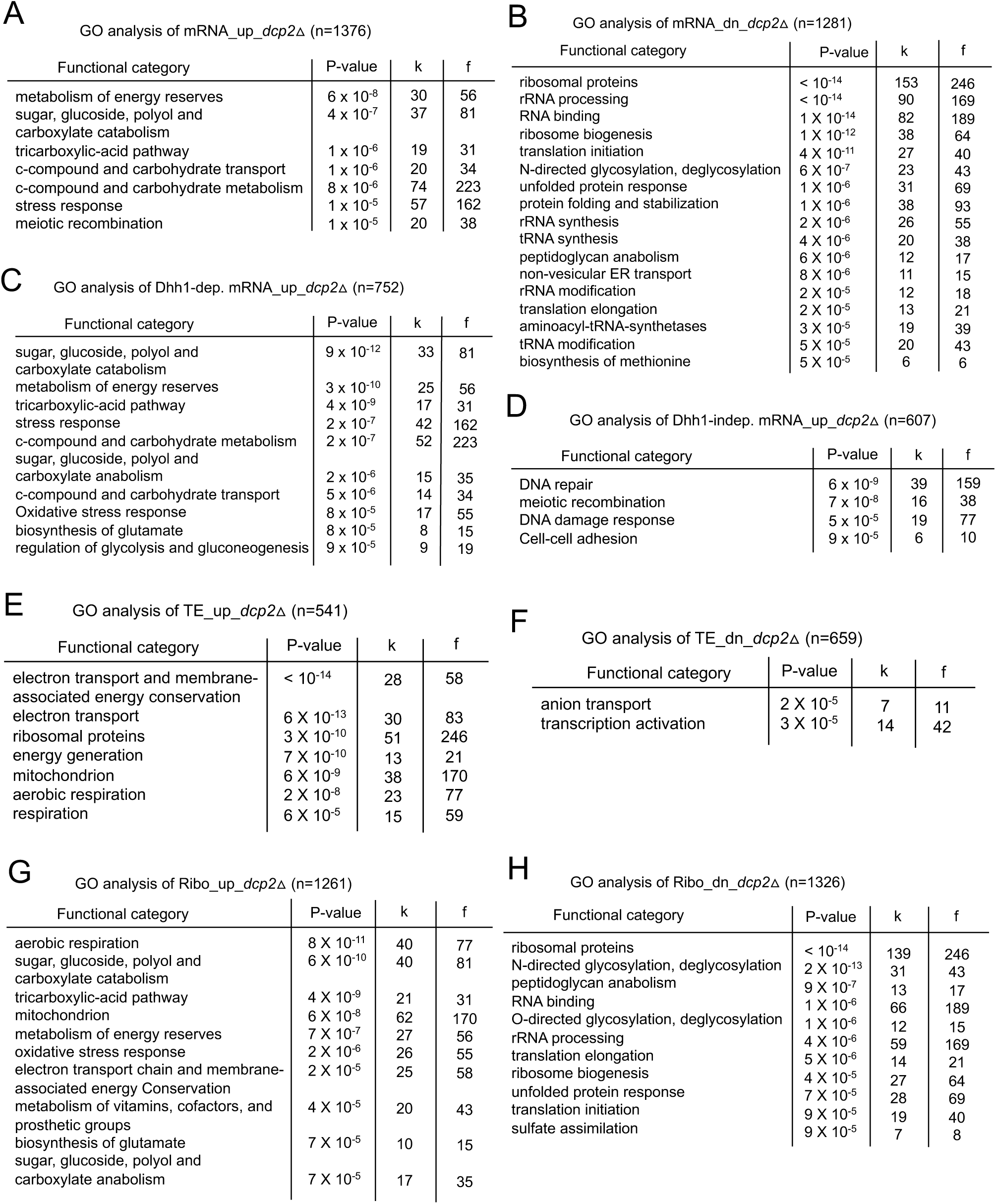
Gene ontology analysis of mRNAs dysregulated by *dcp2*Δ on YPD medium suggests a reprogramming of metabolism and protein synthesis to mimic growth on nutrient-poor medium. **(A-H)** Gene Ontology (GO) analysis was conducted on the indicated sets of mRNAs showing derepression or repression of mRNA abundance (A-D), TE (E-F), or ribosome (Ribo) abundance (G-H) in *dcp2*Δ vs. WT cells. Each panel lists the MIPS functional categories enriched among the genes encoding each group of transcripts, the P-value indicating the statistical significance of enrichment, the number of genes represented by the mRNAs in the set belonging to the functional category (k), and the total number of genes present in the functional group (f). The Bonferroni correction and a minimum P-value of 0.05 were applied.

**Figure 7 – figure supplement 2.**
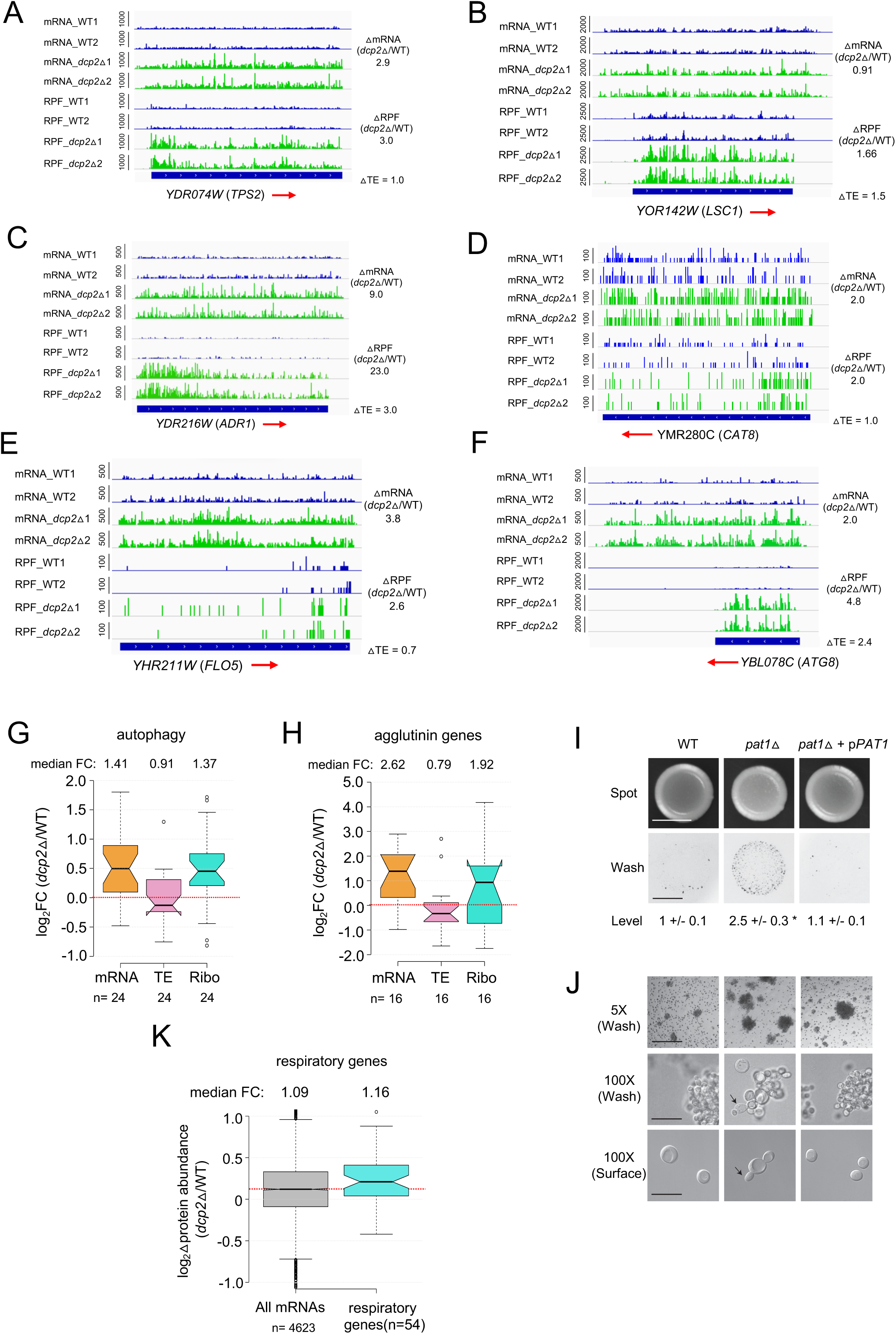
Supporting evidence that Dcp2 represses mRNA abundance, TE, or both of genes involved in different pathways not required for growth on rich medium. **(A-F)** IGV depictions of *TPS2* (A), *LSC1* (B), *ADR1* (C), *CAT8* (D), *FLO5* (E) and *ATG8* (F) showing increased mRNA and RPF abundance in *dcp2*Δ vs. WT cells, as described in Figure 4G. **(G & H)** Notched box-plots showing log_2_ fold-changes in mRNA, TE or RPFs (Ribo) in *dcp2*Δ vs. WT cells for 24 Autophagy-related genes (ATG) involved in autophagy (G) or 16 agglutinins that function in cell adhesion (H). **(I-J)** Increased invasive growth observed for *pat1*Δ vs. WT cells, as described in Figure 7G-H. (**K)** Notched box-plot of log_2_ fold-changes in protein abundance determined by TMT-MS/MS in *dcp2*Δ vs. WT cells for all genes or 54 genes encoding mitochondrial proteins with direct roles in oxidative phosphorylation.

**Figure 7 – figure supplement 3.**
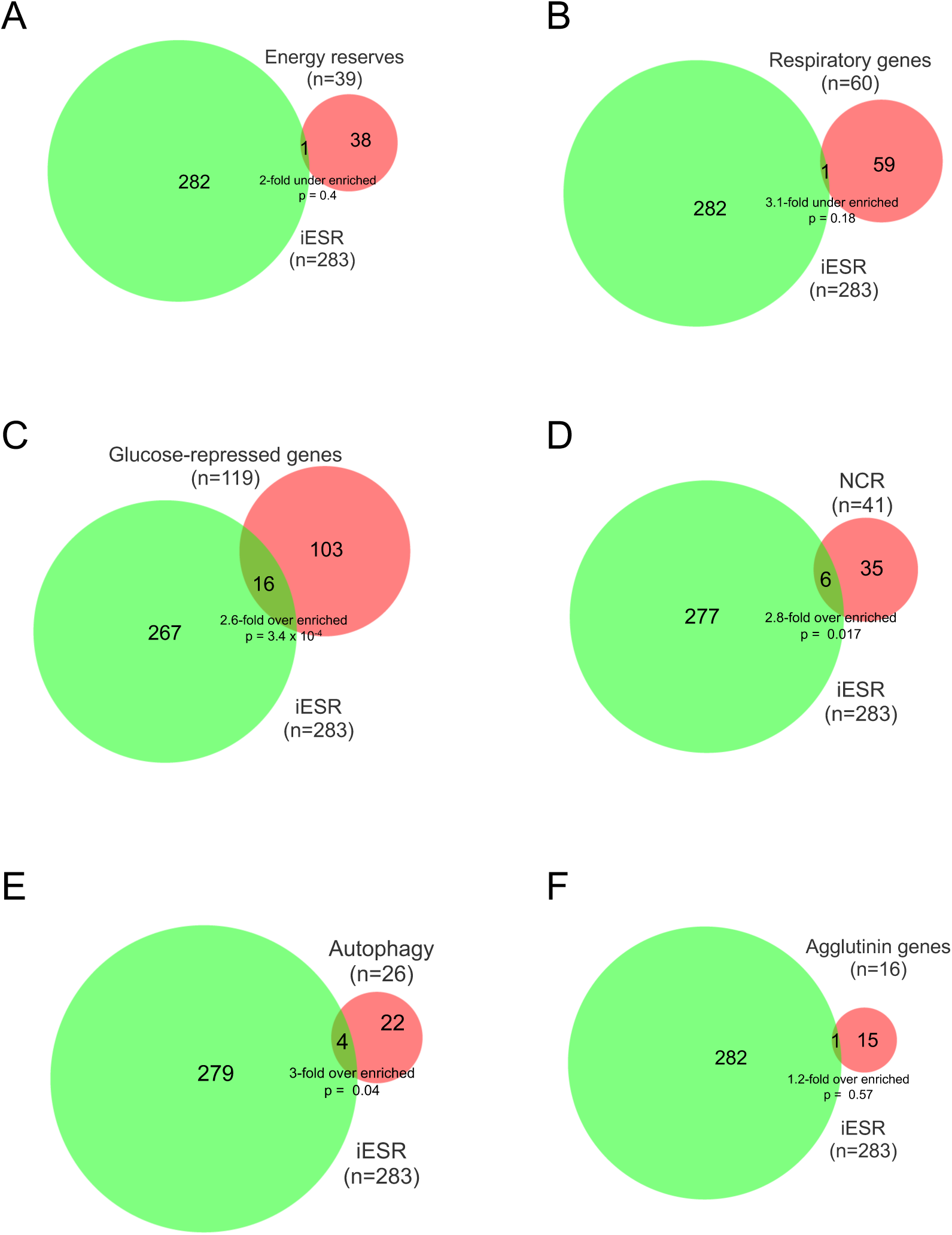
Overlap between iESR genes and genes in various pathways exhibiting up-regulation of mRNA abundance or TE in *dcp2*Δ vs. WT cells on YPD medium. **(A-F)** Overlap between the 283 iESR genes and the indicated groups comprised of all genes involved in metabolism of energy reserves (A), encoding mitochondrial proteins with direct roles in oxidative phosphorylation (B), glucose-repressed or induced by Adr1 or Cat8 (C), nitrogen-catabolite repressed (D), encoding proteins with direct roles in autophagy (E), or encoding agglutinins that function in cell adhesion (F). The number of genes in each group is indicated (n) as is the p-value of the overlap from the hypergeometric distribution.

**Figure 7 – figure supplement 4.**
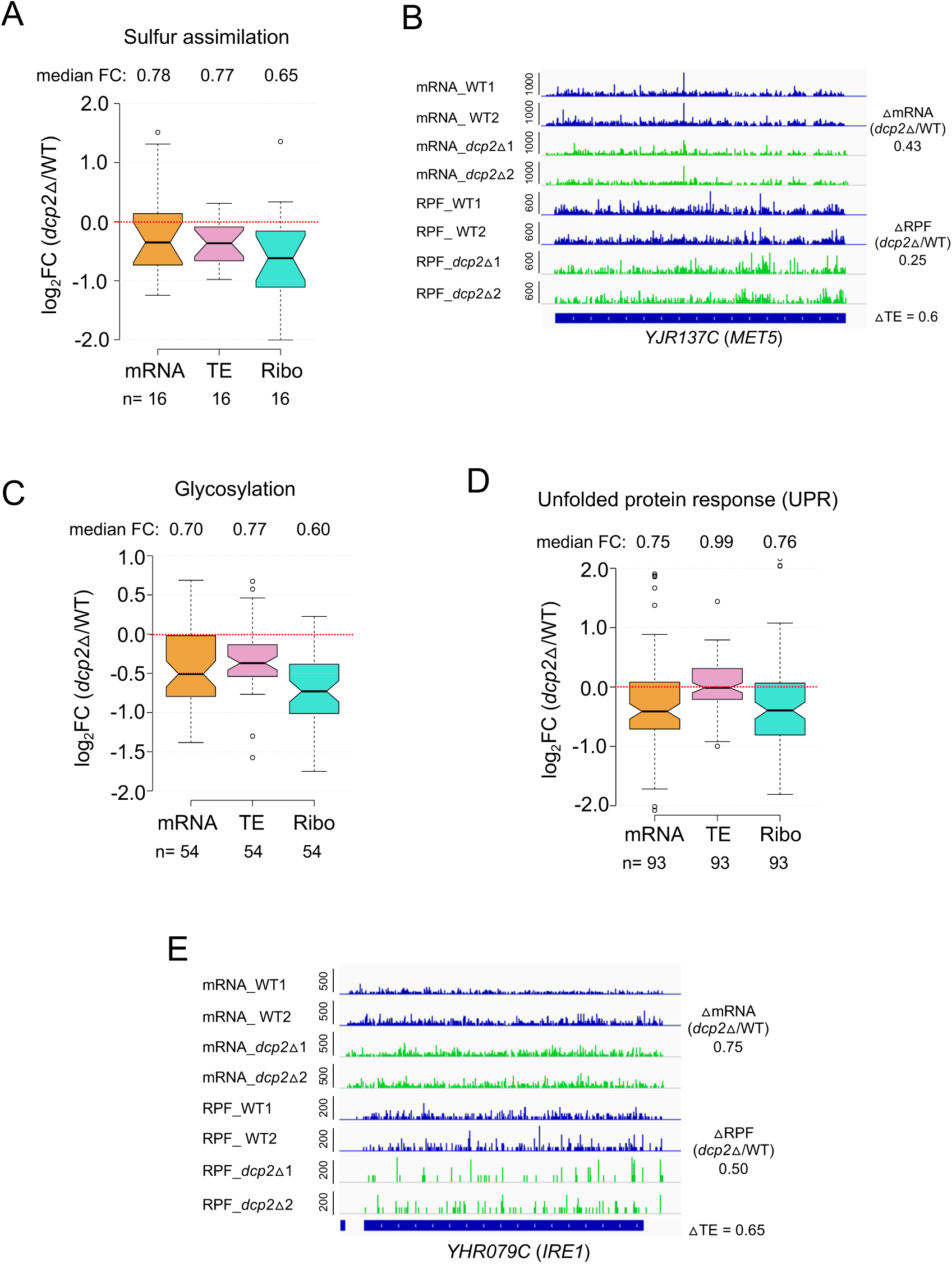
Evidence that *dcp2*Δ reduces mRNA abundance or TE of genes involved in protein glycosylation, sulphur assimilation, or the unfolded protein response on rich medium. **(A & C-D)** Notched box-plots showing log_2_ fold-changes in mRNA, TE or RPFs (Ribo) in *dcp2*Δ vs. WT cells for genes involved in protein sulphur assimilation (A), glycosylation (C), or unfolded protein response (D). The total genes for each functional groups was obtained from KEGG pathway maps for *Saccharomyces cerevisiae* **(B & E)** IGV tracks for exemplar genes *MET5* and *IRE1* showing decreased mRNA or RPF abundance in *dcp2*Δ vs. WT cells.

**Figure 8 – figure supplement 1.**
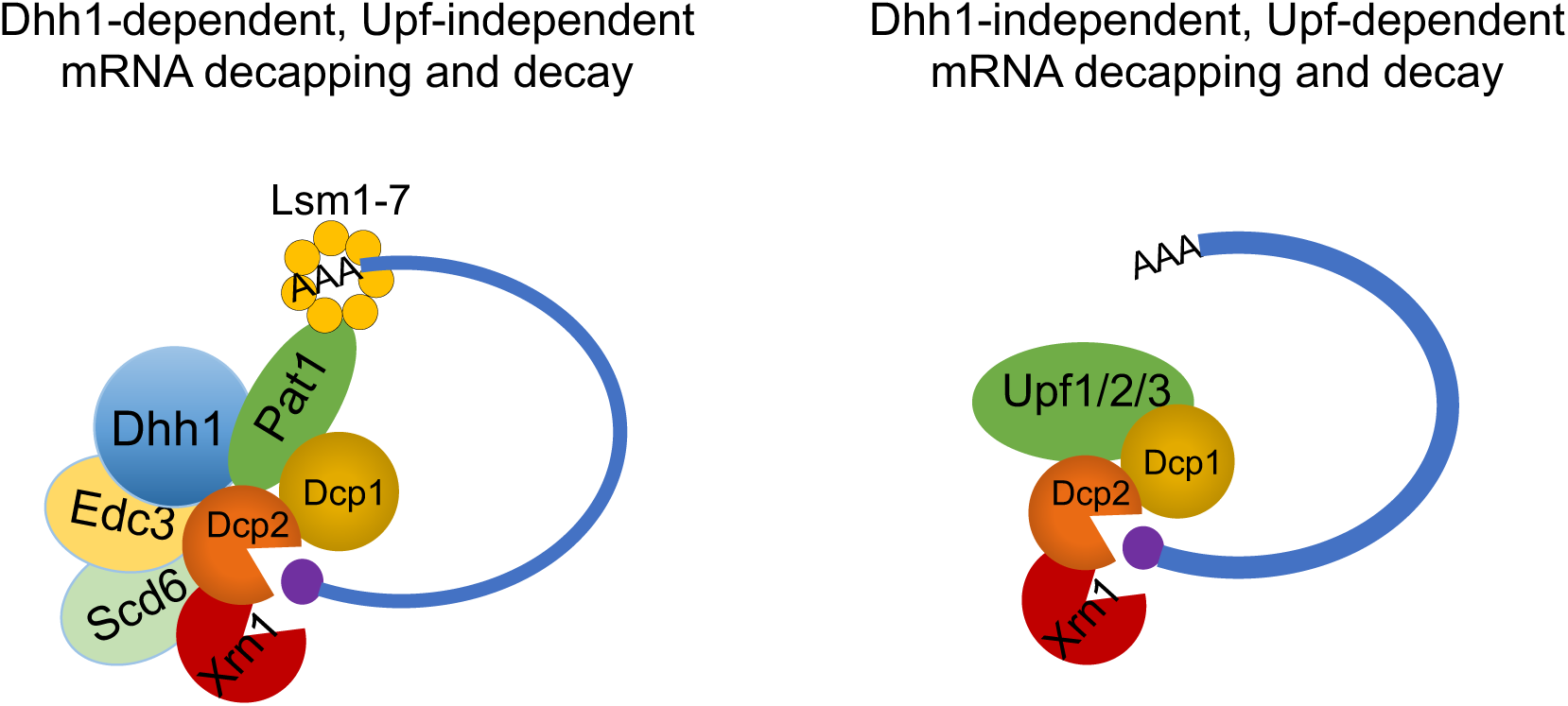
Model depicting concerted functions of multiple decapping activators in activating decapping by Dcp2 for the Dhh1-dependent and Dhh1-independent subsets of mRNAs targeted by Dcp2. **(*Left*)** Dhh1, Pat1, Scd6 and Edc3 form a complex with Dcp1/Dcp2 and Xrn1 to mediate decapping and 5’ to 3’ decay of Dhh1-dependent transcripts. **(*Right*)** Upf proteins form a complex with Dcp1:Dcp2 and Xrn1 to mediate decapping and 5’ to 3’ decay of the NMD substrates that comprise most of the Dhh1-independent transcripts.

**Figure 8 – figure supplement 2.**
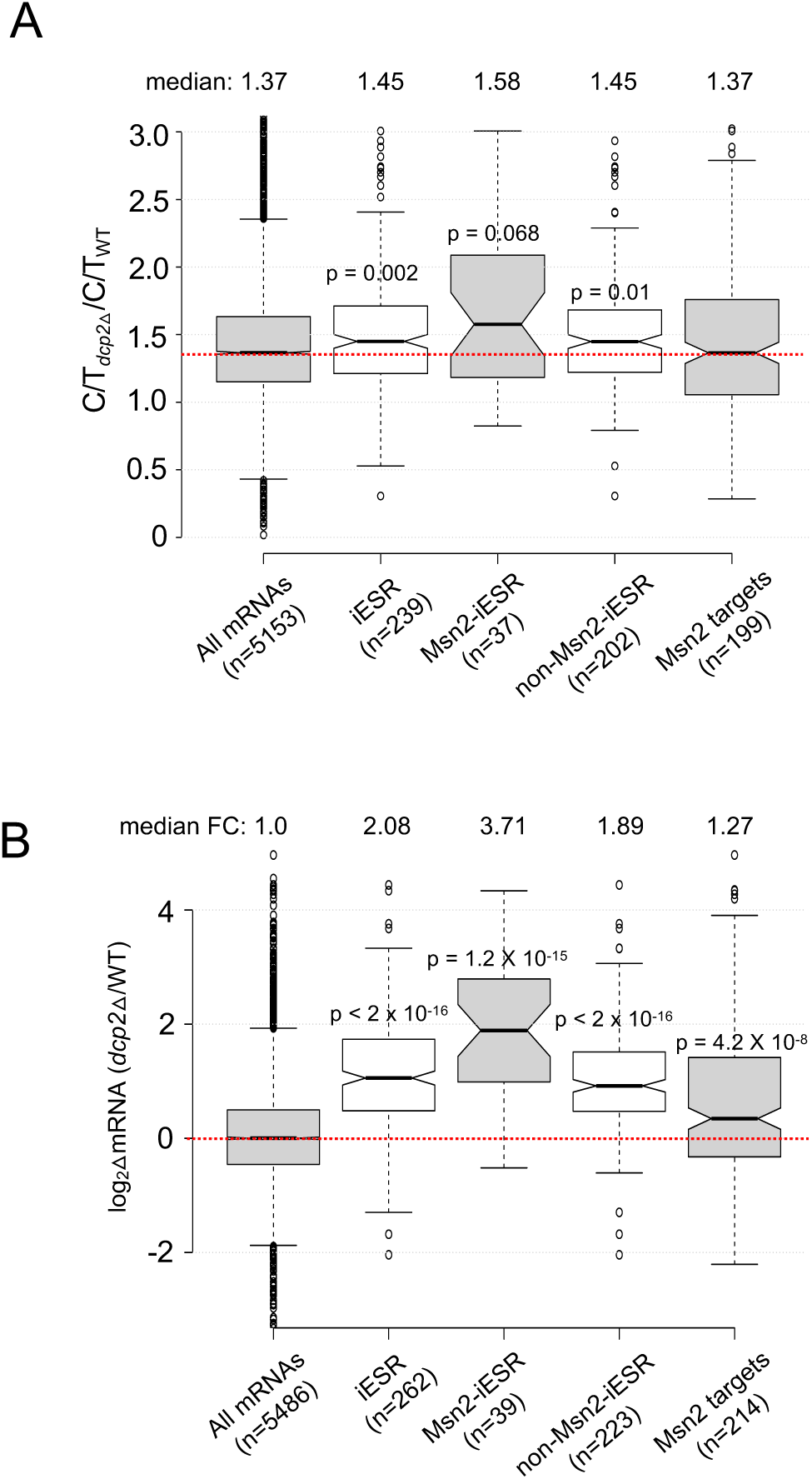
Evidence that decapping by Dcp2 contributes to the iESR. **(A & B)** Notched box-plots showing log_2_ fold-changes in relative mRNA abundance (A) and the change in C/T ratios (B) in *dcp2*Δ vs. WT cells for all mRNAs, iESR transcripts, iESR mRNAs found to bind Msn2 in their promoter regions by ChIP-Seq analysis following a shift from glucose to glycerol as carbon source (Elfving, Chereji et al. 2014), the remaining iESR genes lacking detectable Msn2 binding, and all genes shown by ChIP-Seq to bind Msn2.

**Figure 8 – figure supplement 3.**
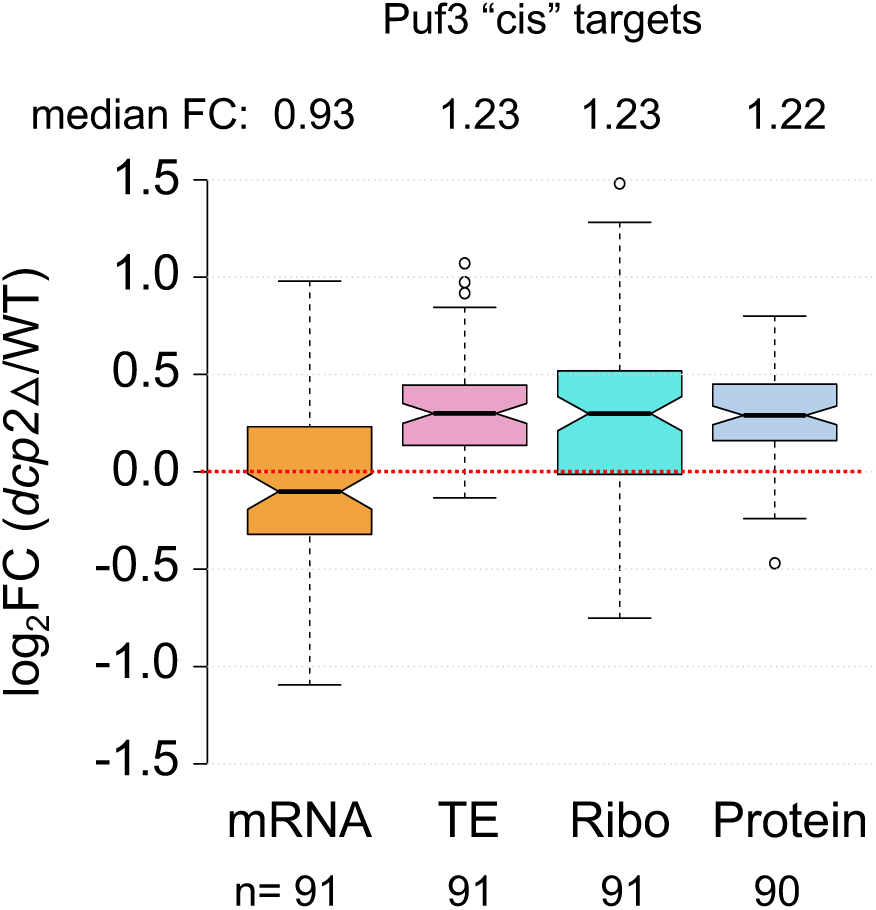
Dcp2 primarily represses the translation but not abundance of a group of high-confidence Puf3-repressed mRNAs highly enriched for mitochondrial proteins. Notched box-plots showing log_2_ fold-changes in mRNA, TE, RPFs (Ribo), or steady-state protein levels (determined by TMT-MS) in *dcp2*Δ vs. WT cells for a group of 91 mRNAs that bind Puf3 and show derepressed protein expression in *puf3*Δ vs. WT cells, of which 86 function in mitochondria, dubbed Puf3 “cis” targets by Lapointe et al. (2018).

### Supplementary Tables

**Table 1:**
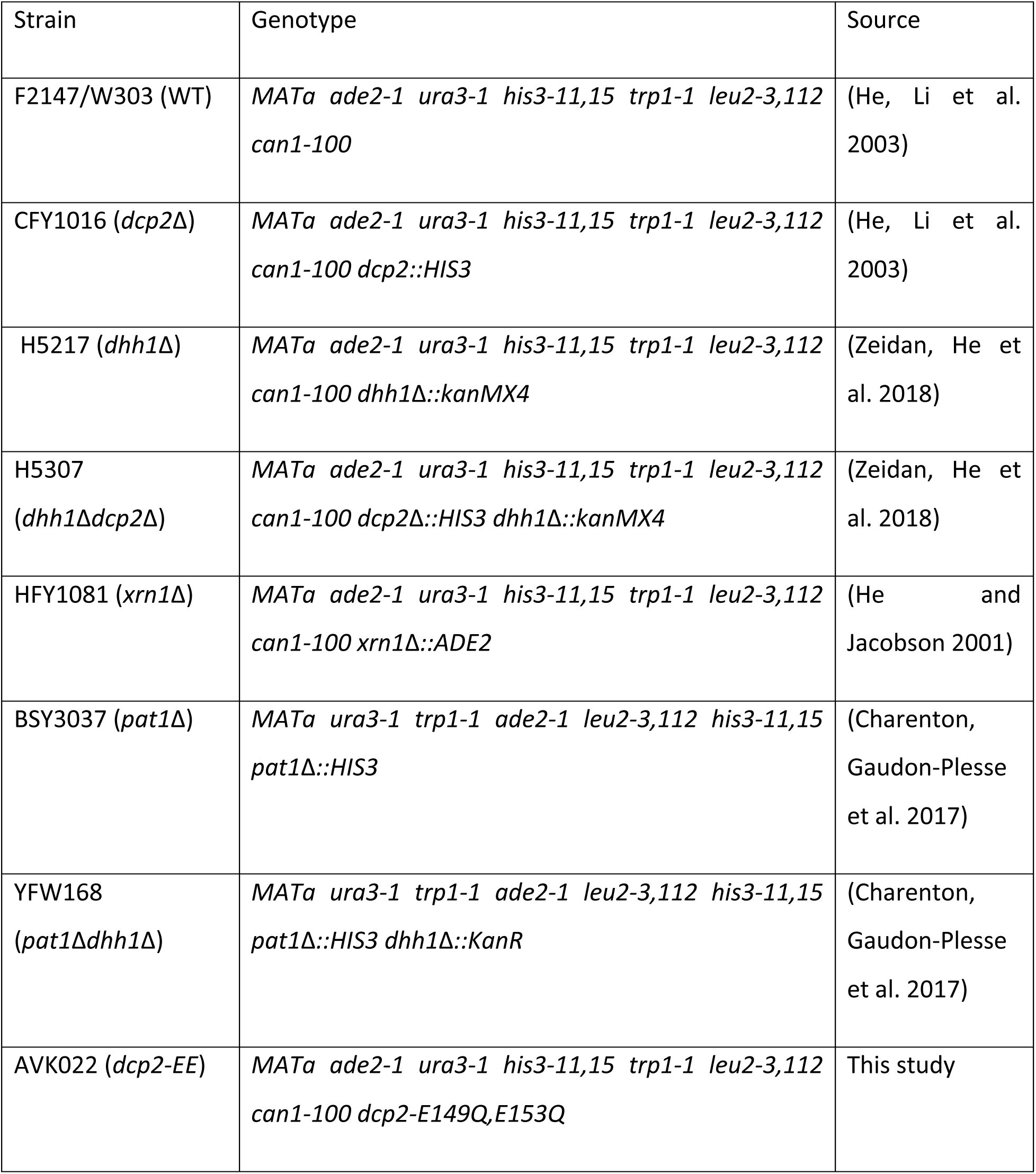
Yeast strains used in this study.

| Strain | Genotype | Source |
| --- | --- | --- |
| F2147/W303 (WT) | *MATa ade2-1 ura3-1 his3-11,15 trp1-1 leu2-3,112 can1-100* | (He, Li et al. 2003) |
| CFY1016 (*dcp2Δ*) | *MATa ade2-1 ura3-1 his3-11,15 trp1-1 leu2-3,112 can1-100 dcp2::HIS3* | (He, Li et al. 2003) |
| H5217 (*dhh1Δ*) | *MATa ade2-1 ura3-1 his3-11,15 trp1-1 leu2-3,112 can1-100 dhh1Δ::kanMX4* | (Zeidan, He et al. 2018) |
| H5307 (*dhh1Δdcp2Δ*) | *MATa ade2-1 ura3-1 his3-11,15 trp1-1 leu2-3,112 can1-100 dcp2Δ::HIS3 dhh1Δ::kanMX4* | (Zeidan, He et al. 2018) |
| HFY1081 (*xrn1Δ*) | *MATa ade2-1 ura3-1 his3-11,15 trp1-1 leu2-3,112 can1-100 xrn1Δ::ADE2* | (He and Jacobson 2001) |
| BSY3037 (*pat1Δ*) | *MATa ura3-1 trp1-1 ade2-1 leu2-3,112 his3-11,15 pat1Δ::HIS3* | (Charenton, Gaudon-Plesse et al. 2017) |
| YFW168 (*pat1Δdhh1Δ*) | *MATa ura3-1 trp1-1 ade2-1 leu2-3,112 his3-11,15 pat1Δ::HIS3 dhh1Δ::KanR* | (Charenton, Gaudon-Plesse et al. 2017) |
| AVK022 (*dcp2-EE*) | *MATa ade2-1 ura3-1 his3-11,15 trp1-1 leu2-3,112 can1-100 dcp2-E149Q,E153Q* | This study |

**Table 2:** Plasmids used in this study.

| Plasmid | Relevant description | Source or reference |
| --- | --- | --- |
| pQZ145 | *DCP2* in pRS316 | (Zeidan, He et al. 2018) |
| pAV008 | *dcp2-E149Q,E153Q* in pRS316 | This study |
| Ylplac211 | Integrating *URA3* vector | (Gietz and Sugino 1988) |
| pAV013 | *dcp2-E149Q,E153Q* in Ylplac211 | This study |
| pRS316 | *CEN6, URA3* | (Sikorski and Hieter 1989) |
| pEXfz17-2 | *FLO5-nLUC* in pRS316 | This study |
| pEXfz26-3 | *FIG2-nLUC* in pRS316 | This study |
| pEXfz21-1 | *AAD3-nLUC* in pRS316 | This study |
| pEXfz18-5 | *FLO1-nLUC* in pRS316 | This study |
| pEXfz8-4 | *AMD2-nLUC* in pRS316 | This study |
| pEXfz13-1 | *SAG1-nLUC* in pRS316 | This study |
| pEXfz4-2 | *IDP2-nLUC* in pRS316 | This study |
| pEXfz9-1 | *PCK1-nLUC* in pRS316 | This study |
| pEXfz14-1 | *SSA3-nLUC* in pRS316 | This study |
| pEXfz23-3 | *ADR1-nLUC* in pRS316 | This study |
| pEXfz6-2 | *THI21-nLUC* in pRS316 | This study |
| pEXfz1-2 | *TRK2-nLUC* in pRS316 | This study |
| pEXfz19-1 | *SNZ1-nLUC* in pRS316 | This study |
| pNG158 | *PAT1* in pRS315 | This study |

**Table 3:**
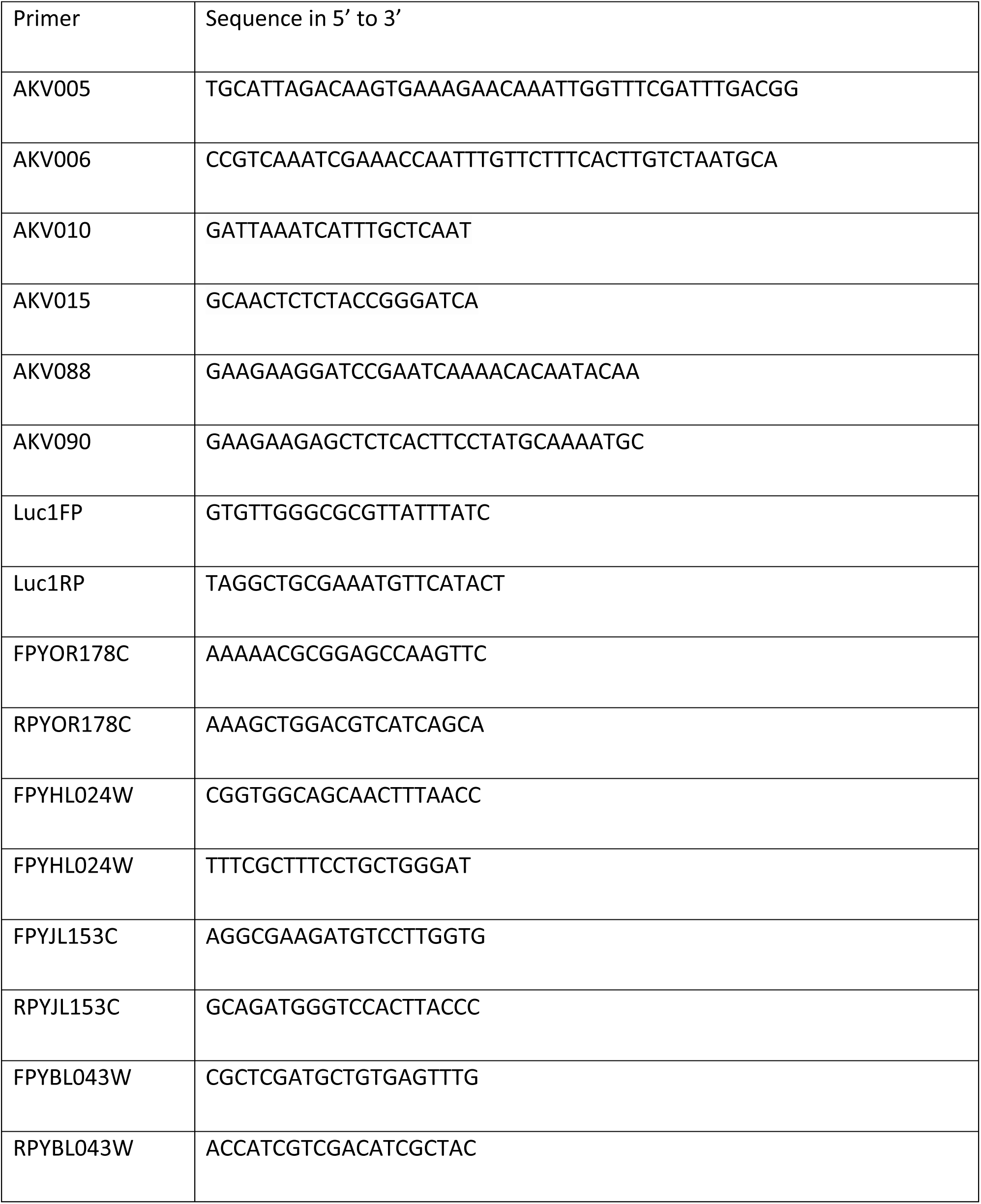
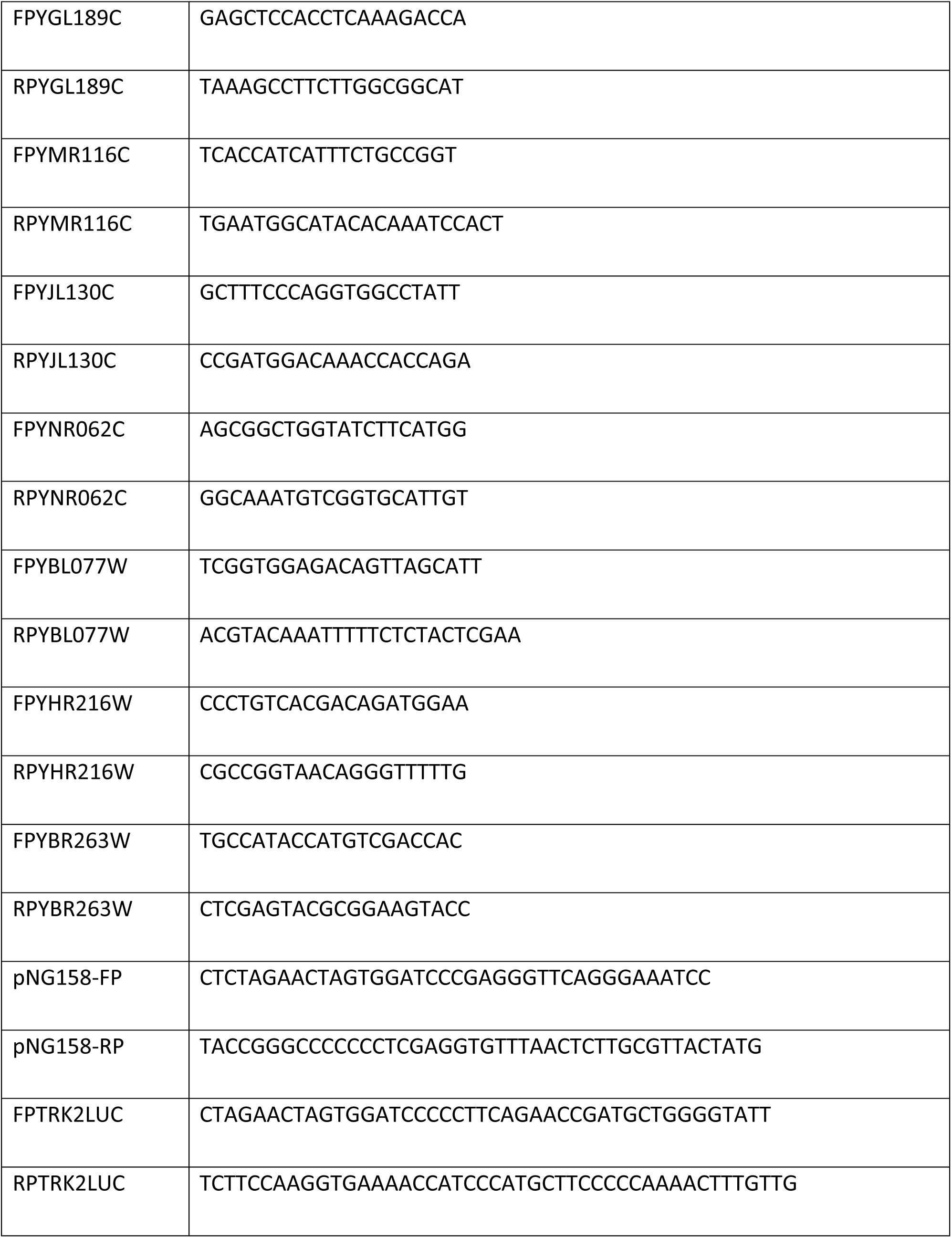
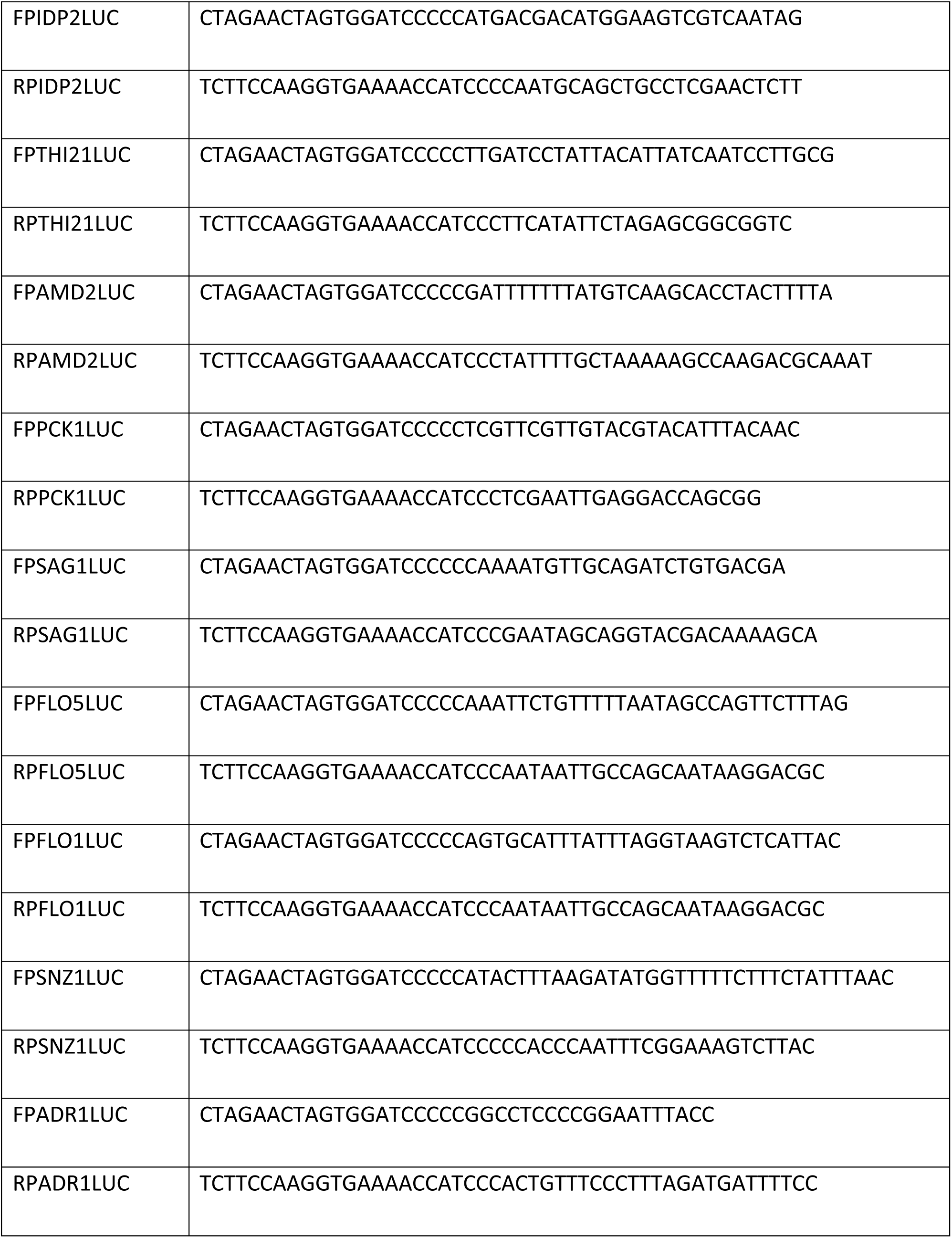
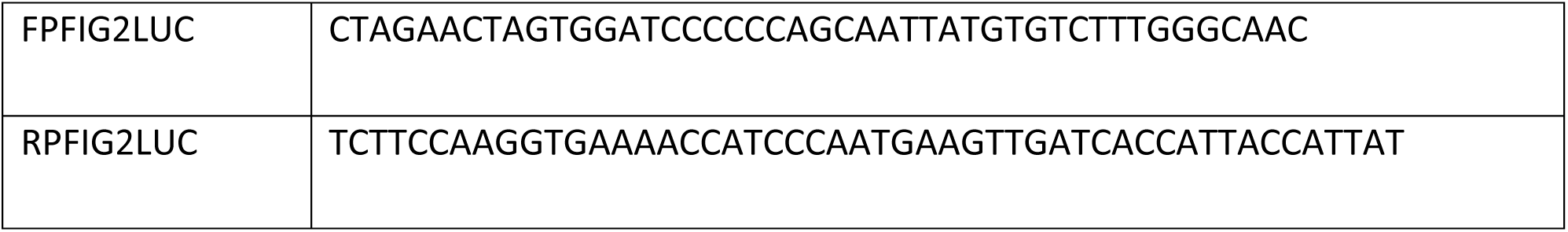
Primers used in this study.

**Table 4:** Details of nanoLUC strains.

| gene_ID | Gene name | 5' UTR length | CDS length | 3' UTR length |
| --- | --- | --- | --- | --- |
| YBL075C | SSA3 | 467 | 1950 | 181 |
| YJR004C | SAG1 | 1126 | 1953 | 332 |
| YKR050W | TRK2 | 1261 | 2670 | 268 |
| YDR242W | AMD2 | 408 | 1650 | 242 |
| YPL258C | THI21 | 1450 | 1656 | 191 |
| YLR174W | IDP2 | 519 | 1239 | 197 |
| YMR096W | SNZ1 | 450 | 894 | 275 |
| YHR211W | FLO5 | 2754 | 3228 | 185 |
| YCR107W | AAD3 | 878 | 1092 | 365 |
| YCR089W | FIG2 | 614 | 4830 | 694 |
| YAR050W | FLO1 | 3185 | 4614 | 162 |
| YDR216W | ADR1 | 958 | 3972 | 407 |
| YKR097W | PCK1 | 771 | 1650 | 157 |

**Table 6:**
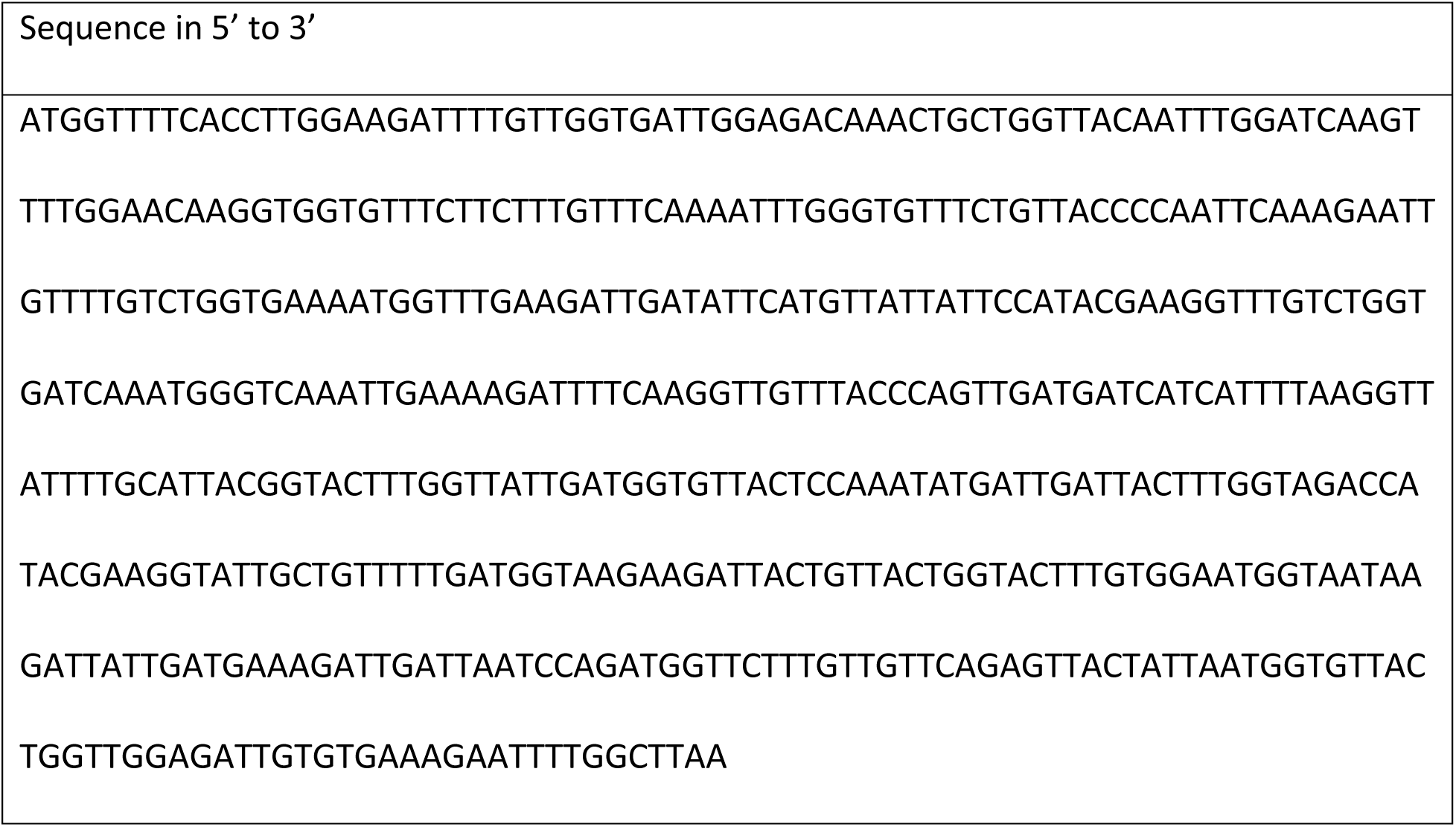
Codon-optimized nanoLuc for yeast (516 nts)

## Notes

### Competing Interest Statement

The authors have declared no competing interest.

